# Detecting and Quantitating Low Fraction DNA Variants with Low-Depth Sequencing

**DOI:** 10.1101/2020.04.26.061747

**Authors:** Ping Song, Sherry X. Chen, Yan Helen Yan, Alessandro Pinto, Lauren Y. Cheng, Peng Dai, Abhijit A. Patel, David Yu Zhang

## Abstract

DNA sequence variants with low allele frequencies below 1% are difficult to detect and quantitate by sequencing, due to the intrinsic error of sequencing-by-synthesis (NGS). Unique molecular identifier barcodes can in principle help NGS detect mutations down to 0.1% variant allele frequency (VAF), but require extremely high sequencing depths of over 25,000x, rendering high sensitivity mutation detection out of reach for most research and clinical samples. Here, we present the multiplex blocker displacement amplification (mBDA) method to selectively enrich DNA variants by an average of 300-fold in highly multiplexed NGS settings. On a 80-plex human single nucleotide polymorphism panel, mBDA achieves a 0.019% VAF limit of detection for single nucleotide variants, using only 250x sequencing depth, and detects human cell line contamination down to 0.07%. Using this technology, we constructed a 16-plex melanoma NGS panel covering 145 actionable mutations across 9 genes, and applied it to 19 fresh/frozen tumor biopsy tissue samples with high tumor fractions. We found low VAF mutations (0.2% to 5%) in 37% of the samples (7/19, 95% confidence interval 19%-58%). These results suggest that tumor heterogeneity could be significantly more pervasive than previously recognized, and can contribute significantly to acquired drug resistance to targeted therapies. We also validate mBDA panels on clinical cell-free DNA samples from lung cancer patients.

---

Sequencing-by-synthesis, also known as next-generation sequencing (NGS), is one of the most powerful methods available today for highly multiplexed characterization of DNA at many genetic loci [1, 2]. NGS is used routinely to detect DNA sequence variants with allele frequency of ≥ 5%. However, NGS struggles to report single nucleotide variants (SNVs) with variant allele frequencies (VAFs) of below 1% because all current NGS platforms have an average intrinsic error of at least 0.2% [3, 4]. Error-correction methods based on unique molecular identifiers (UMIs) have been proposed and can circumvent the NGS intrinsic error limitation to achieve SNV limits of detection of about 0.1% [5–7], but these generally require sequencing to extremely high depths of 25,000x or higher. This causes the per-sample NGS cost of even moderate-size panels to be over $1,000 (e.g. the 70-gene Guardant 360 panel [8, 9]), rendering sensitive detection of SNVs with VAF of below 1% unaffordable for many researchers, clinicians, and patients.

Simultaneously, there is growing research and clinical demand for high sensitivity detection of low VAF DNA sequence variants. Examples of low VAF detection need include cell-free DNA profiling for non-invasive cancer therapy guidance and post-treatment monitoring [10, 11], rare microbe detection for microbiome profiling [12, 13], and persistent bacterial subpopulations contributing to antibiotic resistence [14, 15] Digital PCR [16, 17] and allele-specific PCR [18] can be used to detect and quantitate one or a few suspected DNA variants with VAF down to 0.1%, but cannot reasonably scale to fulfill the multiplexing needs of many research and clinical applications.

Here, we present multiplex blocker displacement amplification (mBDA), a library preparation method that allows for robust NGS detection and quantitation of SNVs with low VAF down to 0.03% with only 250x sequencing depth. mBDA functions by selectively enriching DNA sequence variants during a multiplex PCR target enrichment step. Unlike other allele enrichment methods [20], mBDA can scale well to multiplex panels; we demonstrated 300-fold median enrichment in an 80-plex panel. Because the post-mBDA NGS library exhibits VAF far exceeding the NGS intrinsic error rate, low-depth sequencing is sufficient for variant detection and no error correction is needed. Furthermore, because the fold-enrichment of different variants are conserved across libraries and samples, it is possible to accurately quantitate the initial sample variants, with 95% accuracy within a factor of 2. By reducing the number of NGS reads needed by over 100-fold, mBDA uniquely enables affordable sequencing of DNA variants with very low VAF on lower-throughput NGS instruments such as the Illumina MiSeq.

## Results

The mBDA method allows highly multiplex sequence-selective PCR amplification of SNP alleles through the use of rationally designed Blocker oligonucleotides that perfectly bind to the intended SNP allele [21]. The Blocker binding region overlaps with that of the forward PCR primer, resulting in a competitive hybridization reaction in which the forward primer must displace the Blocker in order to bind to the DNA template (Fig. 1a). The Blocker sequence is designed to bind more strongly than the forward primer to the DNA templates bearing the intended SNP allele, but less strongly to DNA templates bearing the variant SNP allele. Consequently, variant templates are amplified with significantly higher yield per PCR cycle than wildtype templates. Through the course of many PCR cycles, the variant allele can be preferentially amplified with 1000-fold higher efficiency than the wildtype allele.

**FIG. 1:**
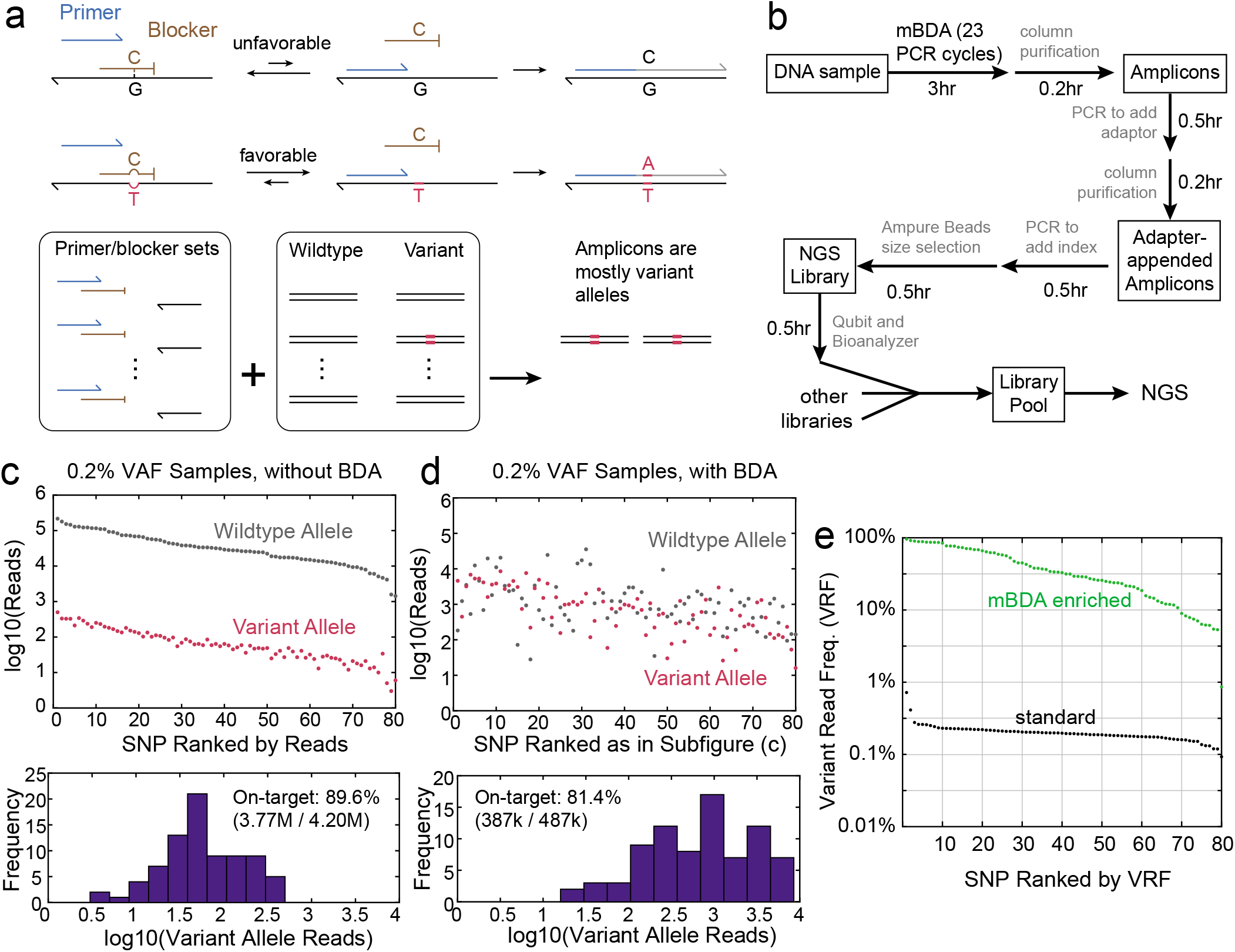
Allele enrichment with multiplex blocker displacement amplification (mBDA) enables rare variant detection with low-depth sequencing. **(a)** In mBDA, rationally designed Blocker oligonucleotides compete with forward PCR primers in binding to DNA templates at loci of interest. The Blocker is designed to be perfectly complementary to the wildtype DNA sequence. Any DNA sequence variant in the ≈20 nt enrichment region results in mismatched Blocker binding, leading to preferential PCR amplification of the variant. In highly multiplexed settings, a DNA sample may possess sequence variants in only a small fraction of the tested loci, resulting in amplicons being dominated by a small number of amplicons bearing the variant sequences. **(b)** The entire mBDA NGS library preparation workflow takes less than 6 hours. **(c)** Summary of NGS results on a library constructed using standard multiplex PCR target enrichment. Here, we used a 300 ng sample of a 99.8%:0.2% mixture of the NA18537 and NA18562 cell line genomic DNA. The panel targeted 80 loci on the human genome bearing single nucleotide polymorphisms (SNPs) in which NA18537 and NA18562 were homozygous for different alleles. Consequently, the VAFs of the NA18562-specific alleles were expected to be ≈0.2% for all 80 amplicons. **(d)** Summary of NGS results on a library constructed using mBDA on 300 ng of 99.8%:0.2% NA18537:NA18562. Compared to the library in panel (c), 9-fold fewer NGS reads were used, but the all variant alleles were sequenced to higher depth. **(e)** Summary of the variant read fraction (VRF) for each SNP locus in the libraries described in panels (c) and (d). The standard multiplex PCR NGS library showed roughly 0.2% median VRF, consistent with expectations.

The mBDA NGS library preparation workflow takes less than 6 hours from DNA to library; the process is summarized in Fig. 1b. To demonstrate allele enrichment with mBDA, we constructed an 80-plex mBDA NGS panel targeting 80 common human single nucleotide polymorphisms (SNPs). These 80 SNPs were selected such that the NA18537 and NA18562 human genomic DNA (gDNA) samples were homozygous for different alleles [22, 23]. A 99.8%:0.2% mixture of NA18537 and NA18562 gDNA was thus an easily-formulated reference sample with 0.2% VAF in all 80 SNP loci, when considering the NA18537 alleles as wildtype. Fig. 1c shows Illumina MiSeq sequencing results on the 0.2% VAF sample using a standard multiplex PCR target enrichment workflow. The number of reads aligned to the variant allele for each SNP locus is roughly 500-fold lower than the intended allele, consistent with expectations. Note that not all of these variants can be confidently called from this data; we will discuss VAF limit of detection (LoD) in the next section.

For comparison, the mBDA NGS results on a different aliquot of the 99.8%:0.2% NA18537/NA18562 sample are shown in Fig. 1d. Despite using 9-fold fewer total reads than the standard amplicon NGS library, the mBDA NGS library exhibits higher variant allele read depth because the number of intended allele reads are dramatically reduced. Fig. 1e displays the variant read fraction (VRF) for each SNP locus, with the median VRF increased from 0.2% in the standard NGS library to about 30% in the mBDA NGS library. Thus, the variant SNP alleles here are enriched by a median of 150-fold.

See Supplementary Section S1 for SNP loci and mBDA design details, and Supplementary Section S2 for NGS protocol optimization experiments. We use a custom bioinformatics pipeline to filter the NGS reads from FASTQ files, make variant calls, and determine corresponding VRF values (Supplementary Section S3). Standard bioinformatics tools such as Picard and GATK are optimized for hybrid-capture NGS panels, and we and others have observed these software to exhibit both false positive and false negative errors when making variant calls in amplicon sequencing libraries.

### VAF quantitation from mBDA NGS results

The relationship between the initial VAF in the sample and the observed VRF in the library can be described mathematically, based on the variant allele enrichment fold (EF).

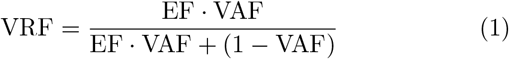

Note that VRF is not linear with EF; VRF values close to 100% imply much greater values of EF. For example, assuming VAF = 0.1%, VRF ≈ 10% when EF = 100; VRF ≈ 50% when EF = 1,000; and VRF ≈ 90% when EF = 10,000 (Fig. 2a).

**FIG. 2:**
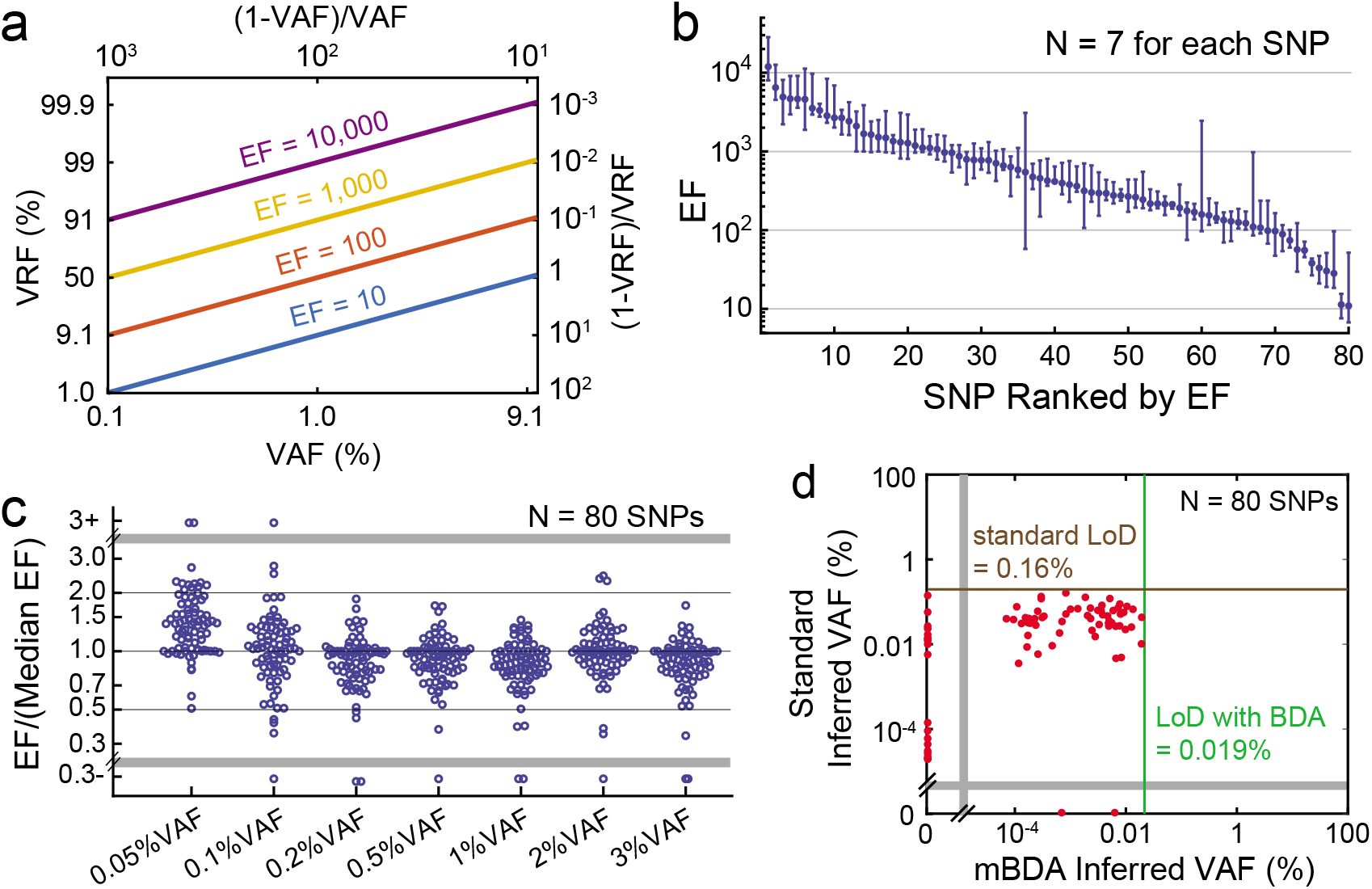
Quantitating variant VAFs based on observed VRF values from mBDA libraries. **(a)** Theoretical relationship between VAF and VRF for different allele enrichment-fold (EF). The relationship between 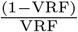 and 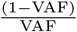 is expected to be linear, with a slope of 1 and an intercept of log10(EF). EF values are expected to vary for different SNPs, but are conserved across different experiments and VAFs for the same SNP allele. **(b)** Summary of inferred EF for each of the 80 variant SNP alleles, based on a set of 7 calibration mBDA NGS libraries using 0.05%, 0.1%, 0.2%, 0.5%, 1%, 2%, and 3% VAF samples. Dots show median values and error bars show maximum and minimum EF values. The 0.05% VAF sample used 200 ng of genomic DNA input; all other samples used 50 ng of genomic DNA input. **(c)** Relative EF values for different VAF inputs. The EF/(median EF) ratio can also be interpreted as the quantitation error for the calibration samples with known VAFs. For sample VAFs ≥ 0.1%, more than 95% of inferred VAFs are quantitated accurately to within a factor of 2. **(d)** VAF limit of detection (LoD) for standard amplicon NGS vs. mBDA NGS, based on triplicate NGS libraries for each method. Here, we define the LoD as the maximum inferred VAF from a pure wildtype (0% VAF) sample; the red dots show the inferred VAFs for each of the 80 SNPs. For mBDA NGS, the VAF is calculated based on the median EF from the 7 EF values summarized in panels (c) and (d). For standard amplicon NGS, the VAF is calculated as the VRF. BDA improves the VAF LoD by more than 8-fold vs. standard amplicon NGS.

The values of EF will vary based on the sequence identity of the variant allele and the intended allele, as well as neighboring context sequence. To calculate the EF for each of the 80 SNP loci, we ran a series of different NGS libraries using samples with known VAF (mixing NA18537 and NA18562). Rearranging equation (1) from above, we obtain:

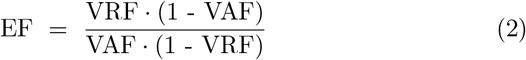

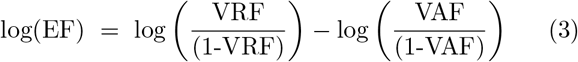

The transformed relationship between EF, VAF, and VRF is plotted in Fig. 2a. Fig. 2b shows the maximum, minimum, and median EF values calculated for each SNP locus, from 7 different mBDA NGS libraries with VAFs between 0.05% and 3%. Among the 80 SNP loci, 100% median EF values were greater than 10, 96% median EF values were greater than 30, and 85% of median EF values were greater than 100. See Supplementary Section S4 for EF calibration experiments.

The variation in EF values means that different limits of detection are achievable for different mutations. Given the standard NGS LoD of 2% VAF, an EF value of 10 corresponds an LoD of about 0.2% VAF, and an EF value of 30 corresponds to an LoD of down to 0.067% VAF. Thus, for the great majority of variants, the LoD for mBDA is not bottlenecked by the varying EF values, but rather by sample input quantity and/or DNA damage.

Fig. 2c shows the relative values of EF at each VAF compared to the median EF values. The median values of EF/(median EF) are close to 1 for mutations at VAF ≥0.1%, indicating that there is limited systematic bias in VAF quantitation. The systematic upward bias of 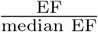 at 0.05% VAF may reflect the false positives due to DNA polymerase misincorporation errors.

To determine the limit of detection(LoD) of both standard amplicon NGS and mBDA NGS, we ran triplicate NGS libraries on pure NA18537 DNA samples that were nominally 0% VAF for all 80 SNP loci. The VAF LoD was determined to be 0.16% for standard amplicon NGS, based on the maximum observed VRF (Y-axis in Fig. 2d). For mBDA NGS, the VAF LoD was determined to be 0.019%, based on maximum VAF values calculated from VRF values and median EF values (X-axis in Fig. 2d). Thus, the mBDA NGS method improved the VAF LoD by roughly 8-fold compared with standard amplicon NGS because it eliminated the false positive variant calls from NGS intrinsic error. The remaining false positive errors were suspected to be primarily due to the misincorporation of incorrect nucleotides by the DNA polymerase at early cycles of PCR [24].

### Low VAF variant calls from mBDA NGS results

We next applied the mBDA NGS method to DNA samples with low fractions (0.07% to 0.25%) of conspecific contaminants (Fig. 3a). Because of the significant reduction in NGS reads required by mBDA NGS libraries, we were able to run 21 different DNA samples on a single Illumina MiSeq chip, while maintaining sensitivity to very low VAFs. In each of the libraries, the inferred VAFs for heterozygous alleles (green shaded region) and homozygous variant alleles (red) are generally consistent with expectations, with 80+% of VAFs within a factor of 2 of the median for each group.

**FIG. 3:**
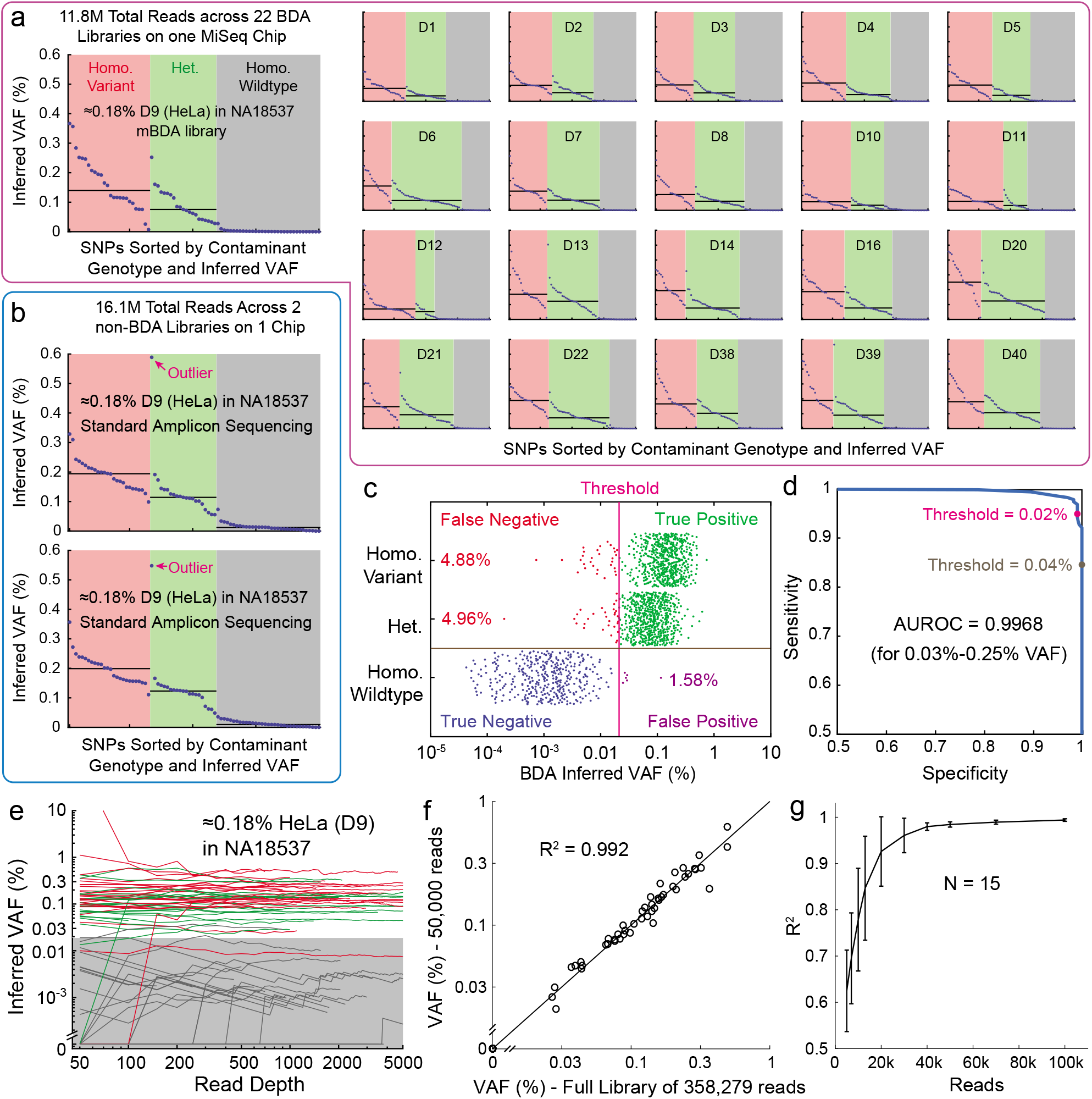
Detection and quantitation of low VAF variants using mBDA NGS. **(a)** Results for 21 mBDA libraries on a single MiSeq chip. Each library was generated from 300 ng of NA18537 contaminated with 0.07% to 0.25% of a different human DNA sample. Consequently, homozygous variant alleles had VAF between 0.07% and 0.25%, and heterozygous variant alleles had VAF between 0.03% and 0.13%. Individual library sizes ranged from 394k to 599k reads, and the library on-target rates varied from 75.2% to 83.8%. Each subfigure shows the inferred VAF for all 80 SNPs, sorted by contaminant genotype and then by inferred VAF; all subfigure Y axes ranged between 0% and 0.6% inferred VAF. The black horizontal lines show the median inferred VAF for homozygous variants and heterozygous variants. **(b)** Comparison libraries using standard amplicon NGS on the NA18537 samples contaminated with ≈ 0.18% D9 (HeLa). As in panel (a), each library used 300 ng of input DNA. There was significantly higher inferred VAF for homozygous wildtype (NA18537) alleles, and one outlier heterozygous SNP was called to be roughly 0.57% VAF. **(c)** Summary of variant call accuracy using the 0.019% VAF LoD threshold described in Fig. 4e. All inferred VAFs from panel (a) are displayed in this beeswarm plot. There was a false positive variant call rate of 1.58%, and a false negative rate of 4.88% or 4.96%, depending on whether the variant allele was homozygous or heterozygous, respectively. **(d)** Receiver operator characteristic (ROC) plot for variant calls using the data in panel (c). Setting the variant call threshold at 0.04% VAF would increase specificity to 100%, at the cost of reducing sensitivity to roughly 85%. The area under the ROC curve was 0.9968 for this set of samples, and likely would be higher for samples with greater contaminant fractions. **(e)** Inferred VAF as a function of read depth per SNP allele. Traces are colored by the contaminant SNP genotype (red for homozygous variant, green for heterozygous, and gray for homozygous wildtype). VAFs converge to their final values at about 250x read depth; shaded in gray are VAFs below our 0.19% LoD. **(f)** A random sample of 50,000 reads out of the mBDA 0.2% HeLa contaminant library (358k reads total) gave essentially the same inferred VAFs as the full library. **(g)** Analysis of reads needed for accurate VAF quantitation using mBDA libraries. Each point shows the mean *R^2^* values for the inferred VAFs of a read sample vs. the full library at different read sample sizes (N = 15 for each read sample size); error bars show 1 standard deviation. At ≥ 40, 000 reads, the R^2^ value reliably converged to above 0.99.

By contrast, with standard amplicon NGS, only 2 NGS libraries could be run on a MiSeq chip (Fig. 3b), and roughly 20% of the homozygous intended alleles had inferred VAFs of greater than 0.02%. Comparing the results from mBDA NGS and amplicon NGS on the same sample (roughly 0.18% HeLa in NA18537), we saw that the standard NGS method had somewhat tighter distributions of called VAFs than mBDA NGS. However, there was one outlier heterozygous allele with an expected VAF of 0.09% which produced a variant call of 0.58% VAF.

We could calculate the false positive and false negative rate of variant allele calls for the 1760 mBDA NGS sample/locus combinations, using the VAF LoD of 0.019% established in Fig. 3d. The overall specificity of variant calls was 98.42%, and the sensitivity was 95.12% for homozygous variant alleles and 95.04% for heterozygous variant alleles (Fig. 3c). Changing the VAF LoD threshold would change the relative tradeoff between sensitivity and specificity (Fig. 3d); the area under the receiver operator characteristic (AUROC) was calculated to be 0.9968. For comparison, the standard amplicon NGS libraries yield a specificity of 100% and a sensitivity of 69.23% for homozygous variant alleles, and a sensitivity of 9.52% for heterozygous variant alleles, based on the previously established VAF LoD of 0.19%.

Because mBDA significantly enriches variant alleles, low depth sequencing is sufficient for detecting and quantitating variants with low VAF. Fig. 3e shows the inferred VAF for each of the 80 SNP loci based on the read depth of each SNP. The inferred VAFs above the 0.019% LoD quickly converge to their final values, typically within 250 reads. Thus, in theory, an NGS library as small as 250·80 = 20,000 reads would be sufficient to achieve the detection of 0.1% VAF. In practice, imperfect amplicon depth uniformity across the 80 loci means that roughly 50,000 reads are needed to obtain accurate inferred VAF values (Fig. 3fg; see also Supplementary Section S5).

The experiments shown in Fig. 3 used 300 ng of input DNA to minimize the effects of Poisson distribution during sample preparation, and does not represent a limitation of the mBDA NGS method. In Supplementary Section S6 we show experimental results on mBDA NGS on 10 ng of input DNA (0.18% HeLa contaminant in NA18537).

The mBDA VAF limit of detection we achieve matches or exceeds the most UMI-based NGS methods with 50-fold fewer reads required, but is worse than the consensus accuracy achieved by Pacific Biosciences sequencing of circularized templates. However, PacBio is an expensive platform that few labs have access to, and its expensive consumables (on a per-read basis) render it noncompetitive in the clinical sequencing space. For most clinical applications PacBio-level error rates (10^−5^ to 10^−7^) are unnecessary, due to biological sample limitations (e.g. 10 ng of DNA from fine needle aspirates). Additionally, the mutation VAF limits of detection for biological samples are frequently limited by DNA chemical damage, such as cytosine deaminations and guanine oxidations; PacBio sequencing will not be able to resolve these from true mutations.

### Contaminant identification based on NGS data

In the specific application to human cell line contamination, the mBDA NGS data can be used to inform the specific identity of the contaminant. In this setting, a database of known possible contaminant genotypes is constructed, and the mBDA NGS results are compared against genotypes of each potential contaminant. For each database genotype *j*, we can compute the likelihood L_*j*_ of it generating the observed mBDA NGS libraries based on the following formula:

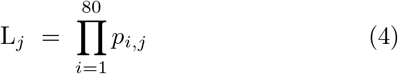

where *p_i,j_* is the probability of generating the observed variant call at locus *i* based on a true variant state *j*. In this framework, we consider only the presence or absence of a variant, rather than the VAF quantity, so *p_i,j_* is equal to the True Positive rate when the inferred VAF is above the 0.019% VAF LoD and comparison genotype *i* is either homozygous variant or heterozygous. Thus, equation (4) above can be rewritten as:

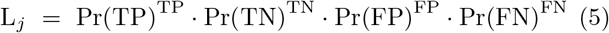

where Pr(TP) is the probability of a True Positive based on Fig. 3c, and TP is the number of True Positives for genotype *j*, etc. Fig. 4a shows graphical examples of the likelihood computation for two different genotypes; the correct comparison genotype has a much higher likelihood L than the incorrect genotype.

**FIG. 4:**
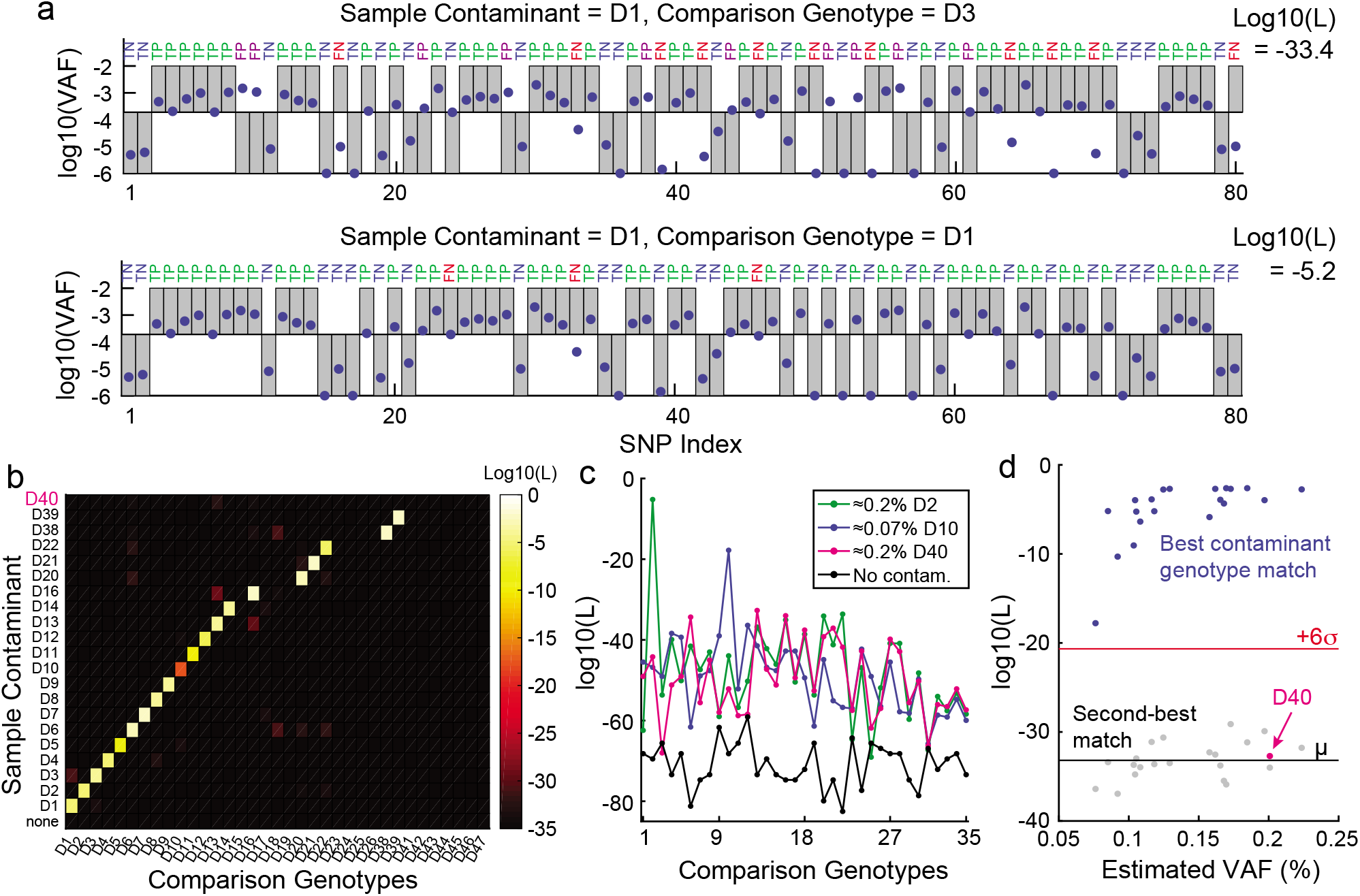
Determination of contaminant identity based on mBDA NGS data. **(a)** Calculation of contaminant likelihood. For each SNP, we decided a positive or negative variant allele call based on whether the inferred VAF (blue dots) was greater than the threshold of 0.019%. For each potential contaminant, based on whether a variant allele existed for each SNP (gray boxes), we determined whether our mBDA-inferred variant would be a True Positive (TP), True Negative (TN), False Positive (FP), or False Negative (FN) relative to the potential contaminant. The overall likelihood L for the potential contaminant was calculated as the product of the likelihoods of all SNPs, with the probabilities of TP, TN, FP, and FN set as 95.08%, 98.42%, 1.58%, and 4.92%, respectively, based on Fig. 2d. The correct contaminant was expected to have a much higher value of L than incorrect contaminants. **(b)** Heatmap plot of log10(L) values for all pairwise combinations of all 21 tested samples (with contamination fraction between 0.07% and 0.22%) vs. 35 database genotypes. **(c)** Representative distributions of log10(L) for 4 different samples. D2 shows a typical sample whose contaminant genotype exists within the database. D10 shows the sample with the weakest maximum log10(L) value whose contaminant’s genotype exists in the database. D40 shows a sample whose contaminant’s genotype is not included in the database. “No contam.” shows a sample of pure NA18537. **(d)** Plot of highest and second-highest log10(L) values against the contamination fraction. The value of log10(L) decreases for lower contamination fractions, consistent with expectations. However, even the D10 sample with 0.07% contamination is confidently identified, at over 6 standard deviations above the mean second-highest log10(L) value. The D40 sample, whose genotype was intentionally omitted from the database, can also be confidently identified as an unknown contaminant based on its highest log10(L) value.

Fig. 4b displays the logarithm of L for every pairwise combination of the 21 mBDA NGS libraries in Fig. 3a and 35 different comparison genotypes. The genotype of contaminant D40 was intentionally withheld from the database, in order to observe the effects of an unknown contaminant. In all cases other than D40, the correct genotype yielded the highest L value. In the case of contamination by D40, no comparison genotype produced very high L, but all L values were significantly higher than the uncontaminated DNA sample (Fig. 4c). The L value of correct contaminant genotype decreased for samples with very low contaminant fractions (Fig. 4d); the D10 contaminant sample had the lowest contaminant fraction (0.07%) and corresponded to the lowest value of L for the correct genotype. However, based on the distribution of second-highest values of L, even D10 could be confidently called with 6 standard deviations of confidence (*p* < 10^−9^).

### Cell Line Contamination Screening with qPCR

Quantitative polymerase chain reaction (qPCR) is by far the most commonly used method for detecting DNA markers due to its reliability, ease-of-use, and short turnaround time. qPCR can be effectively used to detect unique DNA sequences of a known contaminant. For example, qPCR-based detection of mycoplasma contamination of human cell lines is well-established as a method for detecting inter-species contamination [25]. However, there are many potential human cell line contaminants. Studies report that the top 10 most common contaminants collectively only account for about 50% of all known contamination cases [26]. To our knowledge, no qPCR methods have been reported allowing for detection of arbitrary conspecific DNA contamination.

Here, we show that mBDA is compatible with qPCR, and can be used to build rapid and easy-to-use assays for detecting arbitrary conspecific cell or DNA contamination. In our mBDA qPCR implementation, we design a mBDA primer/blocker set that specifically suppresses a set of SNP alleles on which the desired cell line is homozygous. When the set of SNPs selected is sufficiently large, any contaminant DNA’s genotype is likely to differ from the desired cell line in at least one of the panel’s SNPs. A double-stranded intercalating dye such as SybrGreen can be used to report the total quantity of PCR amplicons. In an ideal mBDA qPCR implementation with infinite EF, an uncontaminated sample of cell line DNA would give no amplification, and even an infinitesimal contamination could be detected and quantitated via amplification cycle threshold (Ct).

The closed-tube nature of qPCR reactions, while minimizing contamination risk, precludes solid-phase reversible immobilization (SPRI) bead-based size selection steps that mitigate primer dimers and non-specific genomic amplification. Thus, we found that the full 80-plex panel developed for NGS exhibited significant primer dimers that generated a small qPCR Ct value even in the absence of DNA templates. We found that reducing the mBDA panel size to 40-plex significantly improved our qPCR LoD for detecting contaminants, and reducing to 21-plex resulted in an additional marginal improvement (Supplementary Section S7).

Fig. 5ab shows qPCR detection of DNA samples with various fractions of HeLa contaminant in NA18537, using a 21-plex SNP assay. HeLa was selected as the contaminant because it is the single most frequent source of human cell and DNA contamination, accounting for roughly 25% of all reported cases [26]. Out of the 21 SNP loci in the qPCR assay with NA18537 as the intended genotypes, HeLa had 17 variant alleles (Fig. 5a). The mean qPCR Ct values observed for uncontaminated NA18537 vs. NA18537 with 0.1% HeLa were dramatically statistically different, with a p-value of 1.8 · 10^−30^.

**FIG. 5:**
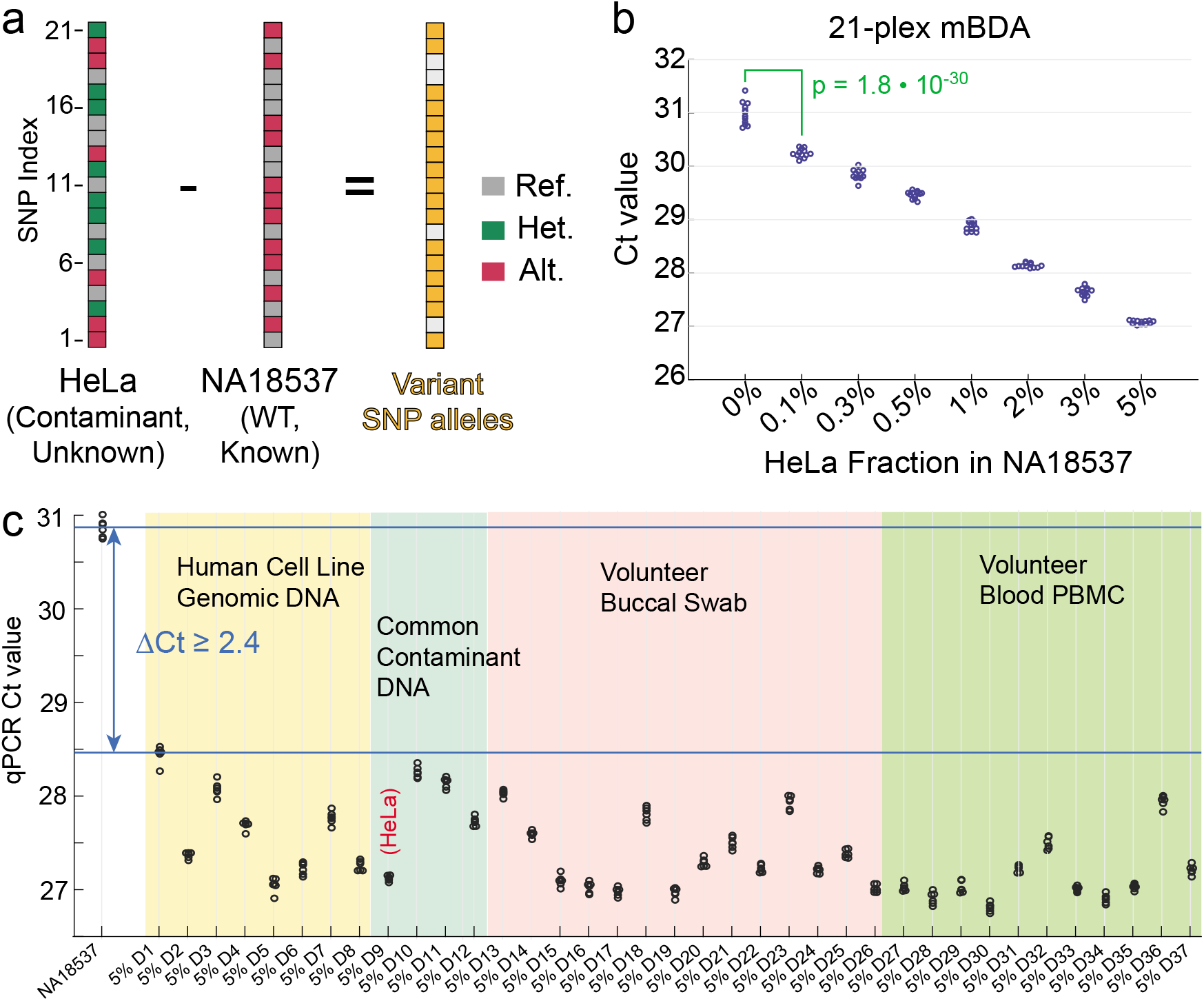
Detecting cell line contamination using mBDA with qPCR. **(a)** Given an intended human cell line (here NA18537), our mBDA qPCR assay will detect potential contamination by any other human cells. The unknown contaminant (here, HeLa) is likely to differ in genotype in at least one SNP from the intended cell line. Variant SNP alleles from the contaminant will be preferentially amplified, resulting in lower cycle threshold (Ct) values when using a double-stranded DNA intercalating dye such as SybrGreen I. Here, we selected a 21-plex subset of the prior 80-plex mBDA panel. **(b)** Experimental mBDA qPCR Ct values for 20 ng NA18537 with varying fractions of HeLa contamination. The beeswarm plot shows the observed Ct values for 12 independent reactions for each sample. Even the 0.1% HeLa contaminant could be confidently distinguished from the pure NA18537 gDNA sample. **(c)** Summary of qPCR results for detection of 5% contaminant in NA18537 with 37 different human DNA contaminants (6 replicates per sample, 20 ng DNA each). Because the different contaminants had different numbers and sets of variant alleles, there was variation in the Ct values of the contaminated samples. Based on the observed ΔCt values at 5% contaminant fraction, we extrapolated that 1% contaminant fraction should be detectable for all human DNA contaminants.

We expect that the 21-plex qPCR assay is able to detect arbitrary human mutations, due to the low probability that a contaminant would have identical genotype in the 21 SNP loci as the intended cell line. To experimentally support this hypothesis, we next ran the 21-plex mBDA qPCR assay on NA18537 with 5% contaminant fractions of 37 different human DNA samples (Fig. 2c). The contaminants included DNA from common cell lines as well as DNA from volunteers. All 5% contaminant samples could be confidently differentiated from the uncontaminated sample, and the large ΔCt difference suggested that all of the contaminants could be detected at 1% using qPCR.

Our simulations suggested that any arbitrary cell line would be homozygous in at least 21 of the 80 SNP loci that we built our mBDA NGS panel for (Supplementary Section S8). Of a 21-plex subset mBDA panel, our analysis and simulations suggested that there would be at least 8 variant alleles out of 21 SNP loci for an arbitrary human cell contaminant, based on the population frequencies of each SNP allele reported in the 1000 Genomes database [23]. To help readers who work with cell lines construct their own qPCR panels to detect contamination in their cell lines, we have provided mBDA designs for both alleles of 80 SNPs (Supplementary Excel dataset). We envision that the genotype of the intended cell line, if unavailable in public databases, can be quickly and inexpensively obtained through Sanger sequencing, a small microarray, or a small NGS library (without mBDA).

Although we expect that the primary interest of cell line researchers is to detect arbitrary contaminants in known cell lines, it is also possible to use mBDA qPCR assays to detect known contaminants without genotype information regarding the specific intended cell line. In this scenario, we selected SNP loci in which the alternate allele has extremely low frequency in the population, and the contaminant is heterozygous. As a demonstration, we designed a 3-plex mBDA assay for HeLa-specific SNPs, and experimentally showed that we could detect 1% HeLa contaminant in 8 different cell lines with unknown genotypes (Supplementary Section S9).

### Melanoma mBDA NGS Panel Reveals Frequent Heterogeneity in Tumor Tissue Samples

Most qPCR, Sanger, and NGS-based assays and panels for oncology targeted therapy selection [40–42] have mutation VAF sensitivities of between 1% and 5%, and thus are unable to identify low VAF drug resistance mutations arising from trace subclones due to tumor heterogeneity. However, under the selective pressures of targeted therapy, tumor subclones with drug resistance mutations can rapidly expand and cause treatment failure or cancer recurrence [43–46]. Thus, reliable detection of low VAF drug resistance mutations can inform personalized treatment selection, including the use of combination therapies, to improve patient outcomes.

To address this challenge, we next constructed a 16-plex mBDA NGS panel covering 145 commonly observed melanoma mutations across 9 genes (Fig. 6a). The BDA enrichment regions were designed to cover the most frequently observed mutations in the 9 genes, based on the COSMIC database [47]. Because this cancer panel covers a range of different mutations, we arbitrarily selected one of the two DNA strands to be the target strand for Blocker binding, unlike the case of SNP detection where we can intentionally target the strand with a larger mismatch thermodynamic penalty. To measure the VAF limit of detection and EF values for the cancer mutations, we ran calibration NGS experiments using human genomic DNA (NA18537) spiked in with varying quantities of synthetic gene blocks with length between 490 nt and 500 nt (see Supplementary Section S10 for reference sample preparation and EF calibration details). To analytically validate the accuracy of our mBDA NGS panel in detecting and quantitation low VAF mutations, we performed comparison experiments on spike-in samples against the Bio-Rad QX200 ddPCR (Supplementary Section S11).

**FIG. 6:**
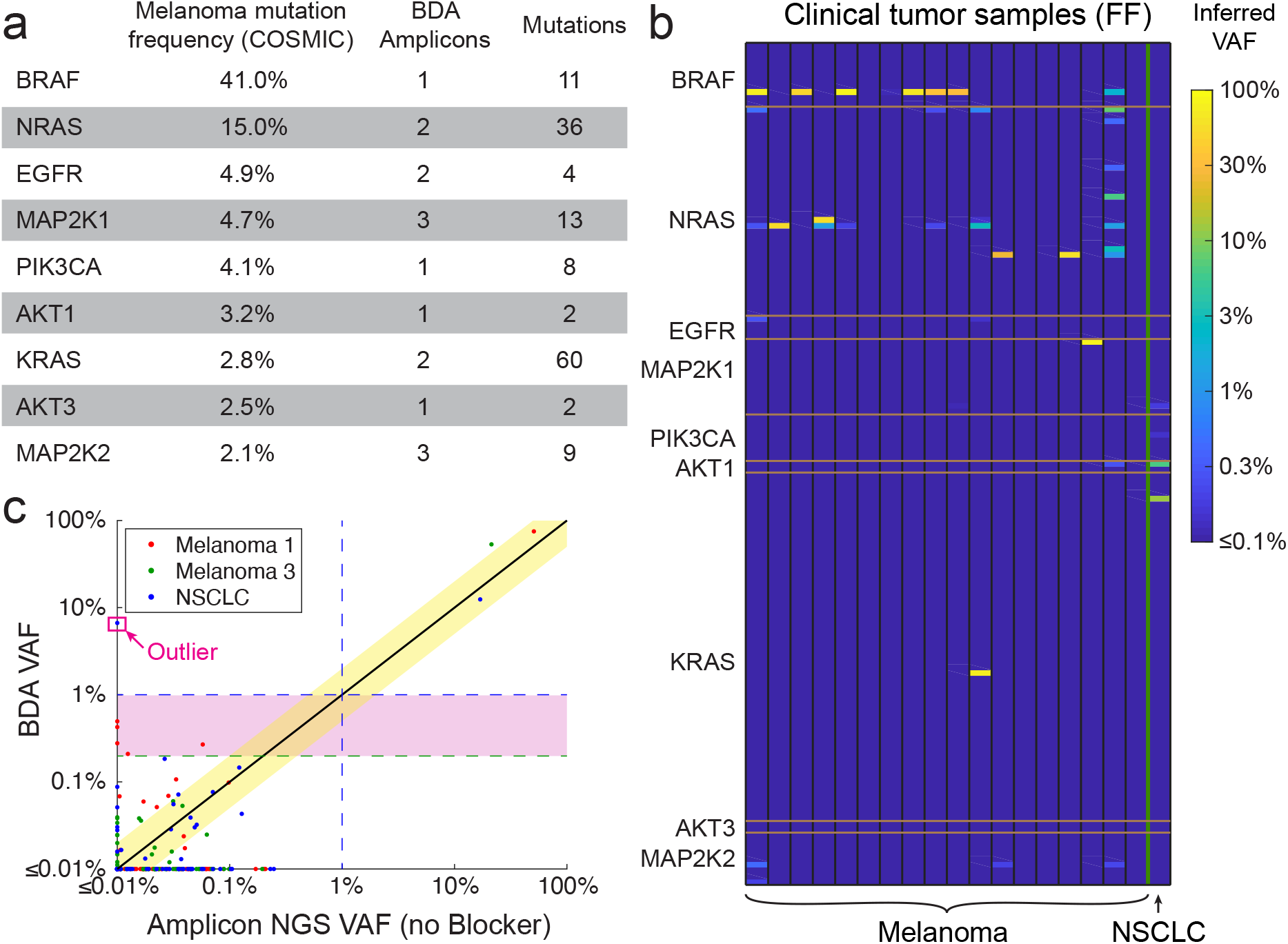
Melanoma mBDA NGS panel covering 9 genes and 145 mutations in the COSMIC database; the panel comprises a 16-plex mBDA reaction. **(a)** Panel contents. See Supplementary Section S10 for detailed list of mutations covered. **(b)** Inferred VAFs from the mBDA NGS panel for 18 melanoma tumor samples (fresh/frozen) and 1 non-small cell lung cancer (NSCLC) tumor sample. Samples were purchased de-identified from Origene. 50 ng of extracted DNA were used as input for each library. The number of NGS reads for each sample ranged between 45,844 and 89,718. **(c)** Comparison plot of inferred VAF from standard amplicon NGS. Shown in the pink region are 8 mutations at VAF between 0.1% and 1% identified by the mBDA panel. The boxed outlier corresponds to an AKT1 mutation in the NSCLC sample with extremely low sequencing depth in the amplicon NGS panel (1 mapped read). On the melanoma samples, the sequencing depth for the AKT1 amplicon was 3,058 and 4,788, indicating that this outlier may be specific to the NSCLC sample.

We applied the melanoma mBDA NGS panel to a total of 19 clinical fresh/frozen tumor samples purchased deidentified from a commercial supplier (Origene); of these 18 were melanoma tissue samples and 1 was a non-small-cell lung cancer tissue sample. All samples had a tumor fraction that was at least 75%, based on histological analysis. The called variants are summarized in Fig. 6b and Table 1. A total of 7 of the 19 samples (37%) had low VAF mutations between 0.2% and 5%. The 95% confidence interval of fresh/frozen tumor samples with low VAF mutations is 19% to 58%, based on binomial distribution analysis. To confirm our findings, we performed ddPCR comparison experiments for these fresh/frozen tissues as well as 4 FFPE tissue samples (Supplementary Section S11).

**TABLE 1:** Summary of mBDA NGS variant calls from FF tumor samples, using a VAF cutoff of 0.2% for all mutations. Melanoma tissue samples 2 and 14 yielded very low extracted DNA concentrations, and were not analyzed. Note that because mBDA VAF quantitation is only accurate to within a factor of 2, called variants with VAFs greater than 50% likely

| FF Sample | Age/Sex | Stage | Tumor % | Called Variants | FF Sample | Age/Sex | Stage | Tumor % | Called Variants |
| --- | --- | --- | --- | --- | --- | --- | --- | --- | --- |
| 1 | 42/M | IIIB | 80% | 75% BRAF p.V600E (c.1799T>A)<br>0.50% NRAS p.G12S (c.34G>A)<br>0.27% NRAS p.Q61K (c.181C>A)<br>0.28% EGFR p.G719S (c.2155G>A)<br>0.43% MAP2K2 p.C125S (c.373T>A)<br>0.21% MAP2K2 p.E207K (c.619G>A) | 2 | 81/F | N/A | N/A | (not analyzed) |
| 3 | 51/M | III | 98% | 53% NRAS p.Q61K (c.181C>A) | 7 | 56/F | III | 80% | None |
| 4 | 35/F | IV | 75% | 48% BRAF p.V600E (c.1799T>A) | 8 | 42/F | IV | 95% | None |
| 5 | 43/M | IIIB | 75% | 56% NRAS p.Q61K (c.180_181delinsTA)<br>1.2% NRAS p.Q61K(c.181C>A) | 14 | 42/F | N/A | N/A | (not analyzed) |
| 6 | 49/F | III | 80% | 90% BRAF p.V600E(c.1799T>A)<br>0.21% NRAS p.Q61K (c.181C>A) | 15 | 46/M | N/A | 90% | None |
| 9 | 57/M | IIC | 95% | 70% BRAF p.V600E (c.1799T>A) | 16 | 34/F | III | 90% | None |
| 10 | 46/F | III | 90% | 37% BRAF p.V600E (c.1799T>A)<br>0.26% NRAS p.G12S (c.34G>A)<br>0.22% NRAS p.Q61K (c.181C>A) | 20 | 50/M | IV | 90% | None |
| 11 | 41/M | IIIA | 90% | 31% BRAF p.V600E (c.1799T>A) |  |  |  |  |  |
| 12 | 66/M | IIIA | 85% | 0.72% NRAS p.G12S (c.34G>A)<br>3.0% NRAS p.Q61K (c.181C>A)<br>76% KRAS p.G13P (c.37_38inv) |  |  |  |  |  |
| 13 | 72/F | IIIB | 70% | 26% NRAS p.Q61L (c.182A>T)<br>0.20% MAP2K2 p.C125S (c.373T>A) |  |  |  |  |  |
| 17 | 66/F | IV | 95% | 60% NRAS p.Q61L (c.182A>T) |  |  |  |  |  |
| 18 | 64/F | IV | 75% | 81% MAP2K1 p.K57E (c.169A>G) |  |  |  |  |  |
| 19 | 55/? | IV | 90% | 2.3% BRAF p.V600E (c.1799T>A)<br>7.9% NRAS p.G12S (c.34G>A)<br>0.44% NRAS p.G12C (c.34G>T)<br>1.3% NRAS p.Q61K (c.181 C>A)<br>0.27% AKT1 p.Q79K (c.235C>A)<br>0.22% MAP2K2 p.C125S (c.373T>A) |  |  |  |  |  |
| NSCLC | 55/M | IIB | N/A | 12% KRAS p.G12D (c.35G>A)<br>6.7% MAP2K2 p.Q79K (c.235C>A) |  |  |  |  |  |

As part of our design and bioinformatics interpretation process, we design primers and blockers to avoid non-pathogenic SNPs with population allele frequencies greater than 1% using the 1000 Genomes and Kaviar databases. The high VAF oncogene mutations that we observed in the tumor samples, such as BRAF-V600E and NRAS-Q61K, are not reported to be present in the population at significant frequencies, presumably due to the strong selective pressure against individuals with these mutations. Consequently, we believe these are likely to be somatic tumor mutations, though we cannot be sure without matched normal tissue or blood samples.

Interestingly, all samples with low VAF subclonal mutations also had at least one high VAF mutation (Table 2). A chisquare analysis weakly suggests that the presence of low VAF mutations and high VAF mutations are not statistically independent (*p* = 0.046). In order words, the presence of low VAF subclonal drug resistance mutations appears to be higher in tumor samples with high VAF clonal mutations.

**TABLE 2:** Summary of distribution of samples with called high VAF mutations (≥ 5%) and low VAF mutations (between 0.2% and 5%). Chi square analysis of this data suggests that presence of high VAF mutations and low VAF mutations are not independent (*p* = 0.046). Significant tumor heterogeneity is observed more frequently in tumors with high VAF mutations.

|  | High VAF (5+%) Negative | High VAF (5+%) Positive | Total |
| --- | --- | --- | --- |
| Low VAF (0.2% to 5%) Negative | 5 | 7 | 12 |
| Low VAF (0.2% to 5%) Positive | 0 | 7 | 7 |
| Total | 5 | 14 | 19 |

### Validation on clinical cfDNA samples

Cell-free DNA (cfDNA) in peripheral blood represents a promising class of biomarkers for non-invasive tumor profiling, and can be useful in cancer management not only for therapy selection in cases where tumor biopsy is not convenient, but also longitudinally for post-treatment monitoring. Detection of actionable mutations in cfDNA is also well-suited for mBDA, since mutation VAFs in cfDNA can be quite low (<1%) because of the wildtype cfDNA derived from healthy dying cells as normal part of homeostasis.

Here, we validate the effectiveness of mBDA NGS panels on clinical cfDNA samples from Stage IV non-small cell lung cancer patients. The mBDA panel covers 31 hotspots in 14 genes (AKT1, ALK, BRAF, DDR2, EGFR, ERBB2, KRAS, MAP2K1, MET, NRAS, PIK3CA, PTEN, ROS1, and TP53), and was developed commercially by Nuprobe as the VarMap NSCLC panel. The called mutations and their VAF are compared against the VAF called by deep sequencing using UMIs (Fig. 7). Qualitative and quantitative concordant calls are made for mutations with VAF down to 0.23% in clinical samples (see Supplemental Excel document). Discordant calls were observed only for mutations below 1.3% VAF, and could be due to a combination of Poisson distribution for small numbers of molecules and biases in conversion yield for different approaches on different genetic loci.

**FIG. 7:**
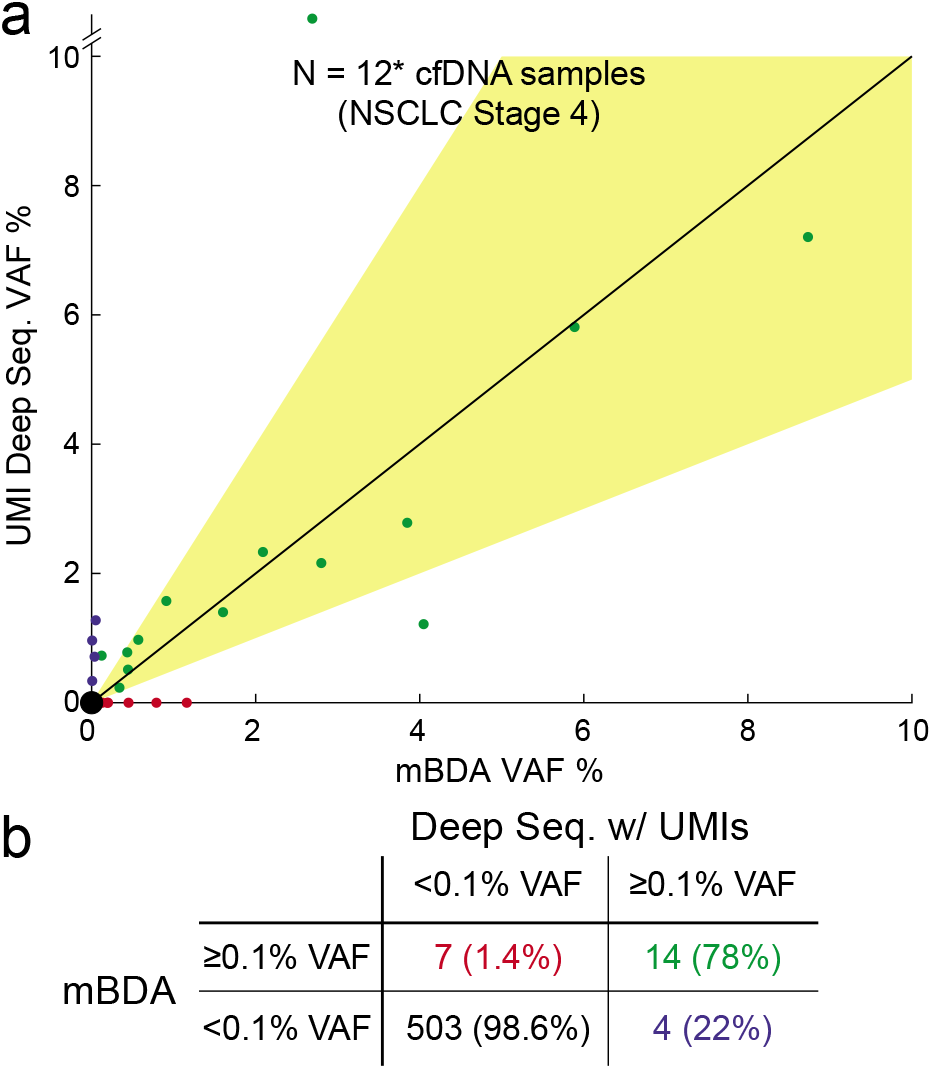
Validation on clinical cfDNA samples. An 14-gene, 31-amplicon mBDA panel was applied to 12 deidentified cfDNA samples from Stage 4 NSCLC patients. Input cfDNA quantities ranged between 6.0 ng and 18.9 ng. Simultaneously, aliquots of these cfDNA samples were also analyzed for tumor mutations through deep sequencing with UMIs following the method of Ref. [19]. One clinical cfDNA sample tested was excluded from this analysis because of strong suspicion of sample mislabeling (Sample R in Supplementary Excel Table). **(a)** Comparison of inferred VAFs between mBDA and deep sequencing with UMIs. The yellow shaded region indicates VAF agreement to within a factor of two. There is one EGFR exon 19 ELREA deletion outlier in which mBDA called 2.7% VAF and deep sequencing called 20.2% VAF. In another sample, the same EGFR exon 19 ELREA deletion was called as 8.7% VAF and 7.2% VAF by mBDA and deep sequencing, respectively. **(b)** Qualitative concordance of called tumor mutations. Across a total of 528 loci (12 samples * 44 loci), we observe a 78% positive concordance rate, and a 98.6% negative concordance rate.

## Discussion

The mBDA technology presented here significantly scales the multiplexing of BDA to 80-plex and demonstrates integration with NGS to allow broad detection and quantitation of rare DNA mutations with ≤0.1% VAF LoD. Although allele enrichment methods have been extensively researched (e.g. ICE-COLD PCR [20], Boreal Genomics [27], PNA/LNA blocker PCR [28], nuclease assisted minor-allele enrichment [29], CUT-PCR [30]), we are not aware of any other allele enrichment technologies that have been successfully scaled and integrated with NGS to allow low-depth sequencing analysis of low VAF mutations in over 10-plex settings. Generally speaking, other allele enrichment methods are sensitive to operational conditions (temperature, time, enzyme/DNA purity). This renders multiplexed panels challenging to design and optimize, because variant alleles from multiple loci may enrich optimally at different conditions.

In this work, we first demonstrated mBDA using an 80-plex amplicon panel that covered in total roughly 8,000 nt and performed allele enrichment on roughly 1,600 nt. Given our observed on-target rates of over 80%, we do not think that primer dimers and non-specific amplification are currently the bottleneck for scaling up mBDA. Primer dimers are typically much shorter than on-target amplicons, and nonspecific genomic amplification results in amplicons that are typically much longer than on-target amplicons. Consequently, the vast majority of primer dimers and nonspecific amplicon can and are removed through SPRI bead size selection. Commercial amplicon target enrichment NGS panels (e.g. Ampliseq) include up to 24,000-plex PCR primers, so we expect that scaling up mBDA to over 1000-plex in a single tube should be possible with some experimental optimization.

The mBDA technology, as currently designed, is primarily directed to the detection and quantitation of mutations and small insertions/deletions within hotspot regions. Thus, mBDA is well-suited to detection of actionable cancer mutations in clinical guidelines such as by NCCN, as well as to personalized mutations found from initial whole exome sequencing or whole genome sequencing analysis of tumor biopsies [31]. For standard single-base replacement mutations, we are able to achieve mBDA success rates of well over 95%, with failures primarily due to extremely high G/C content, or context sequences with highly repetitive DNA sequences that cause PCR mis-priming. We observed slightly worse performance in enriching single-base insertions and deletions from homopolymeric repeats of over 6 nt, but these are sequences that all PCR and NGS struggle with.

mBDA is less suitable for discovery of new mutations across the entire exon regions of many genes, and is unsuitable for discovery of new structural variations (e.g. chromosomal translocations). To comprehensively detect mutations in tumor suppressor genes such as BRCA1/BRCA2, we would need to pursue a split tube strategy wherein the exons are tiled by different amplicons split across multiple tubes. Tube splitting is needed for most amplicon sequencing approaches for comprehensive exon coverage, because mutations in regions covered by primers cannot be detected in amplicon sequencing; different primers in tube 2 are needed to detect potential mutations in loci covered by primers in tube 1.

For SNVs with VAFs between 0.03% and 3%, we showed that results from mBDA NGS could accurately quantitate sample VAF through a mathematic transformation assuming conserved enrichment fold (EF) values, with up to 95% accuracy within a factor of 2. For low VAF somatic mutations, quantitation accuracy is frequently limited by biological variability. Even adjacent tissue sections can have different VAFs, and in cfDNA a wide range of biological factors (e.g. exercise, time of day, and bacterial/viral infection) all affect tumor mutation VAF. However, mBDA suffers from less accurate VAF quantitation when initial VAF and/or EF is very high. For example, a 10% VAF sample with EF = 10,000 would result in a mBDA library VRF of 99.9%, which cannot be accurately quantitated for the same reason that 0.1% VRF cannot be accurately quantitated. When accurate quantitation of high VAF mutations is needed, we recommend constructing a standard amplicon NGS library with low sequencing depth (e.g. 250x) in addition to the mBDA library. The mBDA library will accurately quantitate low VAFs between 0.03% and 3%, and the standard amplicon library will accurately quantitate VAFs between 3% and 97%.

We demonstrated mBDA using a panel of non-pathogenic human SNPs, and showed that it could be used for the detection of human cell line contamination in both NGS and qPCR settings. Cell lines serve as critical models for biomedical research, allowing the use of effective and inexpensive experiments to understand cell and organism function as well as to establish feasibility for potential therapeutics [32–34]. However, contamination and misidentification of cell lines have become a widespread problem that limit the reproducibility of experimental results and threaten the validity of published conclusions [35–37]. Recent estimates suggest that up to 35% of cell lines in use today suffer from contamination, and over 30,000 research publications’ findings may be compromised [38]. With our mBDA qPCR assays, researchers will be able to detect low-level contamination by conspecific DNA down to less than 1%, increasing the rigor of scientific research based on cell lines.

As a proof-of-concept for clinical applications of mBDA-based NGS panels, we constructed a 16-plex melanoma mBDA NGS panel and applied it to 19 tumor tissue samples. We found that 37% of the samples (95% CI: 19% - 58%) contained detectable subclonal mutations, including known drug resistance mutations, within our panel at between 0.2% and 5% VAF. Furthermore, we expect that the fraction of patient tumor samples with low VAF mutations would be higher given a broader panel covering more genes and loci. The rise of drug resistance mutations in patients undergoing targeted therapies has generally been considered as a combination of (1) rise of *de novo* mutation(s) during treatment, and (2) expansion of pre-existing subclones with the resistance mutation(s). Our results here statistically indicate that the latter mechanism may be applicable to a significant fraction of patients. Alternatively, reports of genomic heterogeneity of in the skin of healthy-ageing individuals [39] suggest that the correlation between low VAF mutations and high VAF mutations may be due to more complex mechanisms not yet fully understood.

In this work, we applied the mBDA NGS panel to fresh/frozen (FF) tumor tissue, rather than formalin-fixed paraffin-embedded (FFPE) tissue, because of well-documented deamination and oxidation damage FFPE DNA [48, 49]. With FF tissue samples, we can be more confident that the low VAF mutations called are truly indicative of subclonal mutations due to tumor heterogeneity, rather than artifacts from DNA damage.

In our preliminary experiments applying mBDA panels to melanoma FFPE samples, we observed systematic cytosine deamination damage resulting in C>T and G>A false positives at up to 0.6% VAF. Because we typically set our LoD reporting cutoff at double the highest observed false positive VAF, this means that for FFPE samples, we cannot report C>T and G>A variants at below 1.2% VAF. Other mutations, such as the BRAF-V600E T>A mutation, can still be reported at down to 0.1% VAF. However, because the extent of FFPE damage is highly dependent on the age and the storage/handling of the sample, it is not generalizable to determine VAF LoDs across different FFPE samples. Thus, for FFPE samples, we expect to determine the LoD for different types of mutations <u>within the same sample</u>, by designing BDA to other portions of the genome that are not expected to be mutated (e.g. housekeeping genes such as GAPDH).

To date, all NGS panels that achieve 0.1% VAF LoD use unique molecular identifiers (UMIs) with ultradeep sequencing of more than 25,000x depth. This renders rare mutation profiling only feasible on the highest throughput NGS instruments such as the NovaSeq (Table 3). However, the high capital expense (roughly $1M for a NovaSeq) and the large number of samples that need to be pooled to obtain economies of scale render high-throughput NGS instruments out of reach for most hospitals and reference laboratories. By enabling accurate and quantitative sequencing of low VAF mutations using only 250x depth, our technology allow lower throughput NGS instruments (e.g. $50k MiniSeq) to analyze clinical samples for low VAF mutations across many genes. The reduced sequencing reads required by mBDA also manifests as a significant reduction of bioinformatics analysis time and data storage. [50] We envision that facilitating the decentralized NGS testing of clinical samples for low VAF somatic mutations will accelerate the adoption of precision medicine and lead to improvements in patient outcomes.

**TABLE 3:**
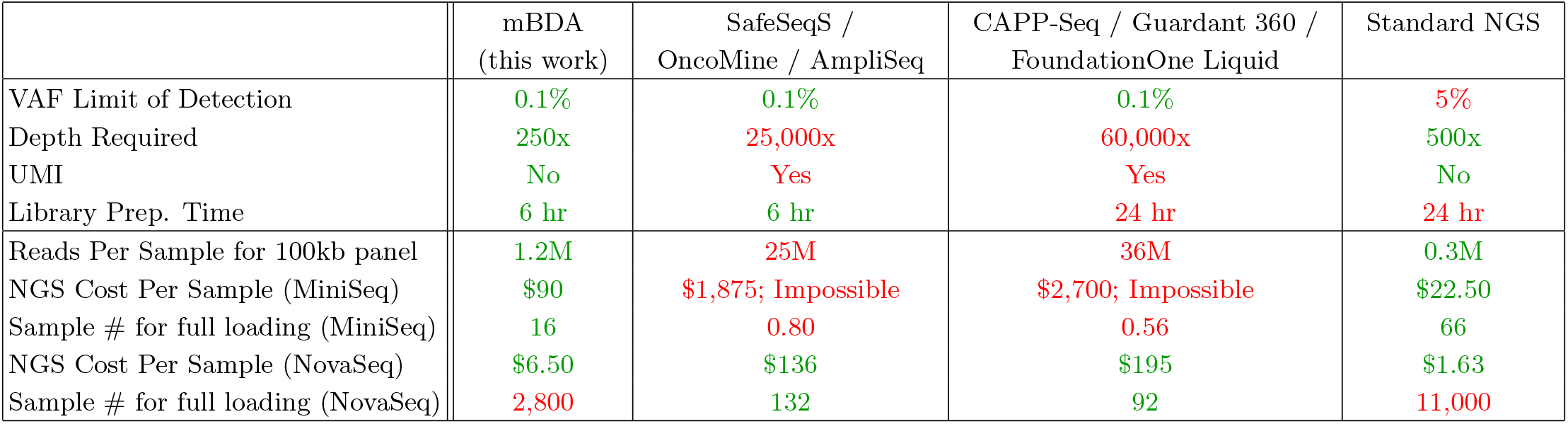
Comparison of performance for different NGS methods for profiling somatic mutations. Only mBDA and standard NGS are feasible on the lower throughput NGS platforms such as the MiniSeq, and only standard NGS is incapable of detecting VAFs of significantly lower than 5%. In contrast, mBDA is unsuitable for high throughput NGS platforms such as the NovaSeq, due to the large number of samples that must be pooled in order to leverage the instruments’ economies of scale. NGS cost for MiniSeq assumes $1,500 for 20M reads on 2×150nt flow cell; NGS cost for NovaSeq assumes $18,000 for 3.3B reads on S2 2×150 flow cell. mBDA assumes an enrichment region of 20 nt per 90 nt amplicon. SafeSeqS, OncoMine, and Ampliseq assumes 140 nt amplicon on average to generate 100 nt of usable sequence. CAPP-Seq, Guardant 360, and FoundationOne Liquid assumes 160 nt DNA input.

|  | mBDA<br>(this work) | SafeSeqS /<br>OncoMine / AmpliSeq | CAPP-Seq / Guardant 360 /<br>FoundationOne Liquid | Standard NGS |
| --- | --- | --- | --- | --- |
| VAF Limit of Detection | 0.1% | 0.1% | 0.1% | 5% |
| Depth Required | 250x | 25,000x | 60,000x | 500x |
| UMI | No | Yes | Yes | No |
| Library Prep. Time | 6 hr | 6 hr | 24 hr | 24 hr |
| Reads Per Sample for 100kb panel | 1.2M | 25M | 36M | 0.3M |
| NGS Cost Per Sample (MiniSeq) | \$90 | \$1,875; Impossible | \$2,700; Impossible | \$22.50 |
| Sample # for full loading (MiniSeq) | 16 | 0.80 | 0.56 | 66 |
| NGS Cost Per Sample (NovaSeq) | \$6.50 | \$136 | \$195 | \$1.63 |
| Sample # for full loading (NovaSeq) | 2,800 | 132 | 92 | 11,000 |

## Supporting information

mBDA Supplemental Excel

## Acknowledgements

This work was supported by NIH grant R01CA203964 and by CPRIT grant RP180147 to DYZ. The authors thank Jianyi Nie for proofreading assistance. The authors thank the Gang Bao for providing access to his Bio-Rad QX200 digital droplet PCR instrument. The authors thank Nuprobe for providing early access VarMap NSCLC kits for cfDNA testing.

## Author contributions

PS and DYZ conceived the project. PS, SXC, and HYY performed mBDA sequence design for cell line contamination panels. LYC and PD performed mBDA sequence design for cancer panels. AAP provided clinical cfDNA samples and performed comparison deep sequencing experiments. PS and HYY performed experiments and analyzed qPCR data. PS and AP performed NGS experiments. PS, SXC, and DYZ analyzed NGS data. PS and DYZ wrote the manuscript with input from all authors.

## Additional information

Correspondence may be addressed to DYZ. There are patents pending on the BDA and mBDA methods used in this work. PS, SXC, LYC, and PD declare competing interests in the form of consulting for Nuprobe USA. AAP declares a competing interest in the form of consulting for Nuprobe USA, and consulting for and significant equity ownership in Binary Genomics. DYZ declares a competing interest in the form of consulting for and significant equity ownership in Nuprobe and Torus Biosystems, and consulting for Avenge Bio. Full code available upon request.

