## Supplementary info for "Detecting and Quantitating Low Fraction DNA Variants with Low-Depth Sequencing"

S0. Experimental Methods 1
S1. SNP Selection and mBDA Design 3
S2. mBDA NGS Protocol Optimization 7
S3. NGS Data Analysis 13
S4. Measurement of Enrichment-Fold (EF) Values 17
S5. NGS Depth and Library Size Needed 22
S6. Variant Allele Detection with Small Input DNA Quantities 24
S7. mBDA qPCR Protocol Optimization 25
S8. Analysis of Conspecific Contaminant Detection using mBDA qPCR Assay 35
S9. Detection of Known Contaminants in Intended Cell Lines of Unknown Genotype 37
S10. Cancer Panel Contents and EF Calibration 41
S11. Cancer Panel Analytic Validation vs ddPCR 46

#### Section S0: Experimental Methods

**Primer and Blocker Oligonucleotides.** Primers and blockers were purchased from Integrated DNA technologies (IDT). All DNA oligos were purchased with standard desalting and LabReady formulation (100 uM in IDT's Tris-EDTA pH 8.0). The DNA stock solutions were diluted using 1× TE buffer (purchased from Sigma Aldrich as 100 × stock solution) to 5$\mu$M. All primers and blockers can be stored in 4 °C.

**Cell Line and Human DNA Samples.** Human gDNA samples (NA18562, NA18537, NA19223, NA12815, NA20507, NA18545, NA18572, and NA20502) were purchased from Coriell Biorepository, and stored at -20 °C. HeLa cell line DNA was ordered from New England Biolabs (NEB). K562, T24, PC3 cells were obtained from collaborators and DNA was extracted using standard methods. Fresh frozen (FF) cancer tissue samples were ordered from OriGene Technologies Inc.

**Volunteer DNA Samples.** Buccal swabs samples were collected from volunteers in the Houston, TX area. Swabs were placed in 400 $\mu$L of 1× PBS buffer in 1.5 ml tubes and incubated in a Multi-Therm instrument (Genesee) at 37 °C for 10 min. The swabs were then removed, and the remaining solution was centrifuged at 12000 rpm for 5 min. Supernatant was discard, and then 200 $\mu$L PBS buffer was added to resuspend the cell pellet, while vortexing vigorously for 1 min. Next, 20 $\mu$L proteinase K and 4 $\mu$L RNase A solution (100 mg/ml) were added, and the suspension was vortexed for 15 sec. Finally, the solution was extracted using a QIAamp DNA Blood Mini Kit (Qiagen) following manufacturer instructions.

Blood samples were purchased commercially from ZenBio, Inc. from consented de-identified volunteers in the Chapel Hill, NC area. Blood samples were centrifuged at 500 pm for 15 min to separate plasma, red blood cells, and buffy coat. DNA was extracted from the buffy coat using QiaAMP DNA Blood Mini Kit (Qiagen) following manufacturer instructions.

**NGS Library Preparation Protocol for mBDA.** For each library, the DNA sample was first mixed with appropriate concentrations of primers and blockers, and then underwent 23 cycles of PCR, using Phusion Hot Start Flex DNA polymerase (NEB). Here, the reaction volume is 50 $\mu$L, and the thermocycling protocol including an initial 98 °C denaturation for 30 s, and then 23 cycles of 98 °C 10 s, 63 °C 5 min, and 72 °C 2 min. The amplicon products were then purified from the reaction mixture using a column-based DNA Clean & Concentrator kit (Zymo Research).

Next, we append sequencing adaptors to the BDA amplicons via 2 cycles of PCR using adapter primers. The forward and reverse adapter primers each have a final reaction concentration of 15 nM in a reaction volume of 50 $\mu$L. The same PCR thermocycling protocol as above was applied for 2 cycles, and the amplicons were subsequently re-purified using the DNA Clean & Concentrator kit. We then optionally quantitate the adaptor-appended amplicons using qPCR (BioRad iTaq SYBR Green Supermix; 95 °C 3 min, then 95°C 10 s, 60 °C 30 s for 40 cycles). The observed cycle threshold (Ct) value was determined via BioRad software.

Next, we perform index PCR, using the Illumina Nextera sequences as index primers. Here, we used Phusion Hot Start Flex DNA polymerase, and added 5 $\mu$L of index primers into the final 50 $\mu$L reaction, and followed the following thermocycling protocol: 98 °C 30 s, then 98 °C 10 s, 63 °C 1 min, 72 °C 1 min for 11 cycles. If qPCR was used in the previous step, we would instead perform index PCR for (Ct + 4) cycles.

Before the next step of amplicon size selection, we incubated Ampure beads at room temperature for 30 min. The prepared Ampure beads were then used to size select the NGS library; 0.7× and 0.3× Ampure beads ratio were used (i.e. 0.7× means 50 $\mu$L of library and 35 $\mu$L of Ampure beads). After size selection, we used a Qubit dsDNA HS kit (Thermo Fisher) to quantify the concentration of the library, and used a Bioanalyzer DNA 1000 kit (Agilent) was used to quantify the length of the amplicons to ensure library quality. Different libraries were pooled into a final concentration of 4 nM, and loaded onto an Illumina MiSeq instrument following standard protocols, with between 5% and 10% of PhiX.

**NGS Library Preparation Protocol for Standard Multiplex PCR Amplicons (No Blockers).** The protocol is mostly the same as the mBDA NGS library preparation protocol, except the number of PCR cycles in the first step is reduced to 13 cycles from 23 cycles. Additionally, for historical reasons, during size selection we used 0.6× and 0.3× Ampure beads (instead of 0.7× and 0.3×); we do not expect this difference to make a significant difference in our conclusions.

**qPCR Protocol for mBDA.** For all qPCR mBDA experiments, we used the PowerUp SYBR Green qPCR Master Mix (Thermo Fisher). qPCR was performed in a CFX96 Touch Real-Time PCR Detection System using 96-well plates. Exact thermocycling protocols varied for different experiments, and are described in Section S2. qPCR reactions were typically performed in replicates of 3 to 12 reactions, with 10 $\mu$L per reaction. In all reactions, the Blocker concentration was set at 10-fold higher than the forward primer concentration.

**NSCLC NGS panel for cfDNA.** The NSCLC mBDA NGS panel for cfDNA was run on clinical cfDNA samples based on manufacturer instructions (Nuprobe VarMap NSCLC kit). Clinical cfDNA samples were obtained from the laboratory of Dr. Patel, who previously had performed deep sequencing analysis as comparison experiments.

**ddPCR Protocol for cancer panel mBDA.** The ddPCR experiments were performed as comparison experiments for analytically validating the melanoma mBDA NGS panel. We ordered official Bio-Rad Laboratory kits containing primers and probes for detecting NRAS G12S and Q61K mutations. For the MAP2K2 p. C125S mutation, we manually designed probes following suggestions in the Bio-Rad QX200 user manual. For all ddPCR input experiments, 17ng of input DNA were used for each reaction. The final primer concentrations were 900nM and probe concentration were 250nM, and the PCR enzyme used was the Bio-Rad 2x ddPCR supermix for probes (no dUTP). The PCR protocol applied was: 95°C, 10min - (94°C, 30s - 55°C, 1min) $\times$ 40 cycles - 98°C, 10min - 4°C hold.

#### Section S1: SNP Selection and mBDA Design

**SNP selection.** Fig. S1-1a shows the genomic positions of the 80 single nucleotide polymorphisms (SNPs) that we used for the majority of studies in this manuscript, based on the Human Genome GRCh37 build. For each SNP, the minor allele's population allele fraction (PAF) was downloaded from the 1000 Genomes database as stored at the National Center for Biotechnology Information (NCBI) website. The SNP loci were selected such that they were at least 10,000 nt away from each other on the genome. The SNPs can be classified into 12 types based on the sequences of the reference and minor allele; the 80 SNPs selected cover all 12 types (Fig. S1-1b).

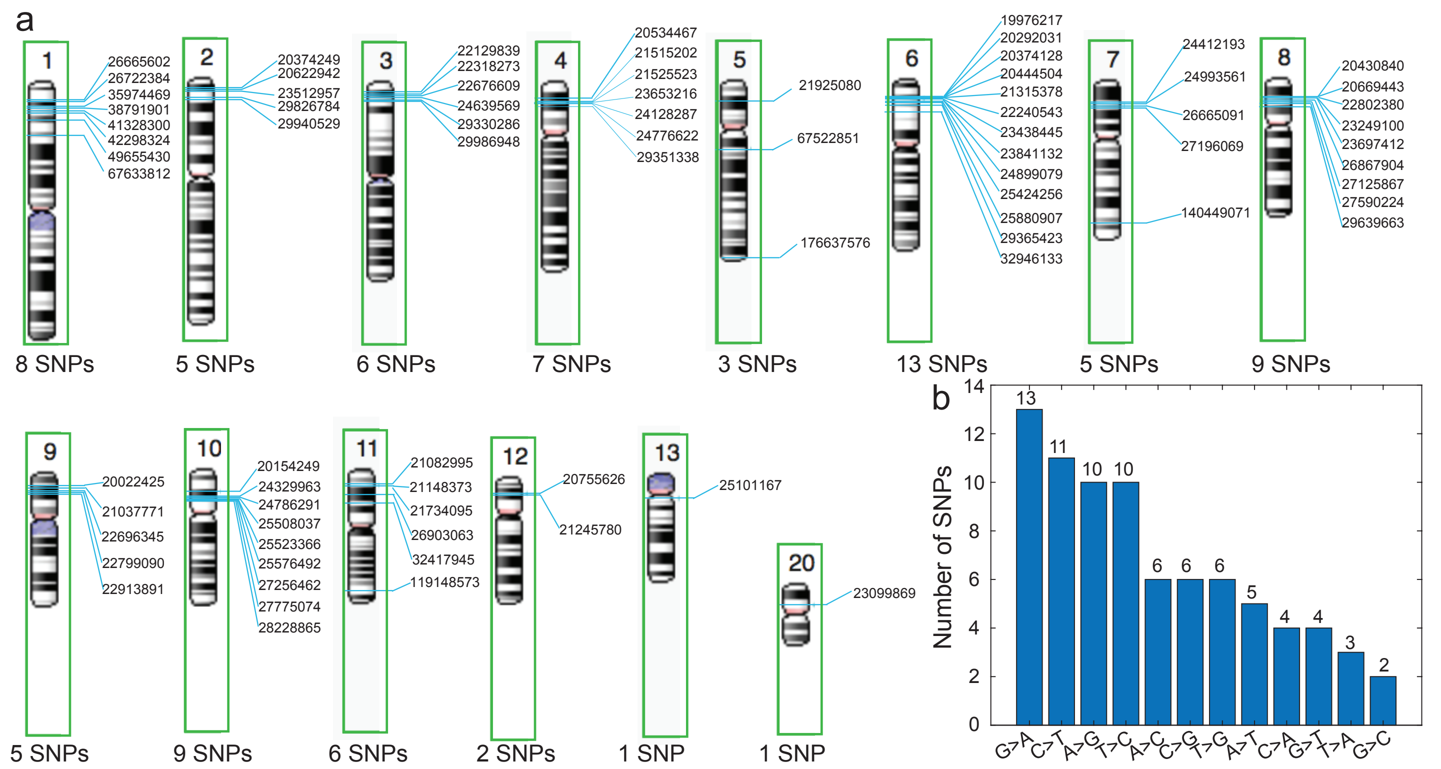


FIG. S1-1. List of 80 SNPs selected for experimental work. **(a)** Chromosomal positions of the 80 SNP loci, based on GRCh37. **(b)** Sequence identity distribution of reference and minor SNP alleles.

**BDA design.** A number of allele-specific PCR methods have been demonstrated over the past 2 decades, with some methods such as castPCR, and ICE-COLD PCR demonstrating selectivity for single-nucleotide variants with a limit of detection (LoD) of roughly 0.1% variant allele frequency (VAF). However, these methods generally require very accurate and uniform temperature control to achieve such selectivity, and thus have not been shown to exhibit similar LoDs in multiplexed settings. Thus, it is very difficult to design a 21-plex assay based on these technologies to detect arbitrary conspecific contaminants. Although it is possible to run 21 separate single-plex assays to detect low fraction contaminants, the significantly higher labor as well as the increased number of different reagents makes this approach unattractive.

Blocker displacement amplification (BDA) is a PCR-based method for selective amplification of DNA templates with sequence variants in a roughly 20 nt region of interest (pink region in Fig. S1-2). A rationally designed Blocker oligonucleotide hybridizes perfectly to DNA template molecules with the intended allele, and forms a mismatch bubble with DNA templates with the variant allele. The Blocker's binding region overlaps with that of the forward primer (FP), such that the binding of the FP and the Blocker is thermodynamically mutually exclusive. The Blocker is typically included in the reaction at 10× higher concentration than the FP, in order to favor the initial binding of the Blocker. Thus, in order for PCR amplification to proceed, the FP must displace the Blocker.

The lengths and sequences of the FP and the Blocker are designed such that the FP displacing the Blocker is thermodynamically unfavorable for templates with the intended allele, but thermodynamically favorable for templates with the variant allele. The FP binding yield thus limits the per-cycle PCR amplification yield, and through the course of many PCR cycles, the relative abundance of amplicons with the variant allele is increased dramatically over the relative abundance of the intended allele.

The fold-enrichment of the variant allele over the intended allele will vary based on the relative binding energies of the FP and Blocker, as well as the energetics of the mismatch bubble formed between the Blocker and the variant allele template. Because G-T wobbles are known to be thermodynamically less destabilizing that all other single-base mismatch bubbles, where the SNP allele identities are T>C or G>A, we will design the Blocker and FP against the (-) strand of the DNA, rather than the default (+) strand. Table S1-1 shows the rs numbers, the DNA strand used, the reference allele, the minor allele PAF, the NA18537 and NA18562 alleles, and the genomic coordinates for all 80 SNPs we used.

| Multiplex BDA | SNP rs- | Strand (+/-) | Ref 1000 | PAF | NA18537 (intended) | NA18562(Variant) | Chr | Position(GRCh37) |
|---|---|---|---|---|---|---|---|---|
| mb1 | rs10230708 | - | A | 0.45 | G | A | 7 | 26,665,091 |
| mb2 | rs10104396 | + | C | 0.84 | C | T | 8 | 23,697,412 |
| mb3 | rs199032 | - | T | 0.49 | T | C | 6 | 23,438,445 |
| mb4 | rs926850 | + | A | 0.27 | A | G | 6 | 24,899,079 |
| mb5 | rs17149369 | + | G | 0.94 | A | G | 7 | 24,412,193 |
| mb6 | rs869720 | + | A | 0.55 | C | A | 1 | 38,791,901 |
| mb7 | rs12478327 | + | A | 0.51 | C | A | 2 | 20,374,249 |
| mb8 | rs2638145 | + | C | 0.46 | C | A | 3 | 22,318,273 |
| mb9 | rs2170091 | + | A | 0.54 | A | C | 3 | 22,129,839 |
| mb10 | rs2043583 | + | G | 0.52 | T | G | 3 | 24,639,569 |
| mb11 | rs955456 | + | T | 0.61 | T | G | 4 | 23,653,216 |
| mb12 | rs966516 | + | C | 0.85 | A | C | 4 | 24,776,622 |
| mb13 | rs354169 | + | T | 0.56 | A | T | 1 | 49,655,430 |
| mb14 | rs1898170 | + | T | 0.37 | T | A | 4 | 29,351,338 |
| mb15 | rs11247921 | + | G | 0.79 | G | C | 1 | 26,665,602 |
| mb16 | rs1635718 | + | C | 0.38 | G | C | 1 | 35,974,469 |
| mb17 | rs10510620 | + | C | 0.78 | C | G | 3 | 29,330,286 |
| mb18 | rs7104025 | + | C | 0.4 | C | G | 11 | 21,148,373 |
| mb19 | rs2246745 | + | T | 0.59 | T | A | 2 | 29,940,529 |
| mb20 | rs3789806 | + | C | 0.65 | C | G | 7 | 140,449,071 |
| mb21 | rs706714 | + | A | 0.58 | A | C | 5 | 67,522,851 |
| mb22 | rs1884444 | + | G | 0.53 | T | G | 1 | 67,633,812 |
| mb23 | rs2510152 | + | G | 0.27 | G | T | 11 | 119,148,573 |
| mb24 | rs16754 | + | T | 0.67 | T | C | 11 | 32,417,945 |
| mb25 | rs206781 | + | A | 0.26 | G | A | 6 | 32,946,133 |
| mb26 | rs28932178 | + | T | 0.25 | C | T | 5 | 176,637,576 |
| mb27 | rs10186821 | - | A | 0.44 | G | A | 2 | 20,622,942 |
| mb28 | rs10508599 | + | T | 0.72 | T | G | 10 | 20,154,249 |
| mb29 | rs10738578 | + | C | 0.7 | C | T | 9 | 21,037,771 |
| mb30 | rs10741037 | - | G | 0.55 | G | A | 10 | 24,329,963 |
| mb31 | rs10770674 | + | C | 0.18 | C | T | 12 | 20,755,626 |
| mb32 | rs10805227 | + | A | 0.49 | A | G | 4 | 21,515,202 |
| mb33 | rs10833604 | + | T | 0.11 | C | T | 11 | 21,734,095 |
| mb34 | rs10964389 | + | T | 0.24 | C | T | 9 | 20,022,425 |
| mb35 | rs11015816 | + | T | 0.92 | C | T | 10 | 27,775,074 |
| mb36 | rs11045749 | + | T | 0.87 | A | T | 12 | 21,245,780 |
| mb37 | rs1123828 | - | T | 0.17 | T | C | 8 | 22,802,380 |
| mb38 | rs11708584 | + | A | 0.54 | A | G | 3 | 29,986,948 |
| mb39 | rs12192635 | + | A | 0.84 | A | T | 6 | 25,880,907 |
| mb40 | rs12213948 | - | G | 0.44 | G | A | 6 | 25,424,256 |
| mb41 | rs12259813 | - | T | 0.56 | T | C | 10 | 27,256,462 |
| mb42 | rs12541300 | + | C | 0.63 | C | G | 8 | 20,430,840 |
| mb43 | rs12681931 | - | C | 0.34 | T | C | 8 | 23,249,100 |
| mb44 | rs12782580 | + | A | 0.75 | A | G | 10 | 24,786,291 |
| mb45 | rs1375977 | + | A | 0.6 | A | G | 11 | 26,903,063 |
| mb46 | rs1516755 | - | A | 0.32 | G | A | 11 | 21,082,995 |
| mb47 | rs1524303 | - | T | 0.32 | T | C | 3 | 22,676,609 |
| mb48 | rs1667087 | + | G | 0.66 | A | G | 9 | 22,696,345 |
| mb49 | rs16871316 | + | G | 0.07 | A | G | 4 | 21,525,523 |
| mb50 | rs16925478 | + | G | 0.79 | G | T | 10 | 25,523,366 |
| mb51 | rs17560702 | + | T | 0.5 | C | T | 6 | 19,976,217 |
| mb52 | rs1937037 | + | G | 0.65 | G | T | 9 | 22,913,891 |
| mb53 | rs2215492 | + | T | 0.6 | C | T | 4 | 24,128,287 |
| mb54 | rs2301720 | + | A | 0.23 | A | C | 7 | 27,196,069 |
| mb55 | rs2616187 | - | G | 0.56 | G | A | 8 | 20,669,443 |
| mb56 | rs2710998 | - | A | 0.36 | G | A | 7 | 24,993,561 |
| mb57 | rs2807238 | + | C | 0.81 | C | G | 10 | 25,508,037 |
| mb58 | rs2874755 | + | C | 0.24 | A | C | 8 | 26,867,904 |
| mb59 | rs3813787 | + | A | 0.77 | A | C | 1 | 41,328,300 |
| mb60 | rs4665582 | + | T | 0.7 | T | G | 2 | 23,512,957 |
| mb61 | rs4712476 | + | T | 0.76 | G | T | 6 | 20,292,031 |
| mb62 | rs611628 | - | C | 0.47 | T | C | 1 | 42,298,324 |
| mb63 | rs6452035 | - | A | 0.72 | G | A | 5 | 21,925,080 |
| mb64 | rs6816854 | + | C | 0.48 | C | T | 4 | 20,534,467 |
| mb65 | rs6937778 | - | A | 0.39 | G | A | 6 | 20,374,128 |
| mb66 | rs7003044 | - | C | 0.28 | T | C | 8 | 27,125,867 |
| mb67 | rs7032336 | - | C | 0.66 | T | C | 9 | 22,799,090 |
| mb68 | rs7816009 | + | T | 0.35 | C | T | 8 | 29,639,663 |
| mb69 | rs7893462 | - | A | 0.53 | G | A | 10 | 28,228,865 |
| mb70 | rs7902135 | - | G | 0.82 | G | A | 10 | 25,576,492 |
| mb71 | rs898476 | + | A | 0.52 | A | G | 8 | 27,590,224 |
| mb72 | rs9368431 | + | A | 0.9 | A | T | 6 | 22,240,543 |
| mb73 | rs9438621 | - | G | 0.28 | G | A | 1 | 26,722,384 |
| mb74 | rs9466035 | + | G | 0.54 | A | G | 6 | 21,315,378 |
| mb75 | rs9466930 | + | C | 0.37 | C | G | 6 | 23,841,132 |
| mb76 | rs9973865 | - | T | 0.2 | T | C | 2 | 29,826,784 |
| mb77 | rs4712498 | + | A | 0.16 | A | T | 6 | 20,444,504 |
| mb78 | rs2073149 | + | A | 0.38 | T | A | 6 | 29,365,423 |
| mb79 | rs2862909 | + | G | 0.65 | T | G | 13 | 25,101,167 |
| mb80 | rs1338945 | + | C | 0.38 | C | A | 20 | 23,099,869 |

TABLE S1-1. Details of the 80 SNP loci selected for mBDA analysis. Here, we display the rs numbers, the DNA strand used, the reference allele, the minor allele PAF, the NA18537 and NA18562 alleles, and the genomic coordinates.

Through the course of our internal calibration experiments (data not shown), we have found that the optimal standard free energies of the three regions (FP-unique, overlap between FP and Blocker, and Blocker-unique) are

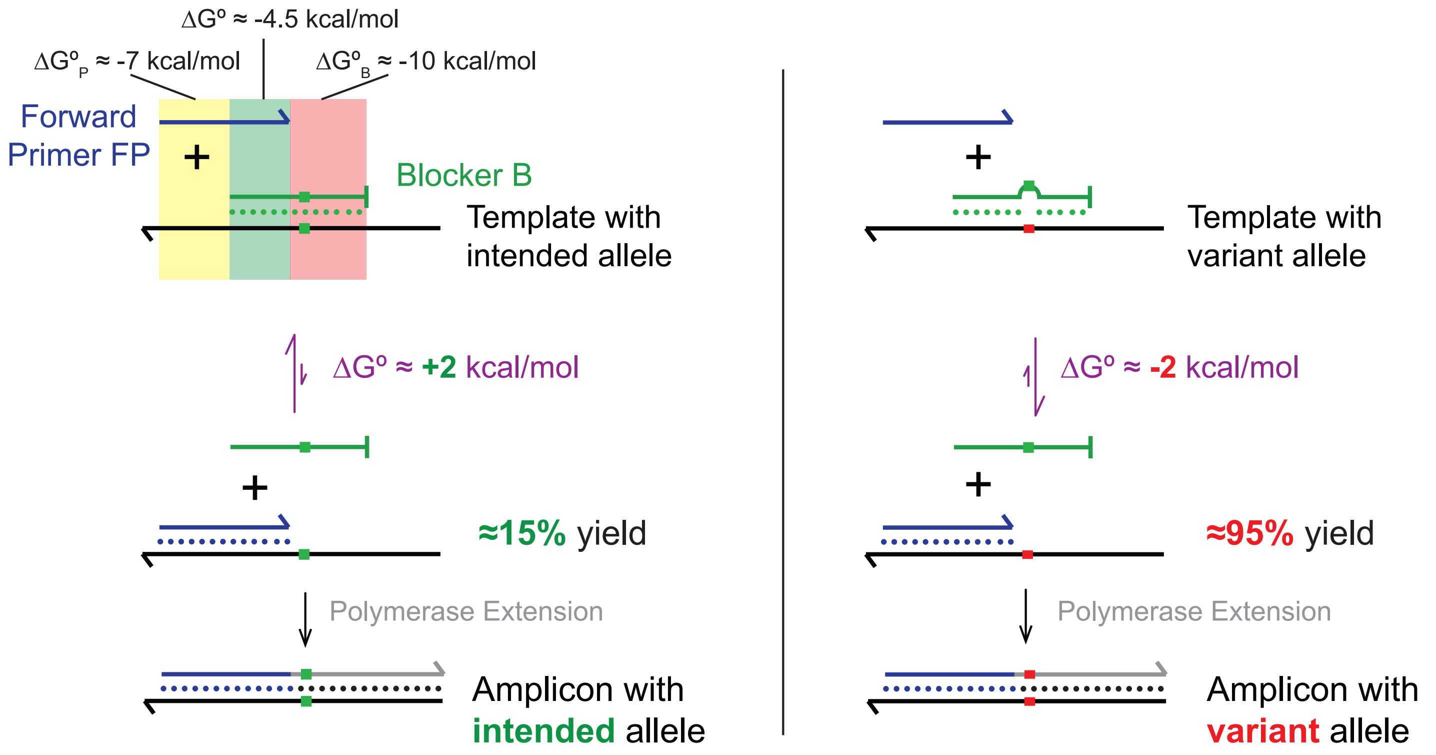


FIG. S1-2. BDA design detail and energy calculation. Yellow boxes show the toehold region, green boxes show the overlap region and pink boxes show the nonhomologous region. Blockers are complementary to primary templates so that FP displacing the blocker from primary template is difficult. This will cause the primary amplicon with low amplification yield. However, variant template bind with blocker is very unstable, so that FP will be easier to displace the blocker to get higher yield variant amplicons. The nonhomologous region is around 20 nt while effective enrichment region is 16 nt (20 nt—4 nt).

$\Delta G^\circ_P$ = -7 kcal/mol, -4.5 kcal/mol, and $\Delta G^\circ_B$ = -10 kcal/mol, respectively (Fig. S1-2). Insofar as sequence-based thermodynamics prediction software (e.g. Nupack, mFold) are imperfect, with errors of up to 1 kcal/mol, some empirical adjustment of the FP and Blocker sequences are needed in certain cases. For example, when the variant allele fails to efficiently PCR amplify, we would increase the length of the FP-unique region. When the enrichment-fold (EF) of the variant over the intended allele is lower than expected, we would either decrease the length of the FP-unique region or increase the length of the Blocker-unique region. Two examples BDA systems are shown with calculated thermodynamics in Fig. S1-3.

To ensure that the Blockers themselves do not act as primers and extend off the templates, we typically will use Blockers with a 3′ modification that prevents enzymatic extension. For qPCR experiments that use the Taq-based PowerUP enzyme mix (Thermo Fisher), a 3-carbon spacer was sufficient. For NGS library preparation experiments that use the high fidelity Phusion enzyme mix (New England Biolabs), we used Blockers with 3′ DXXDM, wherein D denotes matched DNA nucleotide, X denotes internal C3 spacer, and M denotes a mismatched DNA nucleotide.

**mBDA design**. Here, the design of multiplex BDA (mBDA) primers and blockers sequences began with individual design of single-plex BDA sequences. Subsequently, the 80-plex mBDA set was checked for potential primer dimers and nonspecific genomic amplification, as follows: The sequences for every pair of primers in the 160 primer set was compared for 3′ reverse complementarity of at least 5 nucleotides. Where such 3′ reverse complementarity was found, one of the two primers that form the potential primer dimer was randomly selected for redesign, with preference given to redesign of reverse primers (RP) because there are relatively more possible alternative sequences. This process was repeated until the entire 160 primer set had no pairwise 3′ reverse complementarity. Subsequently, each primer was individually subject to BLAST against the human genome, and any primers with more than 5 exact hits on the human genome were replaced by an alternative sequence. The replaced primer is checked for potential primer dimers against the rest of the primer set.

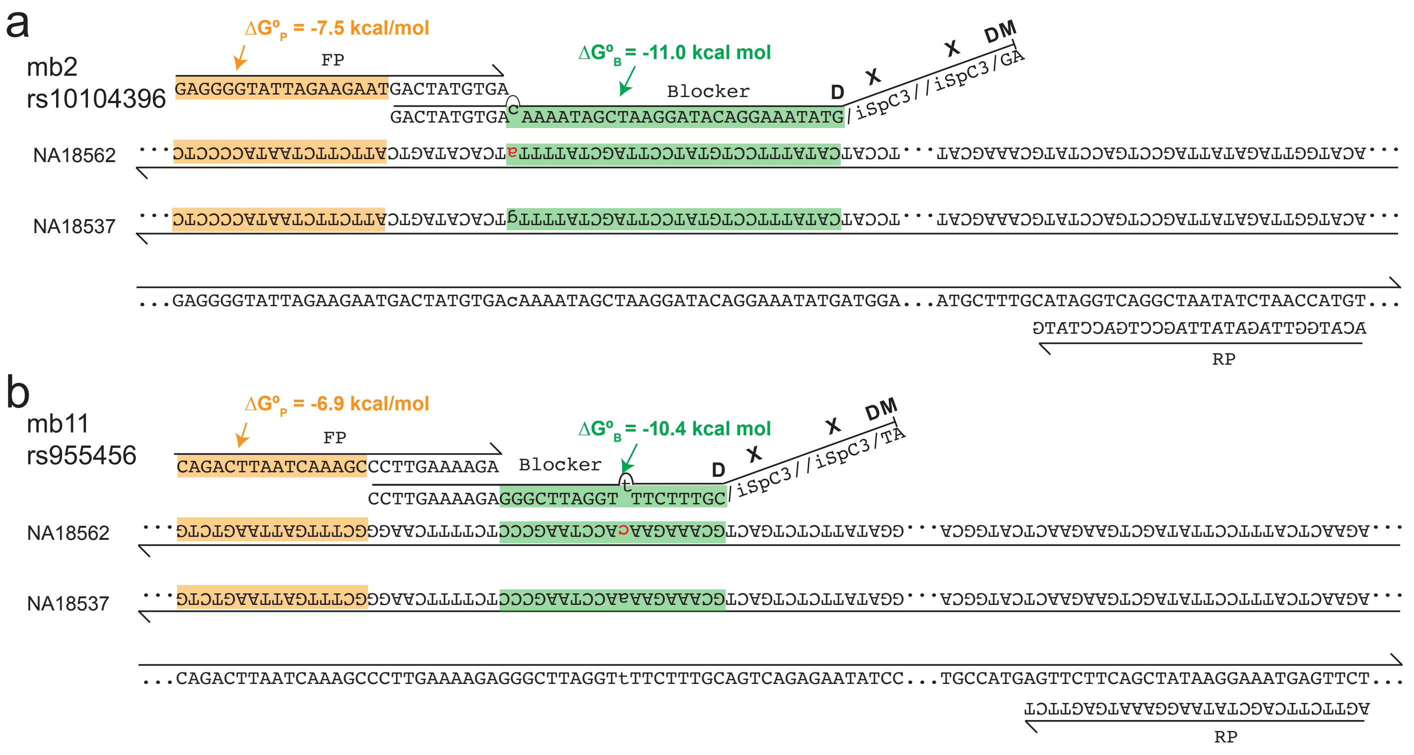


FIG. S1-3. Two examples of BDA designs for the mb2 and mb11 SNPs (rs10104396 and rs955456, respectively), based on the NA18537 allele as the intended allele.

#### Section S2: mBDA NGS Protocol Optimization

**Enrichment-Fold (EF) definition and mathematics.** The value of EF determines the relationship between the variant allele frequency (VAF) in the original sample and the variant read fraction (VRF) observed from NGS reads.

The initial ratio of variant allele DNA molecules to intended allele DNA molecules is VAF : (1 - VAF). Assuming that the intended DNA is amplified by a factor of $C$ through mBDA, the variant DNA is amplified by a factor of EF $\cdot C$, so the final ratio is EF$\cdot C \cdot$ VAF : $C \cdot$ (1 - VAF). Thus, the VRF can be calculated as:

$$\mathrm{VRF} = \frac{\mathrm{VAF} \times \mathrm{EF}}{\mathrm{VAF} \times \mathrm{EF} + 1 - \mathrm{VAF}} \tag{1}$$

For mathematical simplicity, we define two new variables $\chi$ and $\psi$ as follows:

$$\chi = \frac{\mathrm{VRF} - 1}{\mathrm{VRF}} \tag{2}$$

$$\psi = \frac{\mathrm{VAF} - 1}{\mathrm{VAF}} \tag{3}$$

Based on these definitions, VRF can be redefined as a function of $\chi$:

$$\mathrm{VRF} = \frac{1}{1 - \chi} \tag{4}$$

Equation (1) from above can be rewritten as:

$$\begin{aligned} \frac{\frac{1}{1-\psi} \times \mathrm{EF}}{\frac{1}{1-\psi} \times \mathrm{EF} + 1 - \frac{1}{1-\psi}} &= \frac{1}{1-\chi} && (5) \\ \frac{\mathrm{EF}}{\mathrm{EF} + 1 - \psi + 1} &= \frac{1}{1-\chi} && (6) \\ \frac{\mathrm{EF}}{\mathrm{EF} - \psi} &= \frac{1}{1-\chi} && (7) \\ \mathrm{EF} &= \frac{\psi}{\chi} = \frac{(\frac{1-\mathrm{VAF}}{\mathrm{VAF}})}{(\frac{1-\mathrm{VRF}}{\mathrm{VRF}})} && (8) \end{aligned}$$

**VAF Limit of Detection.** Given a value of EF from calibration experiments, we are able to infer the value of VAF based on the VRF present in an NGS library.

$$\mathrm{VAF} = \frac{\mathrm{VRF}}{(\mathrm{EF} - 1) \cdot (1 - \mathrm{VRF}) + 1} \tag{9}$$

However, because of sequencing errors and DNA polymerase misincorporation errors, the actual inferred VAF may be different (typically higher) than the real VAF. To characterize the VAF limit of detection (LoD), we performed a number of mBDA NGS experiments on samples known to be pure in the intended SNP alleles. The inferred VAF from the observed VRF for these libraries shows the range of error rates observed for different SNP alleles and genomic loci, and also serves as a lower bound for the VAF LoD.

Because a number of protocol details will influence both the EF and the VAF LoD values, here we tested and characterized multiple different enzymes, oligo designs, and library preparation protocols for mBDA NGS library construction.

**DNA polymerase and mBDA thermodynamics optimization.** We tested three commonly-used high-fidelity DNA polymerases with $3'$-$5'$ exonuclease activity that reduces DNA polymerase misincorporation error rates: Q5, Phusion, and KAPA. For these studies, we used a 10-plex mBDA NGS library, with amplicons for the SNPs internally referred to as mb1, mb2, mb3, mb4, mb8, mb9, mb11, mb16, mb20, and mb81.

Our results (Fig. S2-1abc) show that (1) KAPA and Phusion exhibit marginally lower error rates than Q5, (2) Phusion and Q5 have significantly better NGS read uniformity than KAPA, and (3) Phusion tends to exhibit higher EF values than Q5. Consequently, we chose to move forward with Phusion for all mBDA NGS studies.

We next examined the thermodynamic design parameters for BDA. Our previous studies on single-plex BDA qPCR suggested designing the primer with the primer-unique sequence having a standard free energy of binding of $\Delta G^\circ_P = -7$ kcal/mol and the Blocker with the Blocker-unique sequence having a standard free energy of binding of $\Delta G^\circ_N = -10$ kcal/mol. We verified these design guidelines in NGS by observing the EF values for the 10-plex mBDA (Fig. S2-1def), and confirmed that $\Delta G^\circ_P = -7$ kcal/mol and $\Delta G^\circ_N = -10$ kcal/mol provides near optimal EF values.

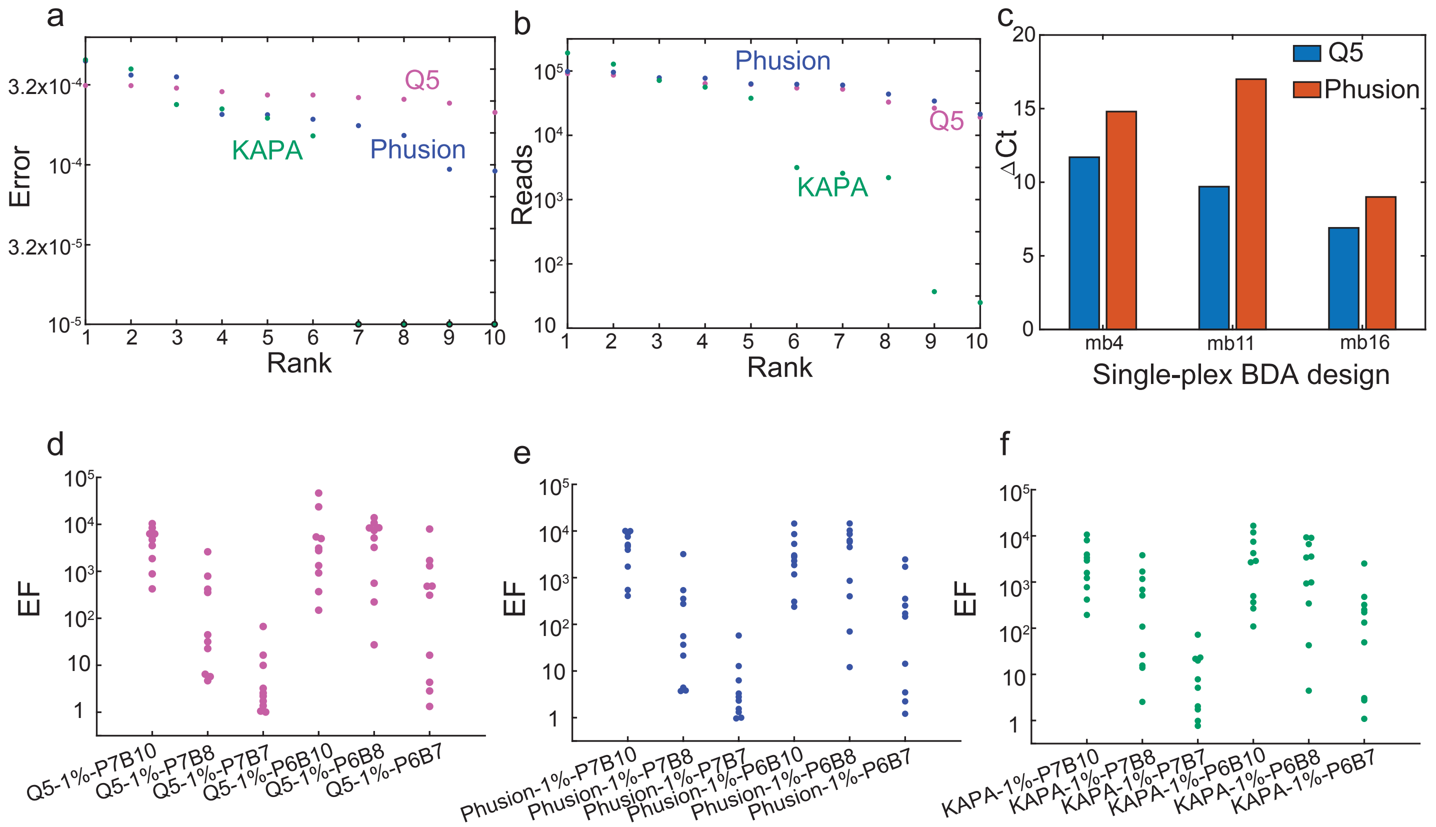


Figure S2-1: Comparison of DNA polymerases and mBDA design methodologies. **(a)** Observed error rate (ranked) for the different three enzymes on a 10-plex mBDA library. **(b)** NGS library uniformity using the 3 different DNA polymerases without primer concentration adjustment. **(c)** Single-plex BDA qPCR to analyze EF for different DNA polymerases. **(d)** EF values for different $\Delta G^\circ_P$ and $\Delta G^\circ_N$ values, when using the Q5 DNA polymerase. **(e)** EF values for different $\Delta G^\circ_P$ and $\Delta G^\circ_N$ values, when using the Phusion DNA polymerase. **(f)** EF values for different $\Delta G^\circ_P$ and $\Delta G^\circ_N$ values, when using the KAPA HiFi DNA polymerase.

**mBDA cycle optimization.** Based on our understanding of BDA's mechanism, increasing the number of PCR cycles during BDA will tend to increase EF values, and could be desirable. However, using a large number of PCR cycles can also lead to two undesirable outcomes: First, the uniformity of the amplicons in the final NGS library is worsened as slight amplification yield biases across different primers compound for a greater number of cycles, and this can only be partially mitigated by adjusting primer concentrations. Second, it is well known that the DNA polymerase misincorporation error rate increases towards later PCR cycles, due to the uneven depletion of different dNTPs, buildup of pyrophosphates, and change in pH. Thus, it is likely that there is a moderate number of PCR cycles for BDA that achieves a desirable tradeoff between high EF values and low error rates.

Fig. S2-2 shows our observed EF values for the 10-plex mBDA NGS panel using different number of PCR cycles for mBDA. To ensure sufficient DNA was available as input for downstream NGS, we applied different number of cycles of index PCR afterwards. EF values do not appear to increase further after roughly 23 cycles. Simultaneously, we did not observe significant increase in error rates for up to 23 cycles. Consequently, we used 23

| | w/o B | | w/B | | | | |
|---|---|---|---|---|---|---|---|
| Q5 | NA18537 (intended) | NA18562 (variant) | NTC | NA18537 (intended) | NA18562 (variant) | ΔCt | Delay |
| mb4 | 17.7 | 18.2 | >40 | 30.9 | 19.2 | 11.7 | 1.0 |
| mb11 | 18.8 | 19.2 | >40 | 31.6 | 21.9 | 9.7 | 2.7 |
| mb16 | 17.7 | 18.3 | 36.7 | 31.6 | 24.7 | 6.9 | 6.4 |

| | w/o B | | w/B | | | | |
|---|---|---|---|---|---|---|---|
| Phusion | NA18537 (intended) | NA18562 (variant) | NTC | NA18537 (intended) | 18562 (variant) | ΔCt | Delay |
| mb4 | 18.6 | 20 | >40 | 35 | 20.2 | 14.8 | 0.2 |
| mb11 | 20.6 | 21 | >40 | 40 | 23 | 17 | 2 |
| mb16 | 19 | 20.6 | >40 | 33 | 24 | 9 | 3.4 |

Table S2-1: Comparison of Q5 and Phusion enzymes for EF value on single-plex qPCR BDA experiments. Primer concentration were 15 nM each. Thermocycling protocol includes annealing at 60 °C for 5 min and extension at 72 °C for 2 min.

cycles as our default mBDA protocol.

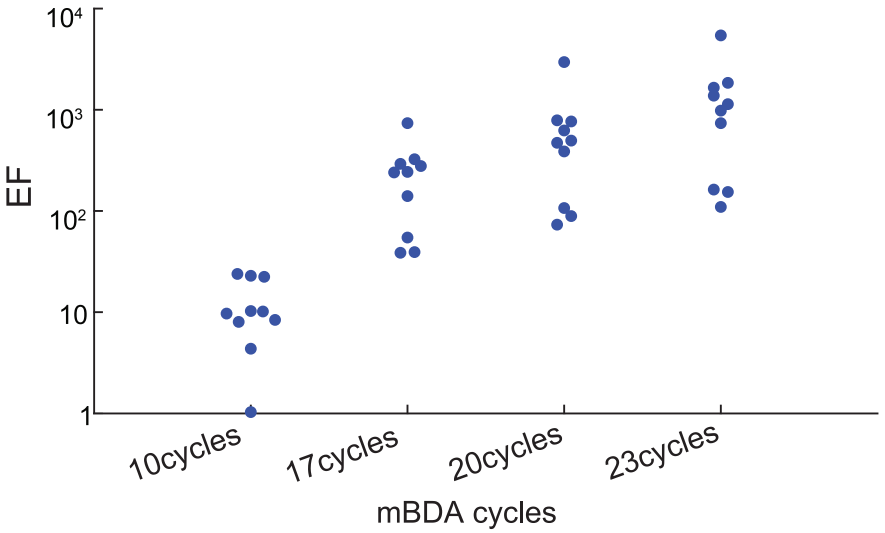


Figure S2-2: EF values as a function of the number of PCR cycles applied for mBDA. Here, we used a 10-plex mBDA NGS comprising forward primers, Blockers, and reverse primers to mb1, mb2, mb3, mb4, mb8, mb9, mb11, mb16, mb20, and mb81.

**Methods for Appending Adapter Sequences.** We considered 3 methods for appending Illumina adapters to our amplicon mixture generated from mBDA: the ligation-based NEB Ultra 2 kit, a two-step PCR method, and blunt-end ligation of adapters (Fig. S2-3). We compared NGS library on-target rates and error rates, and noted that two-step PCR provided both the best on-target rate and the best median error rate, and thus proceeded with two-step PCR as the default library preparation method.

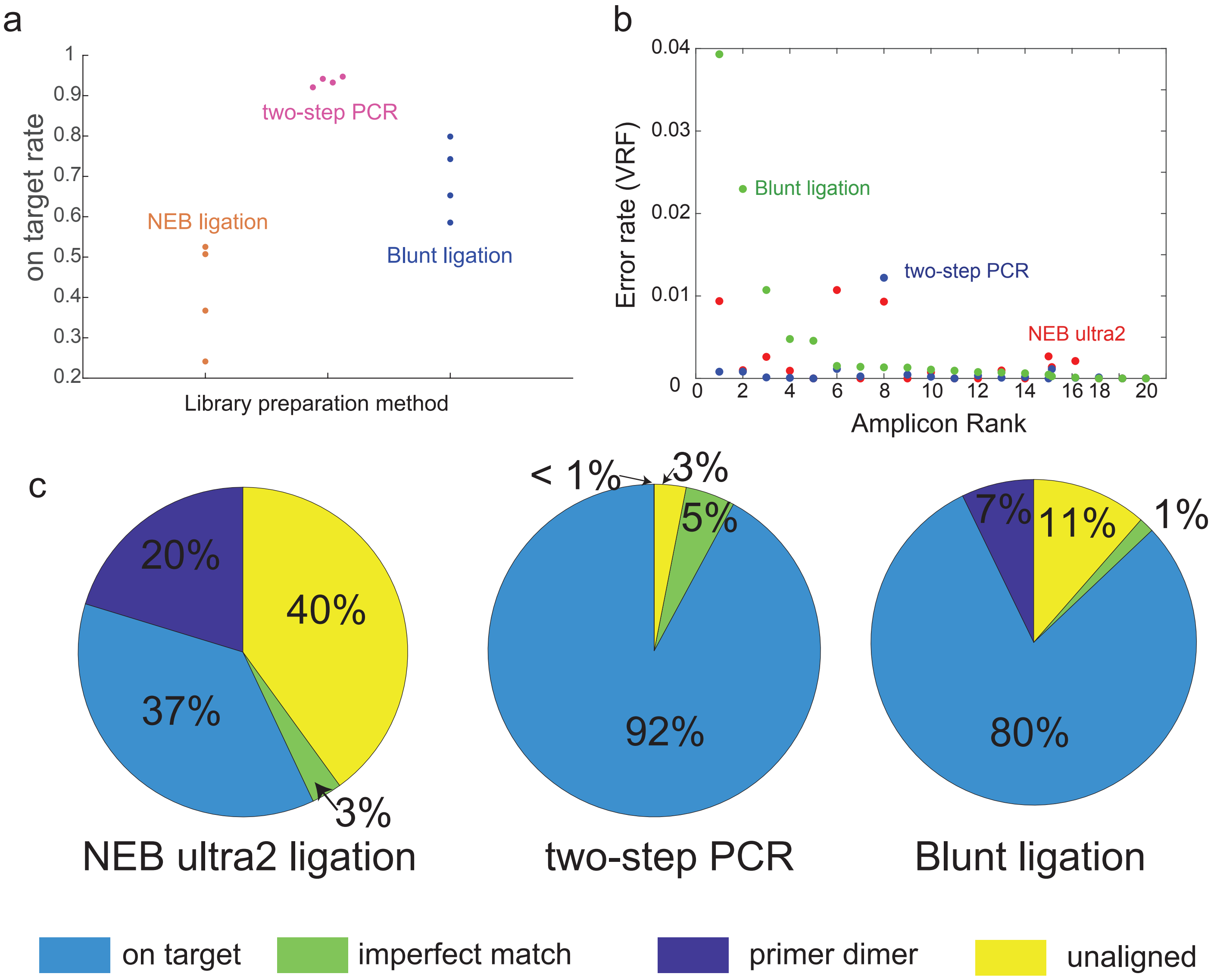


Figure S2-3: Comparison of 3 different NGS library preparation methods. **(a)** On-target rate analysis. **(b)** Error rate analysis of three different methods. **(c)** on target rate, primer dimer rate, unaligned rate, and imperfect matched rate analysis of three methods. Imperfect matched rate means those reads can be aligned but can not perfect matched to target reads, maybe some error in the reads.

| SNP locus | SNP rs # | $\Delta G^\circ_P$ | $\Delta G^\circ_B$ | EF | Strand(+/-) | 1000G ref | 18537 | NA18562 | Chr | Position |
|---|---|---|---|---|---|---|---|---|---|---|
| mb2 | rs10104396 | -7.5 | -11 | 924 | + | C | C | T | 8 | 23,697,412 |
| mb5 | rs17149369 | -7.1 | -10.2 | 1485 | + | G | A | G | 7 | 24,412,193 |
| mb6 | rs869720 | -7.1 | -10.5 | 8254 | + | A | C | A | 1 | 38,791,901 |
| mb7 | rs12478327 | -7 | -10.5 | 1232 | + | A | C | A | 2 | 20,374,249 |
| mb8 | rs2638145 | -7.3 | -10.7 | 3607 | + | C | C | A | 3 | 22,318,273 |
| mb9 | rs2170091 | -7.4 | -10.7 | 1123 | + | A | A | C | 3 | 22,129,839 |
| mb10 | rs2043583 | -7.2 | -10.2 | 970 | + | G | T | G | 3 | 24,639,569 |
| mb11 | rs955456 | -6.9 | -10.4 | 720 | + | T | T | G | 4 | 23,653,216 |
| mb12 | rs966516 | -7.1 | -10.2 | 310 | + | C | A | C | 4 | 24,776,622 |
| mb13 | rs354169 | -7 | -10.1 | 89 | + | T | A | T | 1 | 49,655,430 |
| mb14 | rs1898170 | -7.1 | -10.1 | 3138 | + | T | T | A | 4 | 29,351,338 |
| mb15 | rs11247921 | -7.1 | -10.3 | 126 | + | G | G | C | 1 | 26,665,602 |
| mb16 | rs1635718 | -7.2 | -10.5 | 311 | + | C | G | C | 1 | 35,974,469 |
| mb17 | rs10510620 | -7.2 | -11.2 | 373 | + | C | C | G | 3 | 29,330,286 |
| mb18 | rs7104025 | -7.1 | -10.4 | 649 | + | C | C | G | 11 | 21,148,373 |
| mb19 | rs2246745 | -7.3 | -10.8 | 1009 | + | T | T | A | 2 | 29,940,529 |
| mb22 | rs1884444 | -7.3 | -10.4 | 2518 | + | G | T | G | 1 | 67,633,812 |
| mb23 | rs2510152 | -6.8 | -10.5 | 165 | + | G | G | T | 11 | 119,148,573 |
| mb24 | rs16754 | -7.7 | -10.2 | 3297 | - | T | T | C | 11 | 32,417,945 |
| mb81 | rs6986290 | -7 | -10.08 | 39 | + | T | C | T | 8 | 28,071,119 |

Table S2-2: 20-plex mBDA used for NGS optimization experiments used for library protocol optimization. All toehold $\Delta G^\circ$ are approximately -7 kcal mol$^{-1}$, and all nonhomologous $\Delta G^\circ$ are approximately -10 kcal mol$^{-1}$.

| SNP locus | w/o B final RP Conc. /nM | w/B final RP Conc. /nM | SNP locus | w/o B final RP Conc. /nM | w/B final RP Conc. /nM |
|---|---|---|---|---|---|
| mb1 | 2.0 | 1.7 | mb41 | 2.7 | 1.4 |
| mb2 | 3.7 | 4.7 | mb42 | 13.6 | 15.6 |
| mb3 | 16.5 | 25.7 | mb43 | 2.5 | 1.1 |
| mb4 | 14.0 | 7.9 | mb44 | 4.0 | 3.7 |
| mb5 | 7.8 | 11.6 | mb45 | 11.2 | 4.1 |
| mb6 | 6.2 | 5.1 | mb46 | 56.9 | 33.4 |
| mb7 | 3.2 | 2.4 | mb47 | 19.0 | 3.3 |
| mb8 | 5.5 | 31.2 | mb48 | 30.6 | 37.4 |
| mb9 | 5.2 | 41.0 | mb49 | 24.8 | 1.0 |
| mb10 | 7.7 | 13.7 | mb50 | 13.8 | 44.1 |
| mb11 | 30.1 | 45.7 | mb51 | 3.5 | 13.4 |
| mb12 | 2.6 | 2.7 | mb52 | 9.9 | 17.4 |
| mb13 | 8.3 | 3.4 | mb53 | 1.8 | 2.2 |
| mb14 | 9.5 | 62.3 | mb54 | 126.1 | 62.3 |
| mb15 | 8.0 | 11.4 | mb55 | 12.9 | 5.2 |
| mb16 | 54.0 | 37.4 | mb56 | 4.9 | 0.5 |
| mb17 | 6.3 | 4.1 | mb57 | 12.3 | 1.5 |
| mb18 | 6.6 | 4.5 | mb58 | 3.3 | 38.9 |
| mb19 | 3.7 | 4.0 | mb59 | 30.5 | 2.8 |
| mb20 | 9.4 | 37.4 | mb60 | 3.6 | 6.1 |
| mb21 | 3.9 | 5.6 | mb61 | 4.4 | 2.3 |
| mb22 | 12.3 | 22.9 | mb62 | 4.3 | 1.8 |
| mb23 | 4.1 | 3.1 | mb63 | 70.0 | 37.4 |
| mb24 | 16.1 | 60.1 | mb64 | 126.1 | 37.4 |
| mb25 | 20.1 | 37.4 | mb65 | 1.4 | 0.5 |
| mb26 | 2.4 | 9.3 | mb66 | 4.7 | 8.0 |
| mb27 | 2.9 | 1.4 | mb67 | 25.2 | 37.4 |
| mb28 | 3.0 | 2.5 | mb68 | 21.4 | 12.9 |
| mb29 | 21.8 | 24.0 | mb69 | 51.2 | 30.7 |
| mb30 | 27.7 | 15.3 | mb70 | 3.0 | 1.3 |
| mb31 | 3.3 | 2.5 | mb71 | 39.4 | 37.4 |
| mb32 | 9.8 | 2.4 | mb72 | 49.3 | 23.5 |
| mb33 | 15.6 | 14.8 | mb73 | 21.2 | 0.7 |
| mb34 | 2.2 | 9.1 | mb74 | 2.9 | 31.3 |
| mb35 | 1.7 | 2.4 | mb75 | 5.0 | 2.6 |
| mb36 | 4.4 | 16.5 | mb76 | 17.1 | 10.8 |
| mb37 | 6.7 | 5.7 | mb77 | 5.1 | 10.8 |
| mb38 | 3.0 | 5.0 | mb78 | 7.0 | 3.7 |
| mb39 | 2.0 | 2.3 | mb79 | 4.6 | 3.1 |
| mb40 | 7.6 | 8.8 | mb80 | 3.4 | 5.1 |

Table S2-3: Optimized reverse primer (RP) concentrations for the 80-plex mBDA. For all amplicons, the forward primer concentrations were all 15 nM and the Blocker concentrations were all 150 nM.

#### Section S3: NGS Data Analysis

All NGS experiments in this manuscript were performed as single-end sequencing runs on an Illlumina MiSeq platform. After standard library demultiplexing, we first ran Bowtie2 using the 80 amplicon sequences as reference sequences to identify and remove the grossly off-target NGS reads (due to primer/adapter dimers and nonspecific PCR amplification of other regions of the human genome).

Subsequently, the on-target reads are compared against 3×80 reference DNA sequences, with 3 reference sequences constructed from the (+) strand of each amplicon, corresponding to the variant allele, the intended allele, and the downstream region (Fig. S3-1). Each on-target NGS read is compared to the variant allele, intended allele, and downstream region sequences for all amplicons, and then classified as either a variant read, an intended read, a read with an error, or "other" (Fig. S4-2). The variant reads and the intended reads are used for evaluating the variant read fraction (VRF), which is used for quantitating the initial VAF.

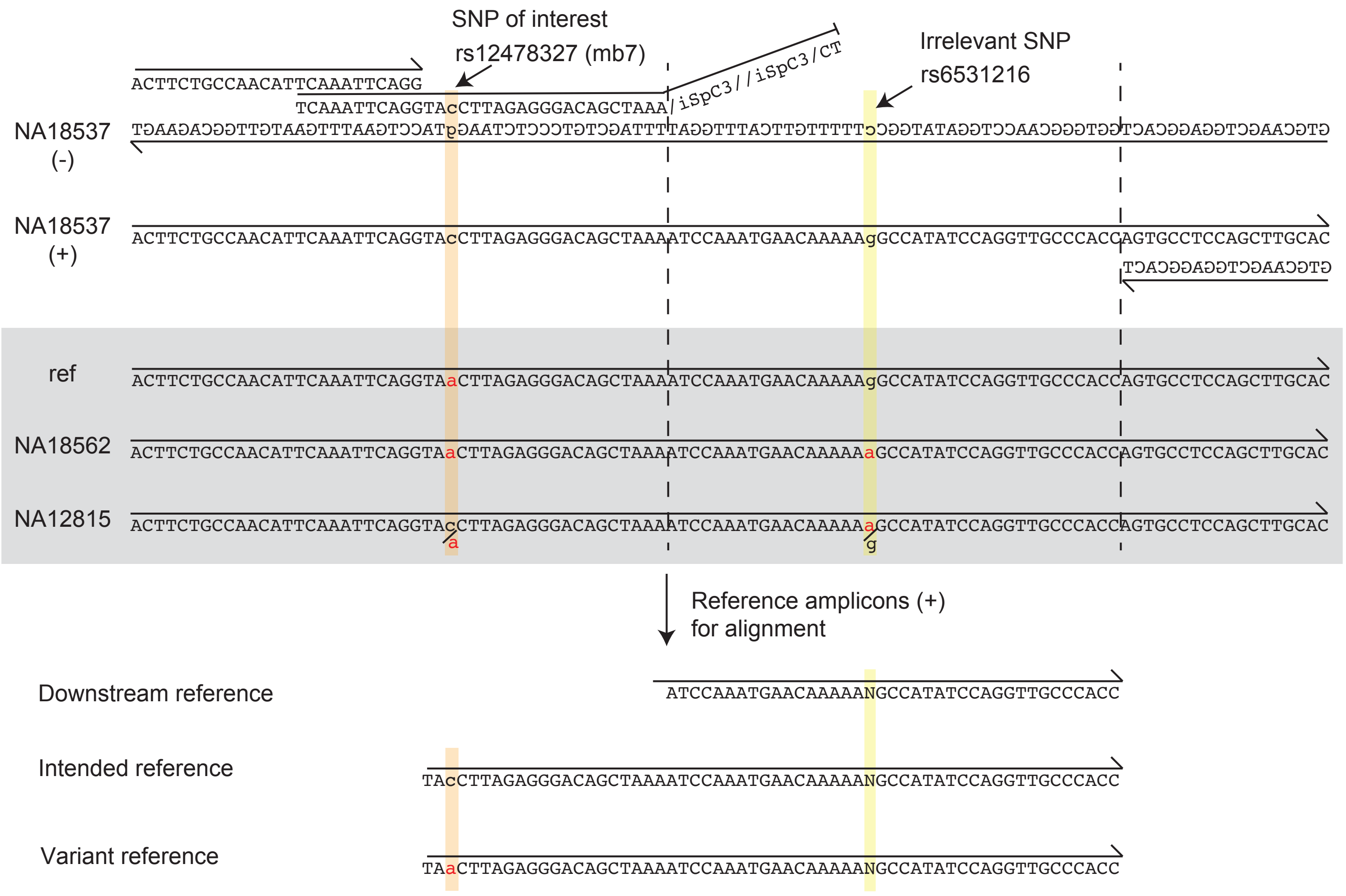


FIG. S3-1: Reference sequences for our NGS bioinformatics, including consideration for incidental SNPs in the amplicon region. Incidental SNPs are listed as wildcard symbols in the "grep" exact string match to determine the number of reads matched to the downstream reference, the intended reference, and the variant reference.

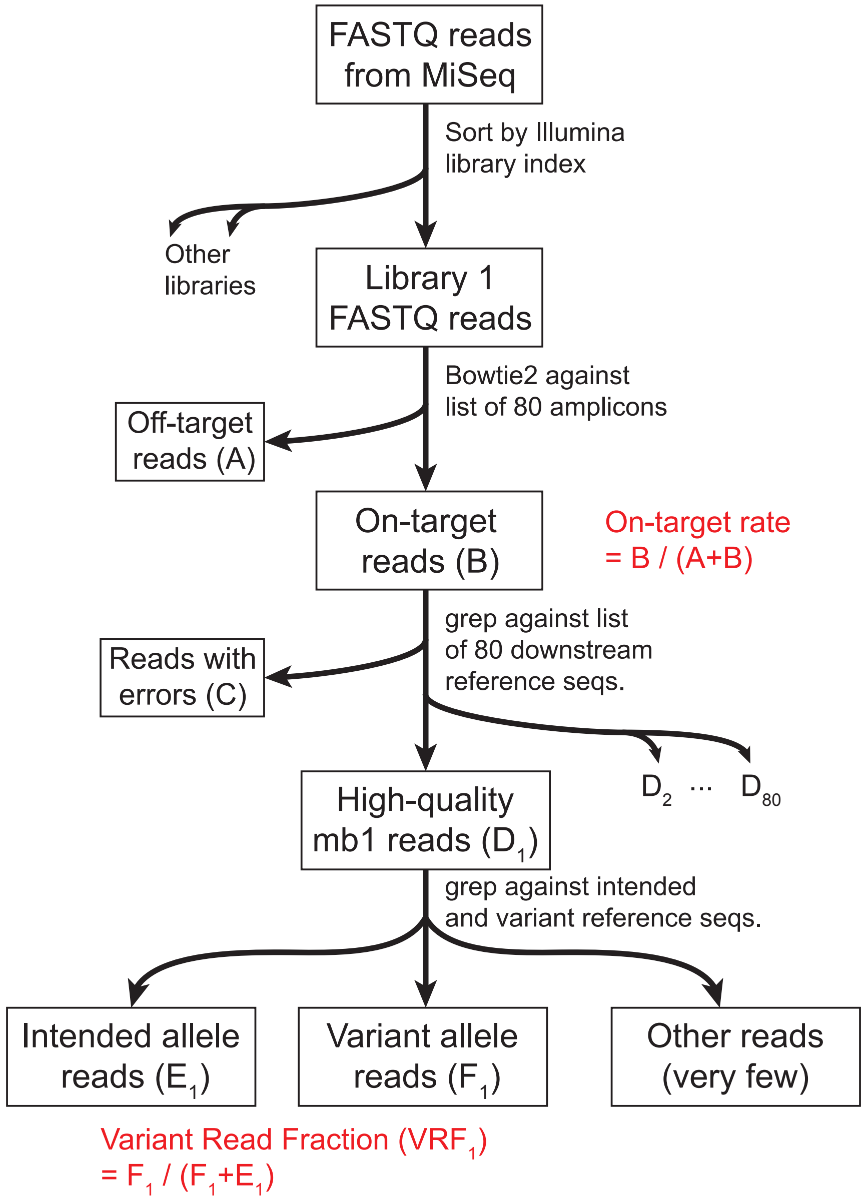


FIG. S3-2: Overall bioinformatics workflow for NGS FASTQ data processing.

**Bioinformatics comparison.** In this work, we developed our own bioinformatics pipeline, which included both a more stringent aligner and a direct counting method for comparing variant reads vs. wildtype reads for calculated variant read frequency (VRF). Our aligner can be compared against the popular BWA-mem and Bowtie 2 aligners (Table S3-1), and produces marginally fewer on-target reads by filtering away more suspect reads.

Our variant call software, compared against the standard Picard+GATK bioinformatics suite (Fig. S3-3), produces more discordance (Table S3-2). In a 16-amplicon mBDA NGS panel, using a reference sample with 15 spike-in variant sequences, we observed that Picard+GATK produces two false positive variant calls and two false negative variant calls. We manually went through our Fastq files to confirm that the Picard+GATK errors were in fact errors; the false positive calls did not have any supporting reads, and the false negative calls had many supporting reads.

For the two false negatives, a search of online forums showed that Picard+GATK has a well-known bug where it would fail to report variants in amplicon sequencing in amplicons with significantly higher read depth than mean read depth. This was the case for Seq9 and Seq10, which respectively had 23,387 reads and 13,919 reads compared to a mean read depth of 3,962. When we down-sampled the reads aligned to Seq9 and Seq10 by 10-fold, the expected variants were properly called. We are not sure the reason for Picard+GATK's false positives for non-present variants in Seq2 and Seq6, as no reads matched the called variants in the Fastq files.

We wish to emphasize that we do believe that our variant call software is generally better than Picard+GATK, only that it was designed with the mBDA NGS library preparation characteristics in mind.

| | On-target rate | On-target reads |
|---|---|---|
| Our method | 92.2% | 63,388 |
| BWA-mem | 93.9% | 64,572 |
| Bowtie 2 | 94.0% | 64,592 |

TABLE S3-1: Comparison of on-target rates and reads for different aligners. Here, we used a stringent method that discards marginally more NGS reads than conventional BWA and Bowtie 2 aligners.

| | Our VRF | Picard+GATK VRF |
|---|---|---|
| Seq1 | 10% | 10% |
| Seq2 | 0% | 10% |
| Seq3 | 75% | 77% |
| Seq4 | 83% | 94% |
| Seq5 | 62% | 68% |
| Seq6 | 25% | 25% |
| Seq6 (extra) | 0% | 35% |
| Seq7 | 61% | 78% |
| Seq8 | 73% | 66% |
| Seq9 | 8% | 0% |
| Seq10 | 6% | 0% |
| Seq11 | 96% | 93% |
| Seq12 | 34% | 39% |
| Seq13 | 23% | 27% |
| Seq14 | 98% | 96% |
| Seq15 | 89% | 93% |
| Seq16 | 17% | 10% |

TABLE S3-2: Comparison of VRF values from our pipeline vs. the Picard+GATK pipeline. False negatives from the Picard+GATK pipeline are displayed in blue, and false positives are displayed in red. Manual examination of the Fastq files confirmed that our pipeline is correct. The false negatives are due to a well-known bug in the Picard software that occurs when some amplicons are sequenced to significantly higher depth than other amplicons. False positive reads were not found in the Fastq file, and we are not sure why Picard+GATK would make this mistake.

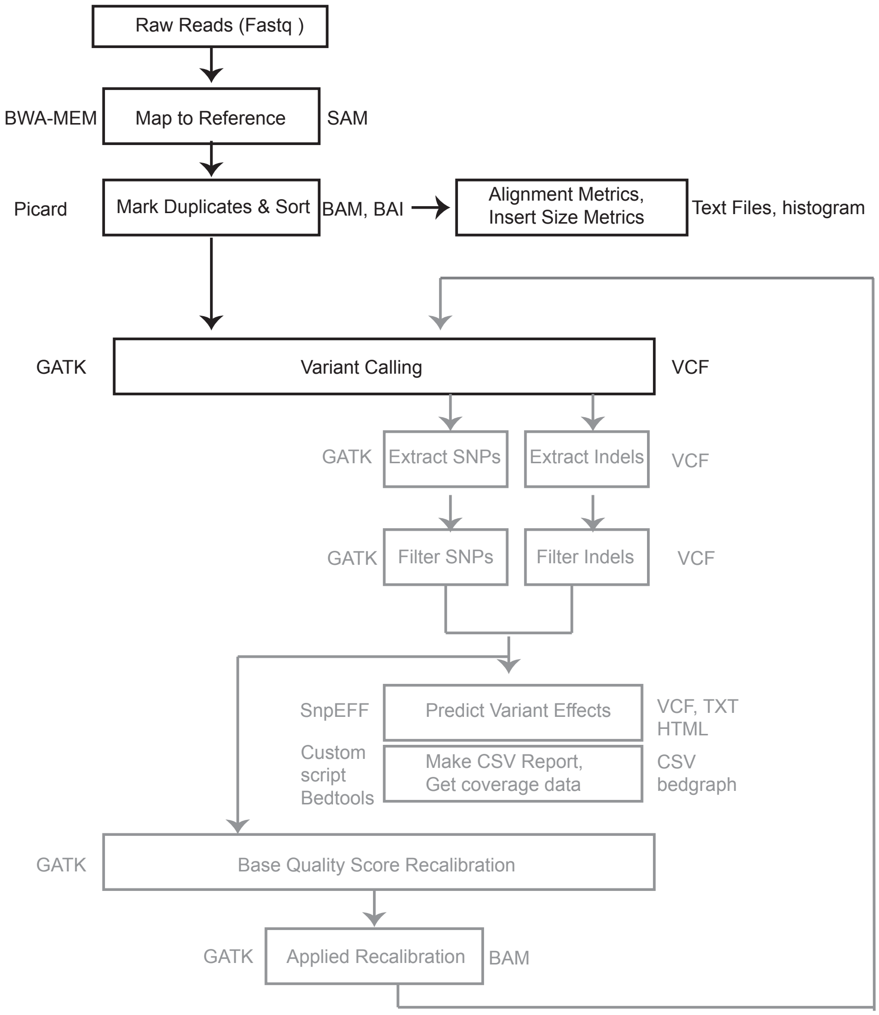


FIG. S3-3: Broad pipeline for variant calling. Here, the grayed out downstream steps not performed because we do not need to predict the pathogenicity nature of mutations.

#### Section S4: Measurement of Enrichment-Fold (EF) Values

The 80-plex mBDA primers/Blockers each have a different EF value, due to slight differences in the thermodynamics of both the mismatch bubble formed between the Blocker and the variant allele template, and due to differences in the secondary structure adopted by different templates/amplicons. However, for a given mBDA panel and protocol, EF is highly reproducible across different quantities and VAFs of input DNA. Here, we show experimental mBDA NGS results on the 80-plex mBDA NGS panel with varying input VAFs. Our experiments are use 0.03%, 0.05%, 0.1%, 0.2%, 0.5%, 1%, 2%, 3% mixtures of the NA18562 and NA18537 genomic DNA samples. Fig. S4-1 and S4-2 shows the nominal VAF and experimental VRF values for each of the 80 SNP variant alleles; these results are summarized in Fig. 1cd of the manuscript. Numerical values for the median EF are displayed in Table S4-1, and the median EF for all 80-plex is 409. Fig. S4-3 shows the corresponding calculated EF values based on VRF and VAF values for 3 SNP loci.

The reproducibility of inferred VAF is significantly impacted by the Poisson distribution statistics of sample preparation and dilution. In each of these experiments, 50 ng of DNA input was used, corresponding to roughly 15,000 haploid genomic copies. At VAF of less than 1%, stochastic variations in the number of molecules due to Poisson distribution can be quite significant. Recall that the variance of a Poisson distribution is equal to its mean. For 0.1% VAF in 50 ng which corresponds to an expected 15 DNA molecules with the variant allele, the standard deviation is 3.87 molecules. Thus, the simple act of preparing the 0.1% VAF sample will have a coefficient of variation of 25.8%. Fig. S4-4 shows that the observed distribution of VAFs is generally consistent with our expectations from the Poisson distribution.

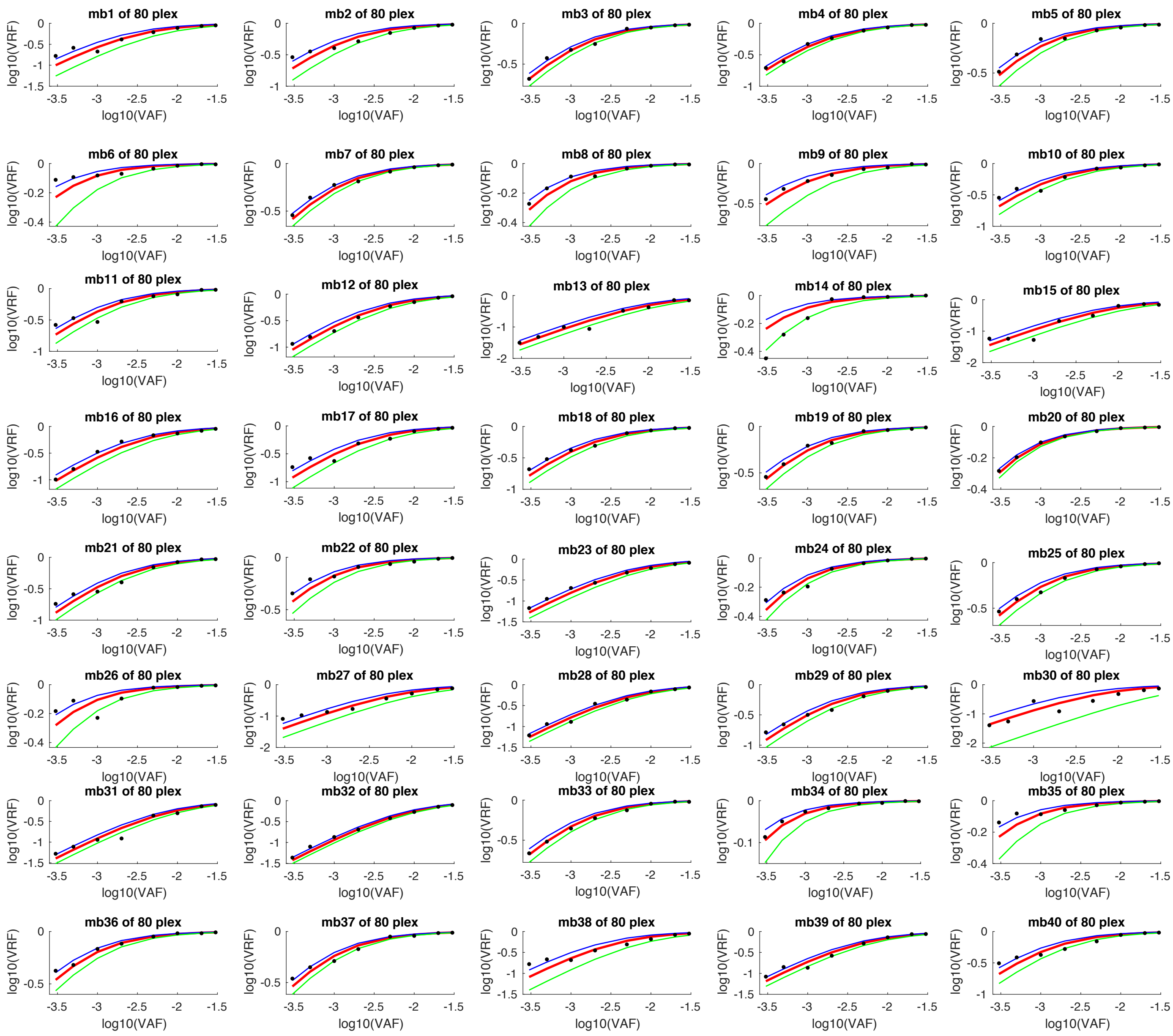

FIG. S4-1: Measured EF values for the 80-plex mBDA NGS panel, part 1 (mb1 to mb40). Red lines indicate the mean of eight different VAF values, and the blue and green lines show ± 1 standard deviation. Black dots indicate observed EF values.

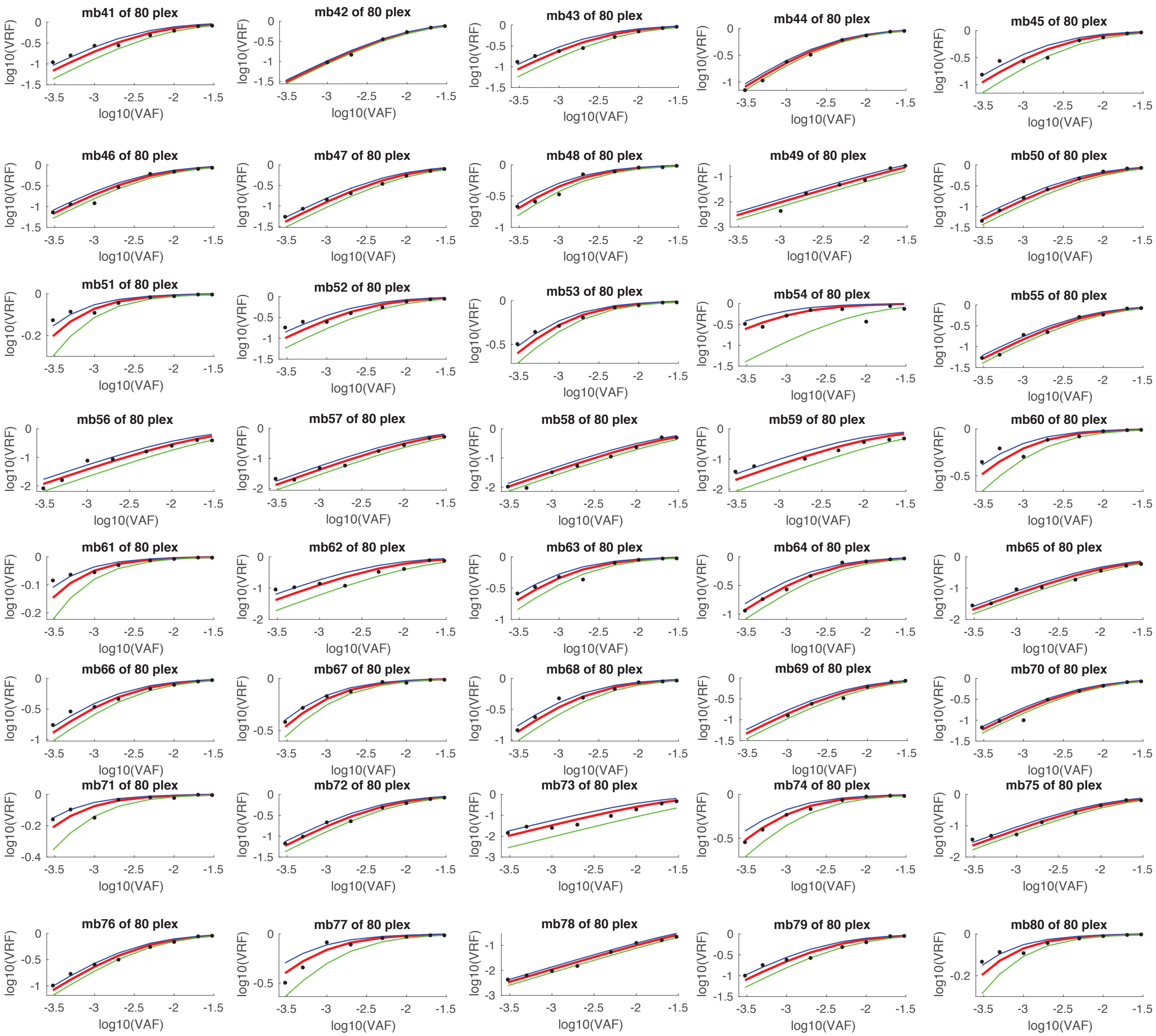


FIG. S4-2: Measured EF values for the 80-plex mBDA NGS panel, part 2 (mb41 to mb80). Red lines indicate the mean of eight different VAF values, and the blue and green lines show ± 1 standard deviation. Black dots indicate observed EF values.

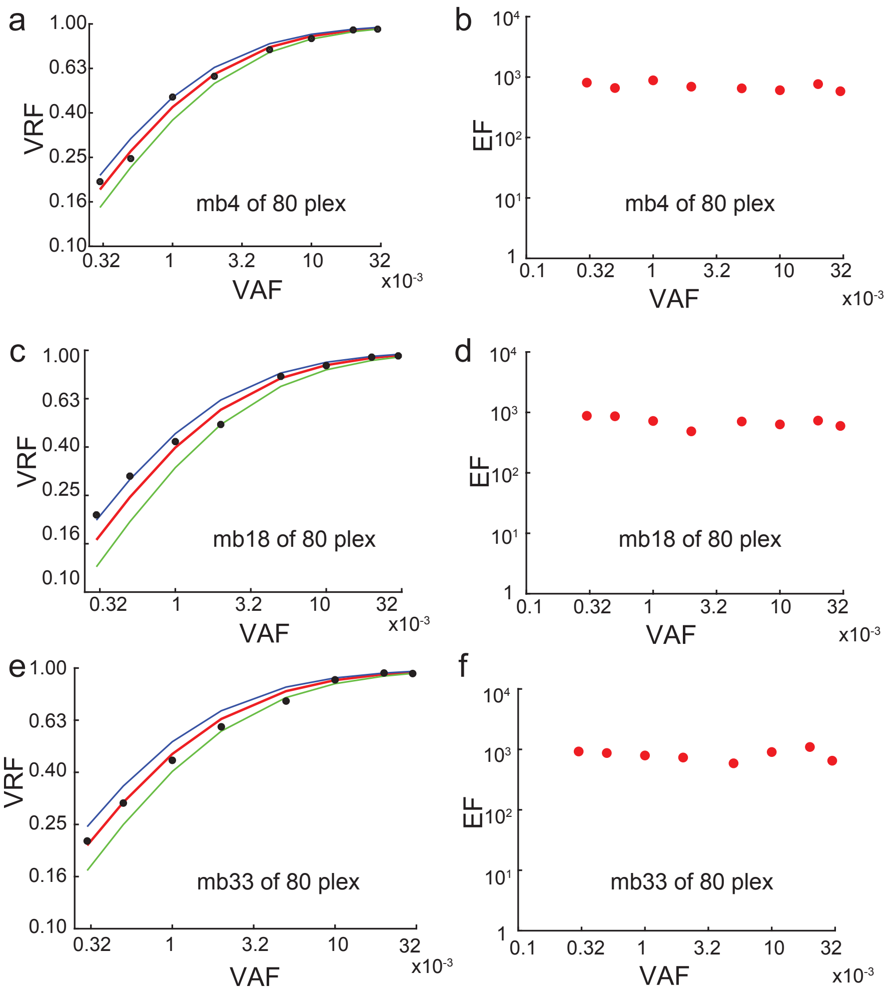


FIG. S4-3: Three example BDA systems in the 80-plex panel. The inferred EF values appear to be similar for all VAF values.

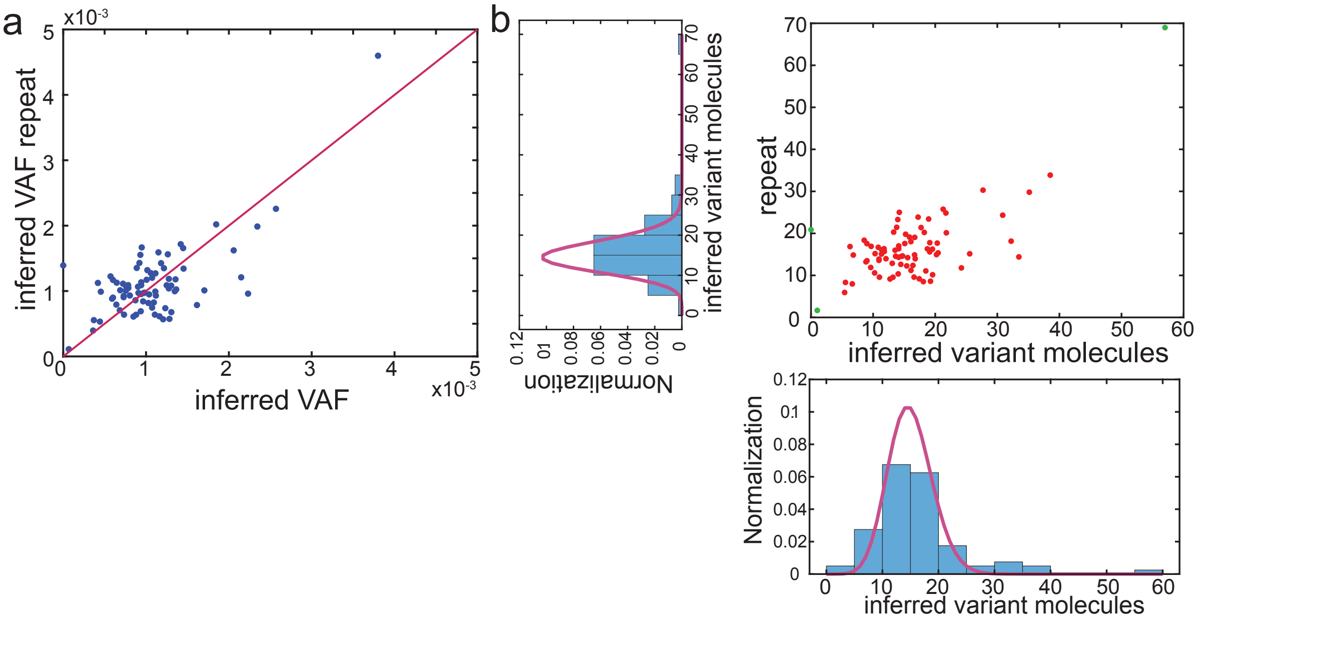


FIG. S4-4: Reproducibility of 80-plex mBDA NGS VAF inference **(a)** Experimental results from replicate NGS libraries using 50 ng human DNA input. Here, the input was a 0.1%:99.9% mixture of NA18562 and NA18537, and the 80 SNPs were selected to be homozygous for different alleles in NA18562 and NA18537, so all SNPs should be at roughly 0.1% VAF, subject to Poisson distribution variation. **(b)** Inferred number of DNA molecules with variant alleles. The purple traces overlaid on the histograms on the horizontal and vertical axes show the expected distribution of variant allele molecules based on Poisson distribution.

| BDA index | SNP rs# | $\Delta G^\circ_P$ | $\Delta G^\circ_B$ | EF | BDA index | SNP rs# | $\Delta G^\circ_P$ | $\Delta G^\circ_B$ | EF |
|---|---|---|---|---|---|---|---|---|---|
| mb1 | rs10230708 | -7.3 | -10.5 | 316 | mb41 | rs12259813 | -7.5 | -10.4 | 199 |
| mb2 | rs10104396 | -7.5 | -11 | 670 | mb42 | rs12541300 | -6.8 | -9.8 | 278 |
| mb3 | rs199032 | -7 | -10 | 870 | mb43 | rs12681931 | -7.4 | -10.2 | 307 |
| mb4 | rs926850 | -7.3 | -10.4 | 690 | mb44 | rs12782580 | -7 | -10.1 | 297 |
| mb5 | rs17149369 | -7.1 | -10.2 | 1195 | mb45 | rs1375977 | -6.6 | -9.8 | 379 |
| mb6 | rs869720 | -7.1 | -10.5 | 4422 | mb46 | rs1516755 | -6.7 | -10 | 252 |
| mb7 | rs12478327 | -7 | -10.5 | 1107 | mb47 | rs1524303 | -7 | -10.3 | 127 |
| mb8 | rs2638145 | -7.3 | -10.7 | 2702 | mb48 | rs1667087 | -7.4 | -10.8 | 726 |
| mb9 | rs2170091 | -7.4 | -10.7 | 1275 | mb49 | rs16871316 | -7.7 | -10.5 | 51 |
| mb10 | rs2043583 | -7.2 | -10.2 | 919 | mb50 | rs16925478 | -7.3 | -10.6 | 174 |
| mb11 | rs955456 | -6.9 | -10.4 | 707 | mb51 | rs17560702 | -7 | -10 | 4647 |
| mb12 | rs966516 | -7.1 | -10.2 | 294 | mb52 | rs1937037 | -6.5 | -10 | 319 |
| mb13 | rs354169 | -7 | -10.1 | 97 | mb53 | rs2215492 | -6.9 | -9.9 | 1030 |
| mb14 | rs1898170 | -7.1 | -10.1 | 4356 | mb54 | rs2301720 | -7.2 | -10 | 872 |
| mb15 | rs11247921 | -7.1 | -10.3 | 129 | mb55 | rs2616187 | -7.7 | -10.8 | 168 |
| mb16 | rs1635718 | -7.2 | -10.5 | 375 | mb56 | rs2710998 | -6.6 | -9.4 | 33 |
| mb17 | rs10510620 | -7.2 | -11.2 | 396 | mb57 | rs2807238 | -6.9 | -10.3 | 40 |
| mb18 | rs7104025 | -7.1 | -10.4 | 669 | mb58 | rs2874755 | -7.3 | -10.4 | 32 |
| mb19 | rs2246745 | -7.3 | -10.8 | 1231 | mb59 | rs3813787 | -7.3 | -10.1 | 57 |
| mb20 | rs3789806 | -6.4 | -9.6 | 3421 | mb60 | rs4665582 | -6.7 | -10 | 1454 |
| mb21 | rs706714 | -7.8 | -10.4 | 474 | mb61 | rs4712476 | -7.4 | -10.8 | 7146 |
| mb22 | rs1884444 | -7.3 | -10.4 | 1980 | mb62 | rs611628 | -7.7 | -10.6 | 108 |
| mb23 | rs2510152 | -6.8 | -10.5 | 179 | mb63 | rs6452035 | -7.3 | -10.1 | 841 |
| mb24 | rs16754 | -7.7 | -10.2 | 2666 | mb64 | rs6816854 | -7.4 | -10.3 | 446 |
| mb25 | rs206781 | -7.1 | -10.5 | 1110 | mb65 | rs6937778 | -7.4 | -10.5 | 66 |
| mb26 | rs28932178 | -6.6 | -10.1 | 3227 | mb66 | rs7003044 | -6.6 | -9.5 | 438 |
| mb27 | rs10186821 | -7.4 | -10.1 | 107 | mb67 | rs7032336 | -7.4 | -10.8 | 1569 |
| mb28 | rs10508599 | -7.6 | -10.8 | 191 | mb68 | rs7816009 | -6.9 | -9.8 | 474 |
| mb29 | rs10738578 | -7.2 | -10.2 | 463 | mb69 | rs7893462 | -7.6 | -10.6 | 158 |
| mb30 | rs10741037 | -6.6 | -9.7 | 92 | mb70 | rs7902135 | -7.6 | -10.6 | 217 |
| mb31 | rs10770674 | -7.4 | -10.4 | 135 | mb71 | rs898476 | -7.6 | -10.6 | 4603 |
| mb32 | rs10805227 | -7 | -10.4 | 123 | mb72 | rs9368431 | -6.8 | -10 | 188 |
| mb33 | rs10833604 | -6.8 | -9.9 | 870 | mb73 | rs9438621 | -6.6 | -9.3 | 25 |
| mb34 | rs10964389 | -7.4 | -10.6 | 11929 | mb74 | rs9466035 | -7.6 | -10.6 | 1324 |
| mb35 | rs11015816 | -7.3 | -10.7 | 3590 | mb75 | rs9466930 | -6.7 | -9.4 | 74 |
| mb36 | rs11045749 | -7.5 | -10.6 | 1659 | mb76 | rs9973865 | -7.1 | -10.2 | 293 |
| mb37 | rs1123828 | -7.2 | -9.9 | 1352 | mb77 | rs4712498 | -7.3 | -10 | 1675 |
| mb38 | rs11708584 | -6.9 | -9.6 | 244 | mb78 | rs2073149 | -7.3 | -10.1 | 11 |
| mb39 | rs12192635 | -6.9 | -10.3 | 213 | mb79 | rs2862909 | -6.9 | -9.9 | 283 |
| mb40 | rs12213948 | -7.1 | -10.2 | 812 | mb80 | rs1338945 | -7.4 | -10.3 | 4735 |

TABLE S4-1: Experimentally observed EF values for each of the BDA systems in the 80-plex mBDA panel. EF values listed in this table are median values of those 8 different VAF values. Also displayed are the computed $\Delta G^\circ_P$ and $\Delta G^\circ_B$ values for each BDA system.

#### Section S5: NGS Depth and Library Size Needed

**Depth Needed for VAF Convergence.** To analyze the minimum sequencing depth needed for reliable detection of SNP allele variants at the 0.1% to 0.2% VAF range, we consider the convergence of inferred VAF values for each SNP locus (Fig. 5e of main text). For each amplicon, we model the variant and intended allele reads as a binary distribution around a true probability $p$ that a randomly selected on-target and error-free NGS read will have the variant allele. The binary distribution has a standard deviation of $\sqrt{N \cdot p \cdot (1-p)}$, and the sample mean converges rapidly to the true mean at $N > 300$ when $p \in [0.05, 0.95]$.

For each amplicon of the 80-plex mBDA panel, we count up the number of exact NGS read matches to the variant amplicon and the intended amplicon (Supplementary Section S5). These reads are represented as a binary vector of 1's and 0's, with 1's representing variant alleles and 0's representing intended alleles. The binary vectors are then randomly permuted, in order to remove any potential bias in reporting order by the Illumina software.

We then iteratively calculated the VRF values considering the first $D$ elements of the permuted binary vector, with $D$ varying between 50 and 5,000. These VRF values are then used to calculated the VAF value using the median EF values reported in Fig. 2c and Supplementary Section S7. The convergence of inferred VAF values plotted in Fig. 3e were similar for multiple different random permutations of the binary vectors, indicating the 250x to 500x depth is sufficient to reproducible VAF quantitation down to 0.03% (data not shown). Simulations were programmed in Matlab and are available upon request.

The sufficiency of 250x sequencing depth for reproducible quantitation of VAF is dependent on both EF values being sufficiently large and VAF being sufficiently high to avoid Poisson distribution error. When $p = 0.05$ and EF = 50, then quantitated VAF is approximately 0.1%. With $N = 300$ reads, the expected number of variant reads is 15, with a standard deviation of 3.87 reads. This is compounded with the Poisson distribution, to create a total coefficient of variation of 36.5% for an independent replicate experiment from DNA sample. Large EF values reduce the error based on sampling from a binary distribution, and large input DNA quantities reduce the error due to Poisson distribution.

**Simulations of Small NGS Libraries.** Next, we sampled subsets of the FASTQ file corresponding to the 0.18% D9 (Hela) contaminant in NA18537 (shown in manuscript Fig. 3a), in order to determine the number of library reads needed to accurately determine VAFs for all 80 amplicons in the mBDA panel. We randomly permuted all 358,279 on-target reads for the library, and computed the VRFs and implied VAFs using samples of 5k (5000), 7k, 10k, 13k, 20k, 30k, 50k, 70k, and 100k reads (Fig. S5-1). Consistent with our expectations, the correlation constant $R^2$ between the VAFs of the full library and the reads sample increased as the number of reads in the sample increased. For statistical replicability, we repeated the above analysis for 15 random permutations of the NGS reads, and the summary of the $R^2$ values are shown in manuscript Fig. 5g.

At 50,000 reads, $R^2$ values reached above 99%, indicating sufficient reads for accurate VAF determination. Considering that the library on-target rates observed were in excess of 80%, rough 65,000 total reads are needed for an mBDA library to achieve a limit of detection of 0.1% VAF. This library size is roughly a factor of 100 lower than what is needed by deep NGS using UMIs.

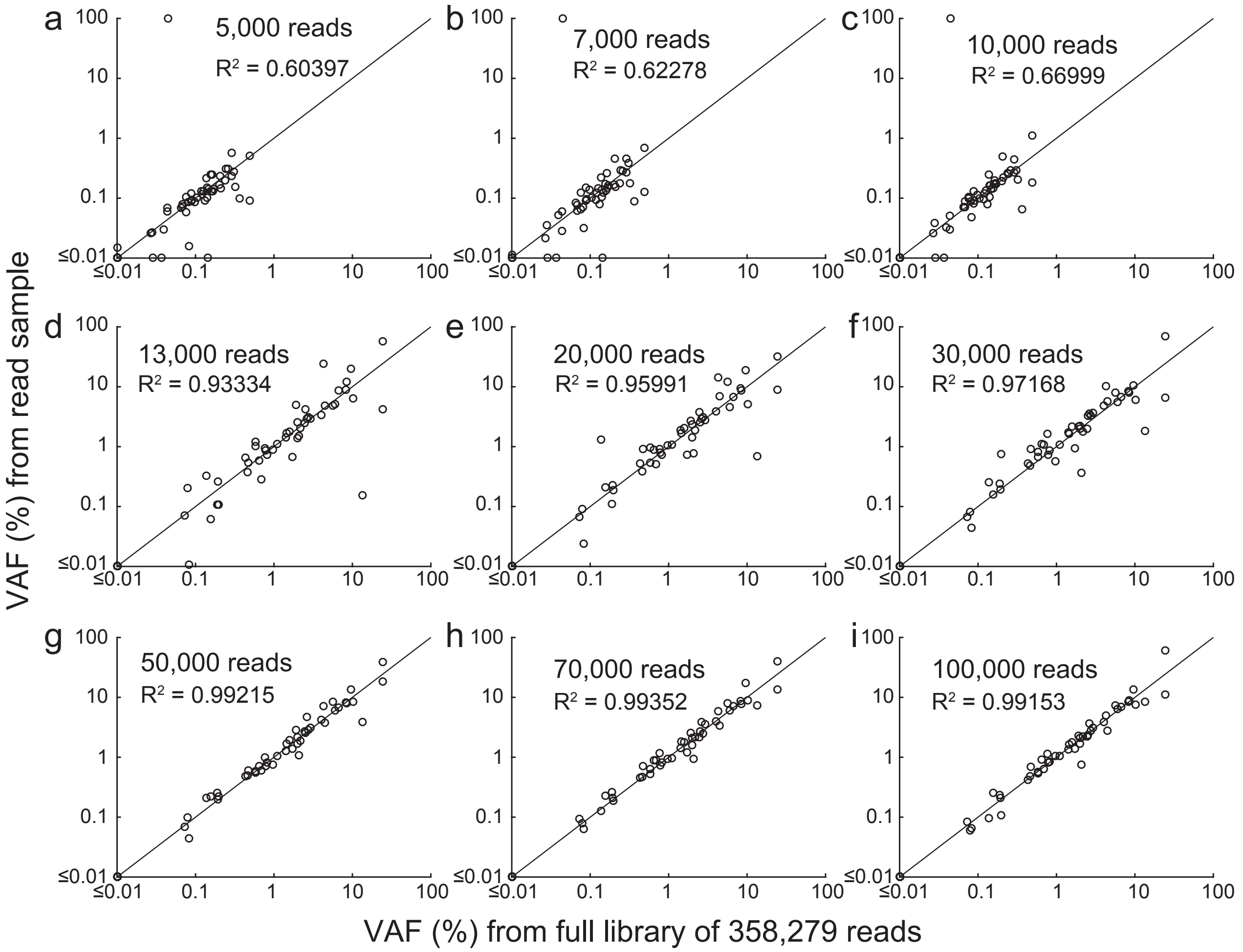


FIG. S5-1. Example results of sampling a subset of NGS on-target reads for the 0.18% D9 (HeLa) contaminant library. Panels (a) through (i) each show the correlation between inferred VAF for the reads sample vs. the entire library, with each circle corresponding to one of the 80 amplicons. The $R^2$ correlation constant is above 99% when over 50,000 on-target reads are analyzed.

#### Section S6: Variant Allele Detection with Small Input DNA Quantities

We performed mBDA NGS analysis of the same 0.18% HeLa (D9) contaminant in NA18537, but using 10 ng of input DNA, as compared to 300 ng used for libraries in Fig. 3a. For cell line contamination detection, 300 ng or more genomic DNA is easily available from DNA extraction. The purpose of this test is to show the applicability of mBDA NGS technology to other applications, such as analysis of cell-free DNA from peripheral blood plasma or tissue biopsy DNA from fine-needle aspirates.

Our results in Fig. S6-1 show that the specificity and sensitivity of variant allele calls are both reduced compared to 300 ng input DNA. The mean sensitivity of the two replicates on 10 ng, assuming a variant call threshold of 0.019% inferred VAF as in the main text, is 88.3%; the mean specificity is 83.3%.

At 10 ng input, homozygous variant alleles at 0.18% VAF correspond to an expected 5.4 molecules, and heterozygous variant alleles correspond to 2.7 molecules. Consequently, the loss of sensitivity is in line with expectation, given Poisson distribution on the actual number of variant allele DNA molecules in the input.

The loss of specificity can be attributed to (1) sample contamination and (2) increased DNA polymerase errors. Although we tried our best to minimize contamination of DNA samples by amplicons from previous experiments, via the separation of different NGS library preparation steps into different enclosed rooms, it is impossible to absolutely eliminate contamination. For 300 ng of input, a single molecule of DNA bearing the variant allele would contribute 0.001% VAF and thus is below our variant call floor. In contrast, with 10 ng of input DNA, a single molecule of DNA variant contaminant contributes 0.03% VAF, and would result in a false positive.

At 10 ng of input, we required 5 more cycles of PCR to reach library concentrations required by Illumina sequencing. Rather than increasing the number of mBDA cycles that could result in potential changes to EF values, we instead increased the number of index PCR cycles by 5. These 5 additional index PCR cycles present additional opportunities for the DNA polymerase to make misincorporation errors.

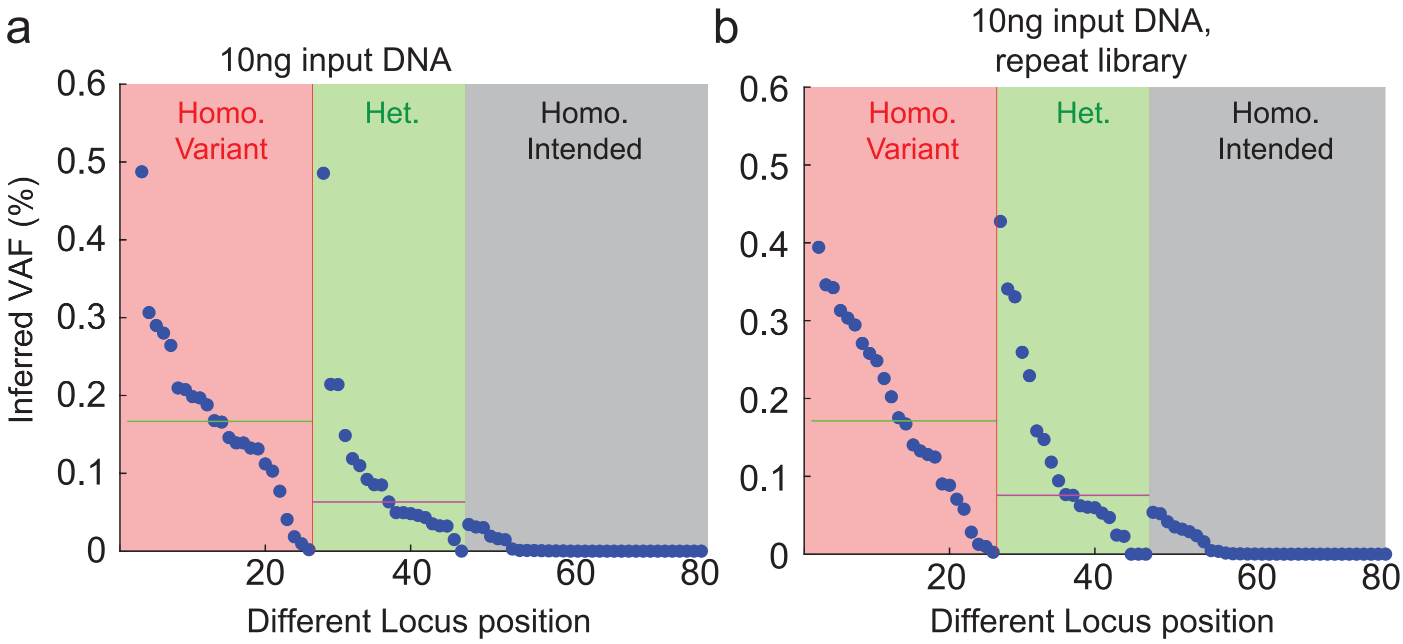


FIG. S6-1. Experimental results for the 80-plex mBDA NGS applied to 10 ng of input DNA (0.18% HeLa in NA18537).

#### Section S7: mBDA qPCR Optimization

**mBDA qPCR plex number and performance.** The closed-tube nature of qPCR reactions, while minimizing contamination risk, precludes Ampure Beads size selection steps that mitigate the impact of primer dimers and non-specific genomic amplification. To assess the multiplexing limit of mBDA in qPCR settings, we constructed and experimentally tested a number of mBDA qPCR assays of different plex numbers. These results are summarized in Fig. S7-1. At 21-plex and above, ΔCt are lower than 10-plex and below. However, 10-plex is not sufficient to guarantee that an arbitrary human cell line contaminant will possess at least 1 variant allele. Consequently, we proceeded with the 21-plex mBDA qPCR assay for cell line contamination detection demonstrations.

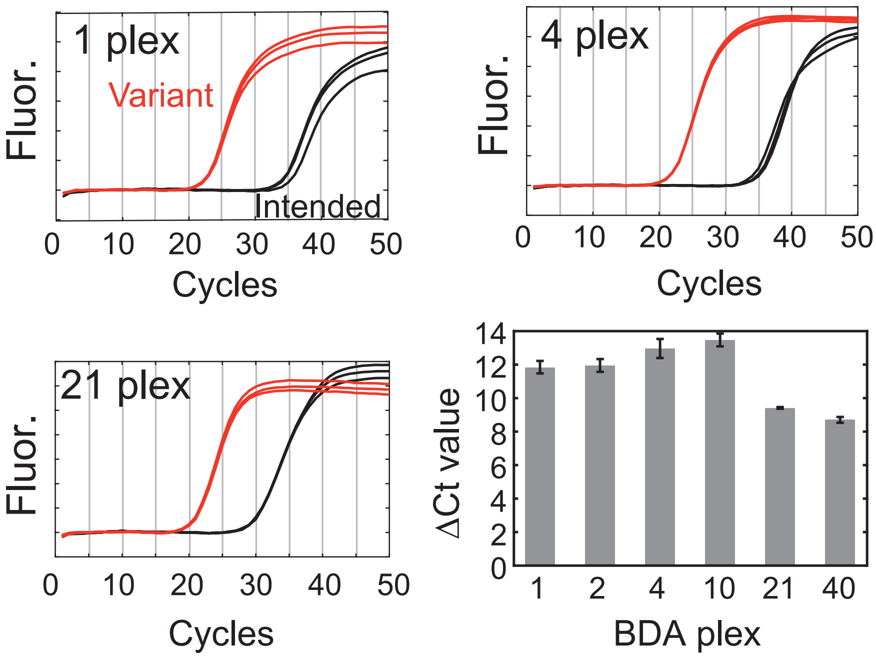


Figure S7-1: Observed difference in Ct values for different plex number mBDA qPCR assays. The intended sample here is the NA18537 human gDNA, and the variant is NA18562. For the SNP loci included in the mBDA qPCR assays, NA18562 is homozygous variant for all SNPs. The ΔCt quantity approximates the VAF limit of detection of the mBDA qPCR assay (LoD $\approx 2^{-\Delta \mathrm{Ct}}$).

**PCR thermocycling protocol optimization.** For mBDA qPCR assays, we reduced the concentrations of each primer and Blocker relative to single-plex BDA assay, because of the increased number of species that increase the total concentration of DNA in the PCR mixture. Literature reports and our own experience suggests that high concentrations DNA will inhibit PCR. Because the purpose of the anneal step of PCR thermocycling is to provide primers with sufficient time to hybridize to templates/amplicons, and we expected that we would need to increase the duration of the anneal steps for mBDA.

For PCR protocol optimization experiments, we performed single-plex analysis on 3 BDA designs (mb4, mb11, and mb16). The Blockers were designed to block NA18537's homozygous alleles (A for mb4, T for mb11, and G for mb16); NA18562's homozygous alleles (G for mb4, G for mb11, and C for mb16) are thus variant alleles that would be efficiently amplified. To ensure that the results would be applicable to NGS mBDA protocols as well, we used the Phusion Hot Start Flex enzyme mix (New England Biolabs). All experiments here used 20 ng of input genomic DNA per reaction.

Tables S7-1, S7-2, and S7-3 show qPCR results for different PCR thermocycling protocols, for single-plex reactions with 15 nM of each primer. Only at the longest PCR thermocycling protocol (5 min anneal time and 2 min extension time) did we observe Ct values similar to those under standard qPCR conditions (400 nM primers). Similarly, we also tested those three single-plex BDA reaction using 50 nM primer concentrations (Table S7-4, S7-5, and S7-6).

The single-plex BDA qPCR Ct values for the 21 SNPs used in Figs. 1 and 2 in the main text are shown in Table S7-7, using the 15 nM primer concentrations used for mBDA qPCR assays. The observed ΔCt (between the intended and variant allele templates) values imply a VAF limit of detection of $2^{-\Delta \mathrm{Ct}}$. For comparison, Ct and ΔCt values for single-plex BDA assays using standard 400 nM primer concentrations are shown in Table S7-8. Single-plex BDA results for the other set of SNP alleles are shown in Table S7-9.

(a)

| | Time | Temp (°C) |
|---|---|---|
| Initial Denaturation (sec) | 30 | 98 |
| Denaturation (sec) | 10 | 98 |
| Annealing (min) | 0.5 | 63 |
| Extension (min) | 0.5 | 72 |
| Final Extension (min) | 5 | 72 |

(b)

| | w/o B | | w/B | | | | |
|---|---|---|---|---|---|---|---|
| Phusion HiFi | 18537(WT) | 18562(SNP) | NTC | 18537(WT) | 18562(SNP) | ΔCt | Delay |
| mb4 | >40 | >40 | >40 | >40 | >40 | NaN | NaN |
| mb11 | >40 | >40 | >40 | >40 | >40 | NaN | NaN |
| mb16 | >40 | >40 | >40 | >40 | >40 | NaN | NaN |

Table S7-1: 30 s anneal and extension cycles fail to amplify at 15 nM primer concentrations.

(a)

| | Time | Temp( °C) |
|---|---|---|
| Initial Denaturation (sec) | 30 | 98 |
| Denaturation (sec) | 10 | 98 |
| Annealing (min) | 1 | 63 |
| Extension (min) | 1 | 72 |
| Final Extension (min) | 5 | 72 |

(b)

| | w/o B | | w/B | | | | |
|---|---|---|---|---|---|---|---|
| Phusion HiFi | 18537(WT) | 18562(SNP) | NTC | 18537(WT) | 18562(SNP) | ΔCt | Delay |
| mb4 | >40 | >40 | >40 | >40 | >40 | NaN | NaN |
| mb11 | >40 | >40 | >40 | >40 | >40 | NaN | NaN |
| mb16 | >40 | >40 | >40 | >40 | >40 | NaN | NaN |

Table S7-2: 1 min anneal and extension cycles fail to amplify at 15 nM primer concentrations.

(a)

| | Time | Temp( °C) |
|---|---|---|
| Initial Denaturation (sec) | 30 | 98 |
| Denaturation (sec) | 10 | 98 |
| Annealing (min) | 5 | 63 |
| Extension (min) | 2 | 72 |
| Final Extension (min) | 5 | 72 |

(b)

| | w/o B | | w/B | | | |
|---|---|---|---|---|---|---|
| Phusion HiFi | 18537(WT) | 18562(SNP) | 18537(WT) | 18562(SNP) | ΔCt | Delay |
| mb4 | 18.6 | 20 | 35 | 20.2 | 14.8 | 0.2 |
| mb11 | 20.6 | 21 | 40 | 23 | 17 | 2 |
| mb16 | 19 | 20.6 | 33 | 24 | 9 | 3.4 |

Table S7-3: 5 min anneal and 2 min extension cycles allow 15 nM primer concentrations to amplify DNA templates with similar efficiency as standard qPCR conditions at 400 nM primers and 30 s anneal/extension cycles.

(a)

| | Time | Temp( °C) |
|---|---|---|
| Initial Denaturation (sec) | 30 | 98 |
| Denaturation (sec) | 10 | 98 |
| Annealing (min) | 0.5 | 63 |
| Extension (min) | 0.5 | 72 |
| Final Extension (min) | 5 | 72 |

(b)

| | w/o B | | w/B | | | |
|---|---|---|---|---|---|---|
| Phusion HiFi | 18537(WT) | 18562(SNP) | 18537(WT) | 18562(SNP) | ΔCt | Delay |
| mb4 | 30 | 29.5 | >40 | 34 | NaN | 4.4 |
| mb11 | 34 | 35.6 | >40 | 39.9 | NaN | 4.4 |
| mb16 | 32.5 | 33.3 | >40 | >40 | NaN | NaN |

Table S7-4: 30 s anneal and extension cycles cause many Ct delay to amplify at 50 nM primer concentrations.

(a)

| | Time | Temp( °C) |
|---|---|---|
| Initial Denaturation (sec) | 30 | 98 |
| Denaturation (sec) | 10 | 98 |
| Annealing (min) | 1 | 63 |
| Extension (min) | 1 | 72 |
| Final Extension (min) | 5 | 72 |

(b)

| | w/o B | | w/B | | | |
|---|---|---|---|---|---|---|
| Phusion HiFi | 18537(WT) | 18562(SNP) | 18537(WT) | 18562(SNP) | ΔCt | Delay |
| mb4 | 22.7 | 23.3 | >40 | 21 | NaN | -2.4 |
| mb11 | 26.8 | 25.7 | >40 | 28.9 | NaN | 3.2 |
| mb16 | 23.7 | 23.4 | >40 | 30.6 | NaN | 7.2 |

Table S7-5: 1 min anneal and extension cycles cause many Ct delay to amplify at 50 nM primer concentrations.

| (a) | | Time | Temp( °C) | (b) |
|---|---|---|---|---|
| | Initial Denaturation (sec) | 30 | 98 | |
| | Denaturation (sec) | 10 | 98 | |
| | Annealing (min) | 5 | 63 | |
| | Extension (min) | 2 | 72 | |
| | Final Extension (min) | 5 | 72 | |

| | w/o B | | w/B | | | |
|---|---|---|---|---|---|---|
| Phusion HiFi | 18537(WT) | 18562(SNP) | 18537(WT) | 18562(SNP) | ΔCt | Delay |
| mb4 | 18.6 | 19 | 32 | 18.9 | 13.1 | -0.1 |
| mb11 | 18.5 | 18.6 | 32.7 | 19.3 | 13.4 | 0.7 |
| mb16 | 17.7 | 18.3 | 32.4 | 20.2 | 12.2 | 1.9 |

Table S7-6: 5 min anneal and 2 min extension cycles allow 50 nM primer concentrations to amplify DNA templates with similar efficiency as standard qPCR conditions at 400 nM primers and 30 s anneal/extension cycles.

| SNP locus | NA18562 no Blocker | NA18537 no Blocker | NA18562 w/ Blocker | NA18537 w/ Blocker | ΔCt | Delay |
|---|---|---|---|---|---|---|
| mb1 | 22.0 | 22.2 | 22.8 | 35.8 | 13.0 | 0.8 |
| mb5 | 23.5 | 23.3 | 25.6 | 38.8 | 13.2 | 2.0 |
| mb9 | 21.6 | 21.6 | 22.3 | 33.4 | 11.1 | 0.6 |
| mb11 | 27.1 | 27.0 | 29.2 | 44.3 | 15.1 | 2.1 |
| mb18 | 24.1 | 24.2 | 24.3 | 38.9 | 14.6 | 0.2 |
| mb22 | 25.2 | 25.0 | 28.2 | 44.1 | 15.9 | 3.0 |
| mb27 | 22.1 | 22.3 | 23.0 | 35.8 | 12.9 | 0.8 |
| mb29 | 23.2 | 23.4 | 24.1 | 39.5 | 15.4 | 0.9 |
| mb34 | 22.6 | 22.5 | 27.1 | 42.0 | 14.9 | 4.5 |
| mb37 | 24.3 | 24.3 | 27.8 | 39.6 | 11.8 | 3.5 |
| mb40 | 23.8 | 23.8 | 26.4 | 40.3 | 13.9 | 2.5 |
| mb46 | 25.4 | 25.3 | 27.7 | 41.3 | 13.6 | 2.3 |
| mb53 | 23.4 | 24.0 | 23.7 | 42.1 | 18.4 | 0.3 |
| mb60 | 22.8 | 22.9 | 26.7 | 41.2 | 14.6 | 3.9 |
| mb62 | 23.0 | 22.7 | 26.2 | 37.1 | 10.9 | 3.1 |
| mb64 | 28.3 | 28.4 | 28.0 | 43.1 | 15.1 | -0.3 |
| mb68 | 26.2 | 26.3 | 28.1 | 42.1 | 14.0 | 1.9 |
| mb69 | 23.6 | 23.5 | 23.9 | 35.1 | 11.2 | 0.3 |
| mb73 | 23.2 | 23.3 | 24.4 | 40.1 | 15.7 | 1.2 |
| mb76 | 23.7 | 23.6 | 23.7 | 33.3 | 9.6 | 0.0 |
| mb78 | 26.8 | 26.9 | 28.8 | 41.8 | 13.0 | 2.1 |

Table S7-7: Single-plex BDA results for the 21 SNPs used as part of the mBDA qPCR panel for detection of conspecific DNA contaminants in NA18537. Here, we used the PowerUp SYBR Green Master Mix as PCR polymerase, and 15 nM of each primer. During thermocycling, the anneal and extension steps were combined as a 5 min step at 60 °C. 20 ng of genomic DNA was used as input for each reaction.

| SNP locus | NA18562 no Blocker | NA18537 no Blocker | NA18562 w/ Blocker | NA18537 w/ Blocker | ΔCt | Delay |
|---|---|---|---|---|---|---|
| mb1 | 25.1 | 25.1 | 25.2 | 39.6 | 14.4 | 0.2 |
| mb5 | 26.1 | 25.6 | 26.1 | 40.3 | 14.1 | 0.1 |
| mb9 | 24.9 | 25.0 | 27.1 | 43.0 | 15.9 | 2.2 |
| mb11 | 25.4 | 25.6 | 27.4 | 42.7 | 15.4 | 2.0 |
| mb18 | 25.3 | 25.4 | 25.5 | 37.6 | 12.1 | 0.2 |
| mb22 | 24.9 | 24.9 | 25.8 | 39.1 | 13.3 | 0.9 |
| mb27 | 24.5 | 24.7 | 24.6 | 37.0 | 12.3 | 0.2 |
| mb29 | 25.5 | 25.7 | 25.2 | 36.8 | 11.1 | -0.3 |
| mb34 | 25.2 | 25.1 | 25.8 | 40.5 | 15.4 | 0.7 |
| mb37 | 24.7 | 24.9 | 26.0 | 38.5 | 13.5 | 1.3 |
| mb40 | 25.0 | 25.3 | 25.3 | 37.1 | 11.9 | 0.3 |
| mb46 | 23.6 | 23.6 | 23.8 | 33.2 | 9.6 | 0.2 |
| mb53 | 23.4 | 23.5 | 23.2 | 31.6 | 8.1 | -0.2 |
| mb60 | 23.1 | 22.7 | 24.5 | 35.5 | 12.8 | 1.4 |
| mb62 | 23.1 | 22.9 | 24.7 | 35.1 | 12.2 | 1.6 |
| mb64 | 23.5 | 23.1 | 23.9 | 33.0 | 9.9 | 0.4 |
| mb68 | 23.8 | 24.8 | 25.7 | 37.6 | 12.7 | 1.9 |
| mb69 | 23.3 | 23.5 | 25.3 | 38.6 | 15.2 | 2.0 |
| mb73 | 23.1 | 24.6 | 24.8 | 36.3 | 11.7 | 1.7 |
| mb76 | 24.5 | 24.7 | 25.6 | 37.0 | 11.4 | 1.1 |
| mb78 | 25.5 | 25.7 | 24.9 | 35.0 | 10.0 | -0.6 |

Table S7-8: Single-plex BDA results for the 21 SNPs used as part of the mBDA qPCR panel for detection of conspecific DNA contaminants in NA18537. Here, we used the PowerUp SYBR Green Master Mix as PCR polymerase, and 400 nM of each primer. During thermocycling, the anneal and extension steps were combined as a 30 s step at 60 °C. 20 ng of genomic DNA was used as input for each reaction.

| SNP locus | NA18562 no Blocker | NA18537 no Blocker | NA18562 w/ Blocker | NA18537 w/ Blocker | ΔCt | Delay |
|---|---|---|---|---|---|---|
| mb1 | 22.0 | 22.2 | 35.9 | 26.4 | 9.5 | 4.2 |
| mb5 | 23.5 | 23.3 | 36.4 | 23.6 | 12.8 | 0.2 |
| mb9 | 21.6 | 21.6 | 34.5 | 22.4 | 12.1 | 0.7 |
| mb11 | 25.1 | 25.0 | 36.2 | 24.7 | 11.4 | -0.3 |
| mb18 | 24.1 | 24.2 | 34.1 | 24.2 | 9.9 | 0.0 |
| mb22 | 25.2 | 25.0 | 34.5 | 25.5 | 9.0 | 0.5 |
| mb27 | 22.1 | 22.3 | 34.5 | 23.5 | 11.0 | 1.2 |
| mb29 | 23.2 | 23.4 | 35.1 | 25.8 | 9.3 | 2.4 |
| mb34 | 22.6 | 22.5 | 35.1 | 23.8 | 11.3 | 1.3 |
| mb37 | 24.3 | 24.3 | 36.3 | 24.9 | 11.4 | 0.6 |
| mb40 | 23.8 | 23.8 | 33.2 | 23.1 | 10.1 | -0.7 |
| mb46 | 24.0 | 24.1 | 37.1 | 25.4 | 11.7 | 1.2 |
| mb53 | 23.4 | 24.0 | 32.2 | 23.1 | 9.1 | -0.9 |
| mb60 | 22.8 | 22.9 | 34.2 | 25.0 | 9.1 | 2.1 |
| mb62 | 23.0 | 22.7 | 34.4 | 23.4 | 11.0 | 0.7 |
| mb64 | 24.3 | 24.4 | 34.3 | 24.2 | 10.1 | -0.2 |
| mb68 | 26.2 | 26.3 | 36.7 | 26.4 | 10.2 | 0.1 |
| mb69 | 23.9 | 24.1 | 35.3 | 23.9 | 11.4 | -0.3 |
| mb73 | 23.2 | 23.3 | 34.4 | 24.1 | 10.4 | 0.7 |
| mb76 | 24.4 | 24.5 | 35.4 | 23.9 | 11.4 | -0.6 |
| mb78 | 25.8 | 25.9 | 33.9 | 25.1 | 8.9 | -0.8 |

Table S7-9: Single-plex BDA results for the 21 SNPs used as part of the mBDA qPCR panel for detection of conspecific DNA contaminants in NA18562. Here, we used the PowerUp SYBR Green Master Mix as PCR polymerase, and 15 nM of each primer. During thermocycling, the anneal and extension steps were combined as a 5 min step at 60 °C. 20 ng of genomic DNA was used as input for each reaction.

**Primer concentration tuning for mBDA qPCR assays.** The 21-plex qPCR assay used in Fig. 5c was subject to one round of primer concentration optimization. Because different primers can hybridize to their respective DNA templates with very different kinetics [1], using identical concentrations for all primers would lead to very different amplification efficiencies, leading to some variant SNPs being detected at much worse sensitivity than others. From an initial concentrations of 15 nM for all primers, we adjusted 17 primer pair concentrations upwards, with the highest primer concentration being in the optimized assay being 100 nM. This optimization was observed to improve our $\Delta$Ct in some cases by over 1 cycle, indicating potential 2-fold improvement in sensitivity through concentration optimization.

For optimized detection of rare contaminants, we need to ensure that the Ct values of all single-plex BDA reactions, when the variant SNP allele is present, are similar. Otherwise, BDA primers with low Ct values will drown out the signals from BDA primers with high Ct values. We can adjust the Ct values of each BDA set by adjusting the primer concentration (lowering the primer concentrations of BDA systems with low Ct values, and increasing the primer concentrations of BDA systems with high Ct values).

A hypothetical 2-plex mBDA assay with unbalanced Ct values is described in the top panel of Table S7-10. The Ct value for the variant allele of SNP1 is 8 cycles smaller than that of SNP2. The pure sample (0% VAF in both SNP1 and SNP2) gives a Ct value of 30, but observing a Ct value of 28 could imply either a 0.4% contaminant of SNP1, or a 100% contaminant of SNP2. Thus, the sensitivity for contaminants with SNP2 is much worse than the sensitivity of contaminants with SNP1. In contrast, a well-balanced 2-plex mBDA assay (bottom panel) provides high sensitivity to all contaminants with variant alleles to either of the SNP loci.

| SNP1 VAF | SNP2 VAF | Ct |
|---|---|---|
| 100% | 100% | 20 |
| 0% | 100% | 28 |
| 100% | 0% | 20 |
| 0% | 0% | 30 |
| 0.40% | 0% | 28 |

| SNP1 VAF | SNP2 VAF | Ct |
|---|---|---|
| 100% | 100% | 20 |
| 0% | 100% | 21 |
| 100% | 0% | 21 |
| 0% | 0% | 30 |
| 0.80% | 0% | 28 |
| 0% | 0.80% | 28 |

Table S7-10: **Top:** Hypothetical qPCR results of poorly balanced 2-plex BDA with different VAF values. An observed Ct value of 28 could correspond to either 0.4% VAF of SNP1, or 100% VAF of SNP2. Thus, this assay is not sensitive to contaminants with SNP1 but not SNP2. **Bottom:** Hypothetical qPCR results of well balanced 2-plex BDA with different VAF values. An observed Ct value of 28 could correspond to either 0.8% VAF of SNP1, or 0.4% VAF of SNP2. Thus, this assay is sensitive to contaminants bearing either SNP1 or SNP2.

**Adjusting the 21-plex mBDA qPCR assay for detecting arbitrary contaminants in NA18537.** The individual Ct values of all 21 single-plex BDA systems that block NA18537's homozygous SNPs, using a constant 15 nM primer concentration, is shown in Fig. S7-2a. This still leads to reasonably good performance in a 21-plex mBDA qPCR assay for detecting conspecific contaminants at the 5% level (Fig. S7-2b) for the 12 contaminants tested. However, the significantly delayed Ct values for some SNPs means that they are essentially not being tested, so the assay would exhibit less sensitivity to other contaminants with variant alleles in the delayed SNP loci.

We adjusted the primer and blocker concentrations based on the initial results in Fig. S7-2a, increasing the primer and blocker concentrations for the SNPs with delayed Ct values. The adjusted primer/blocker concentrations are listed in Table S7-11, and the single-plex BDA qPCR results are shown in Fig. S7-2c and exhibit significantly better uniformity in Ct values both for the variant allele and for the intended allele. The 21-plex mBDA qPCR assay likewise produced larger $\Delta$Ct values, indicating improved sensitivity by the adjusted assay.

**Adjusting the 21-plex mBDA qPCR assay for detecting arbitrary contaminants in NA18562.** Following the same principles as above, we also designed and adjusted a 21-plex BDA panel targeting the NA18562 alleles (Table S7-12, Fig. S7-3). The initial 80-plex SNP loci were all selected such that NA18537 and NA18562

| SNP # | FP Conc. (nM) | RP Conc. (nM) | Blocker Conc. (nM) vs. NA18537 allele |
|---|---|---|---|
| mb1 | 15 | 15 | 150 |
| mb5 | 25 | 25 | 250 |
| mb9 | 15 | 15 | 150 |
| mb11 | 60 | 60 | 600 |
| mb18 | 25 | 25 | 250 |
| mb22 | 100 | 100 | 1000 |
| mb27 | 15 | 15 | 150 |
| mb29 | 25 | 25 | 250 |
| mb34 | 25 | 25 | 250 |
| mb37 | 25 | 25 | 250 |
| mb40 | 25 | 25 | 250 |
| mb46 | 60 | 60 | 600 |
| mb53 | 20 | 20 | 200 |
| mb60 | 25 | 25 | 250 |
| mb62 | 15 | 15 | 150 |
| mb64 | 60 | 60 | 600 |
| mb68 | 25 | 25 | 250 |
| mb69 | 60 | 60 | 600 |
| mb73 | 25 | 25 | 250 |
| mb76 | 60 | 60 | 600 |
| mb78 | 25 | 25 | 250 |

Table S7-11: FP, RP and Blocker concentration of different 21-plex designs for blocker against NA18537.

are homozygous for different alleles, so the 21-plex panel for NA18562 includes a completely different set of oligos. Both this assay and the previous 21-plex mBDA qPCR are selected as a subset from the full list of 80 BDA systems, whose sequences are listed in Supplementary Section S10.

| SNP # | FP Conc. (nM) | RP Conc. (nM) | Blocker Conc. (nM) vs. NA18562 allele |
|---|---|---|---|
| mb1 | 15 | 15 | 150 |
| mb5 | 15 | 15 | 150 |
| mb9 | 15 | 15 | 150 |
| mb11 | 60 | 60 | 600 |
| mb18 | 25 | 25 | 250 |
| mb22 | 60 | 60 | 600 |
| mb27 | 15 | 15 | 150 |
| mb29 | 60 | 60 | 600 |
| mb34 | 25 | 25 | 250 |
| mb37 | 25 | 25 | 250 |
| mb40 | 15 | 15 | 150 |
| mb46 | 15 | 15 | 150 |
| mb53 | 15 | 15 | 150 |
| mb60 | 150 | 150 | 1500 |
| mb62 | 60 | 60 | 600 |
| mb64 | 60 | 60 | 600 |
| mb68 | 15 | 15 | 150 |
| mb69 | 100 | 100 | 1000 |
| mb73 | 60 | 60 | 600 |
| mb76 | 25 | 25 | 250 |
| mb78 | 25 | 25 | 250 |

Table S7-12: FP, RP and Blocker concentration of different 21-plex designs for blocker against NA18562.

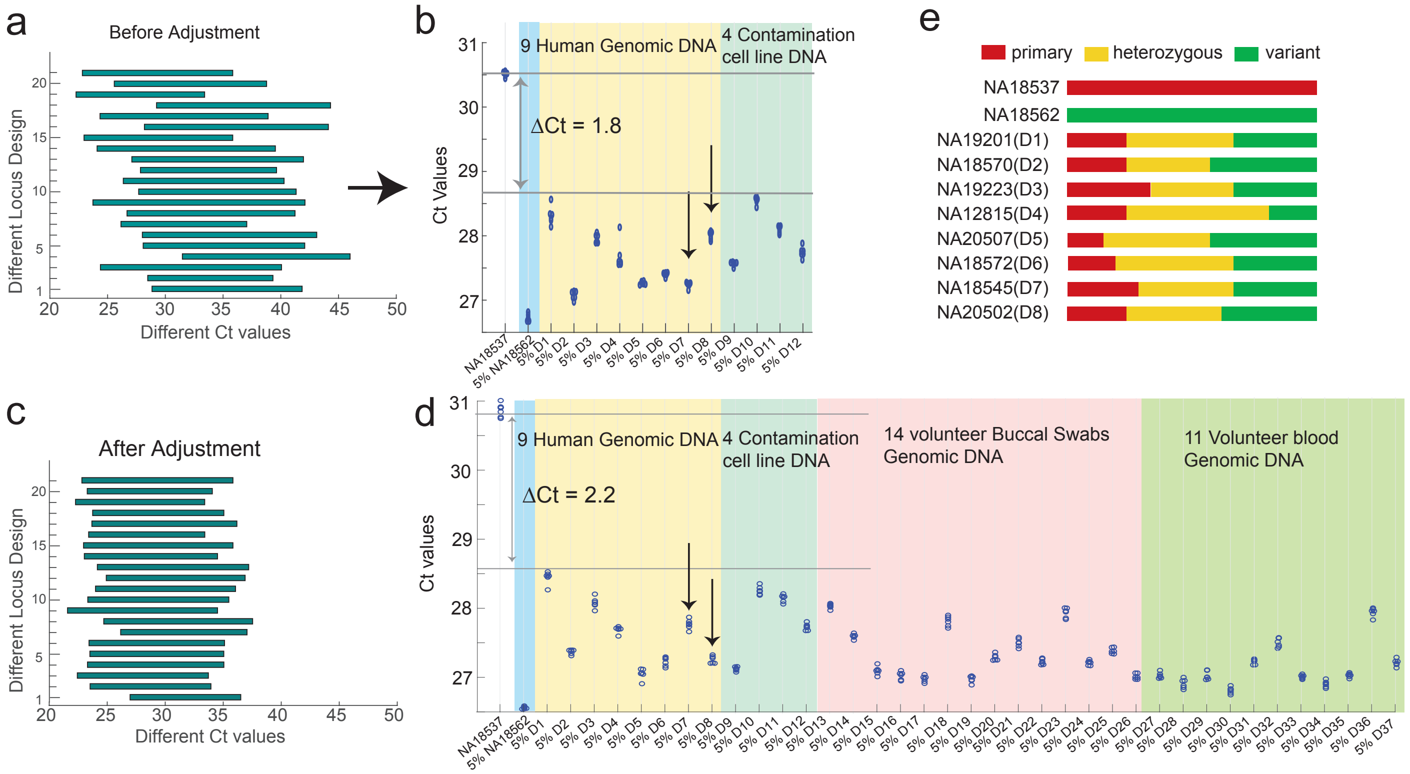


Figure S7-2: Adjusting primer concentration for Ct uniformity improves mBDA qPCR sensitivity to conspecific DNA contaminants. **(a)** Single-plex BDA results for all 21 SNP loci. The left edge of each bar shows the Ct value of 20ng of the NA18562 genomic DNA that is homozygous for the variant allele, and the right edge shows the Ct value of 20ng of the NA18537 genomic DNA that is homozygous for the intended allele. All reactions used 15 nM of each primer, and 150 nM of each Blocker. In this figure, all qPCR reactions were performed using the PowerUp SYBR Green MasterMix (Thermo Fisher), and thermocycling included an anneal/extend step of 5 min at 60 °C. **(b)** Experimental results for the 21-plex mBDA qPCR assay to detect contaminants in NA18537. **(c)** Single-plex BDA results for all 21 SNP loci, using adjusted primer concentrations as listed in Table S7-11. **(d)** Experimental results for the 21-plex mBDA qPCR assay to detect contaminants in NA18537 using the primer-adjusted assay. The minimum ΔCt for 5% contaminant was improved by roughly 0.4 cycles, indicating roughly 2-fold improvement in sensitivity. Sensitivity to some contaminants were improved by more than others (e.g. the right arrows corresponding to D8), due to the contaminant's distribution of variant alleles. D1 to D8 are cell line DNA samples from Coriell Inc.; D9 is DNA from HeLa, D10 is DNA from K562, D11 is DNA from T24, D12 is DNA from PC3. D13 to D26 are DNA extracted from volunteer buccal swabs; D27 to D37 are DNA which extracted from volunteer blood samples. **(e)** Distribution of the number of variant alleles for each contaminant sample, out of the 21 loci.

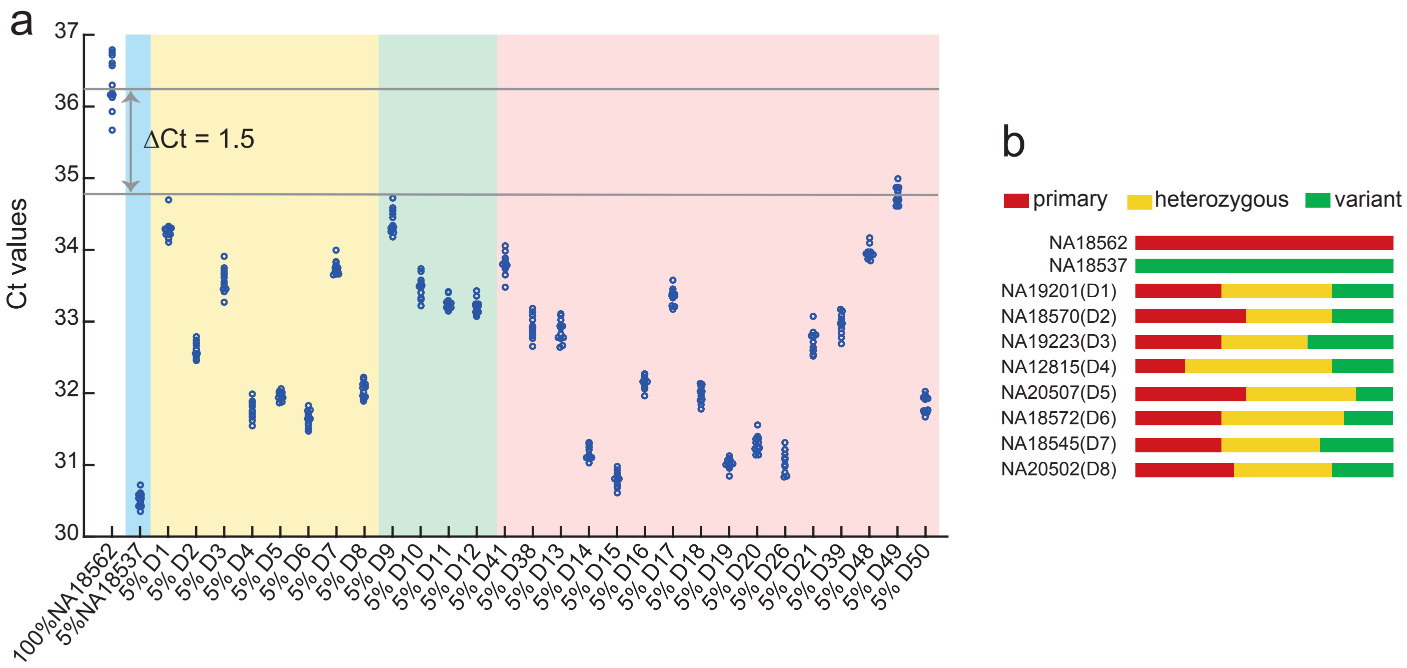


Figure S7-3: 21-plex mBDA qPCR panel for detecting conspecific contaminants in NA18562 **(a)** Experimental results for the 21-plex mBDA qPCR assay to detect contaminants in NA18562 using primer concentrations listed in Table S7-12. In this figure, all qPCR reactions were performed using the PowerUp SYBR Green MasterMix (Thermo Fisher), and thermocycling included an anneal/extend step of 1 min at 60 °C. The input DNA quantity was 20 ng of genomic DNA for all experiments. **(b)** Distribution of the number of variant alleles for each contaminant sample, out of the 21 loci.

Although we expect that the primary interest of cell line researchers is in detection of arbitrary contaminants in known cell lines, it is also possible to use mBDA qPCR assays to detect known contaminants without genotype information regarding the specific intended cell line. In this scenario, we design select SNP loci in which the alternate allele has extremely low frequency in the population, for which the contaminant is heterozygous. As a demonstration, we designed a 3-plex mBDA assay for HeLa-specific SNPs, and experimentally show that we can detect 1% HeLa contaminant in 8 different cell lines without known genotypes (Supplementary Section S4).

[1] Zhang, J. X., *et al.* Predicting DNA hybridization kinetics from sequence. *Nature chemistry,* 10(1), 91 (2018).

#### Section S8: Analysis of Conspecific Contaminant Detection using mBDA qPCR Assay

In this section, we consider two questions: (1) how well would a given mBDA qPCR panel perform across the range of all possible conspecific contaminants, and (2) how easily can we construct a new mBDA qPCR panel for a different intended cell lines? Our conclusions were summarized in Fig. 2bc of the main text; here we describe our assumptions, approach, and potential caveats.

**mBDA qPCR Assay Performance Analysis.** For the 21-plex mBDA qPCR assay to detect arbitrary contaminants in NA18537, NA18537 is either homozygous for the reference allele or for the minor (alternate) allele. Defining the population minor allele frequency as PAF, each diploid human genome is homozygous reference with probability $(1\text{-PAF})^2$, heterozygous with probability $2{\cdot}\text{PAF}{\cdot}(1\text{-PAF})$, and homozygous minor with probability $\text{PAF}^2$. To simulate the distribution of the number of variant alleles (out of 21) for different human DNA contaminant, we performed Monte Carlo simulations of 100,000 human genotypes, with the genotype at each SNP locus following the above probability distribution using PAF values listed in Supplementary Section S1, and assuming that the genotype for each locus is independent. The simulation results (main text Fig. S8-1b) shows that every single simulated genotype possesses at least 8 variant alleles out of the 21 SNP loci.

We can also mathematically calculate the probability that an arbitrary cell line is identical in genotype to NA18537 based on 21 SNP loci as:

$$\Pr(\text{spurious match}) = \prod_{i=1}^{21} (\text{PAF}_i - \text{NA18537}_i)^2$$

where $\text{PAF}_i$ is the population allele frequency of the minor allele as SNP locus $i$, and $\text{NA18537}_i$ is the genotype of NA18537 at the locus $i$ (with $\text{NA18537}_i = 0$ if NA18537 is homozygous for the reference allele, and $\text{NA18537}_i = 1$ for the alternate allele). Using the PAF values listed in Supplementary Section S1, we calculate Pr(spurious match) $= 5.5 \cdot 10^{-16}$, which is far more than roughly 10,000 cell lines in use today.

The central assumption of above analysis is SNP genotype independence across different loci, which may not be true for SNP loci in close proximity on the same chromosome. However, there should be no reason for genotypes of SNP loci on different chromosomes to be not independent. The 21-plex mBDA qPCR panel presented in manuscript Fig. 1 and 2 spans 9 chromosomes, and even considering an unlikely worst-case scenario where all SNP genotypes on a chromosome are perfectly correlated, there is still 9 independent SNPs across different chromosomes, resulting in Pr(spurious match) $= 2.2 \cdot 10^{-7}$. Thus, we are highly confident that with the mBDA qPCR assay will be able to detect arbitrary human DNA contaminants with probability approaching 1.

The other caveat is that many cell lines exhibit significant aneuploidy and/or copy number variations including loss of heterozygosity (e.g. cancer cell lines). Contaminant cell lines with hyperploidy (more than 2 copies of each chromosome) will have lower probability of being homozygous for each intended allele than a standard diploid cells. In contrast, cell lines with hypoploidy or loss of heterozygosity will have higher probability of being hemizygous for the intended allele. A fully hemizygous and genetically uniform genomic DNA sample would have probability of a spurious match being the square root of the standard diploid contaminant:

$$\Pr(\text{spurious match}) = \prod_{i=1}^{21} \text{Abs}(\text{PAF}_i - \text{NA18537}_i)$$

where $\text{Abs}(\cdot)$ is the absolute value function. Thus, the assay is capable of even reliable detection of hemizygous human DNA contaminants.

**mBDA qPCR Assay Construction.** In Supplementary Section S9, we list the BDA sequence designs for 80 different SNP loci, including Blockers for both the reference and alternative alleles for each locus. The purpose of these sequences is to enable the reader to construct their own mBDA qPCR panels to detect arbitrary human DNA contaminants in their specific cell lines of interest. The design workflow can be summarized as the following:

1. Perform NGS or Sanger on the cell line of interest to determine the cell's genotype with respect to the 80 SNP loci.

2. Construct of list of SNP loci in which the cell line is homozygous, and find the appropriate primer/Blocker sequences from Supplementary Section S10.

3. Down-sample to 21-plex, considering the PAF and the chromosomal positions of each of the SNPs, in order to ensure minimal probability of spurious match by an unknown human DNA contaminant.

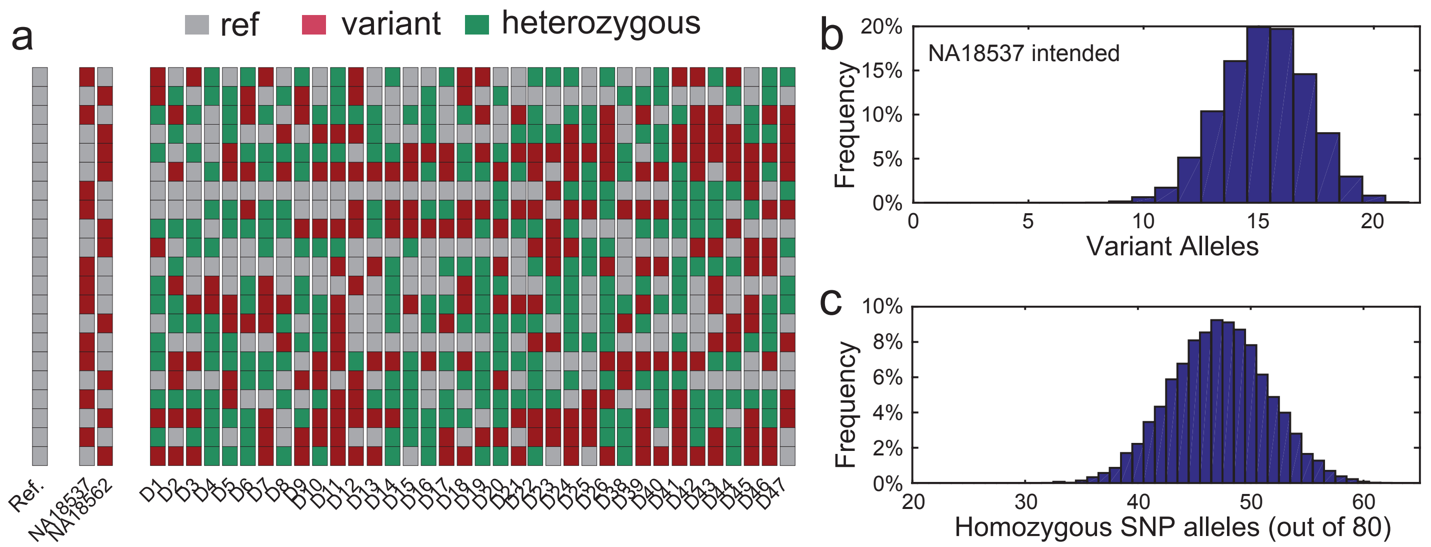


FIG. S8-1. Detecting cell line contamination via low VAF variant SNP alleles. **(a)** SNP genotypes for the 1000 Genomes reference DNA, the NA18537 (wildtype) sample, the NA18562 (homozygous variant) sample, and all other contaminant DNA samples used for this work. See Supplementary Excel spreadsheet for sample details. **(b)** Simulated distribution of the number of variant alleles out of the 21 SNPs observed, based on reported SNP population allele frequencies and assuming independence between SNPs. **(c)** Simulated distribution of the number of homozygous alleles out of the 80 SNP loci that we used to design our 80-plex mBDA set, based on reported SNP population allele frequencies and assuming independence between SNPs.

To simulate the distribution of the number of SNP loci (out of 80) that would be homozygous for an arbitrary intended cell line, we performed Monte Carlo simulations of 100,000 human genotypes, with the genotype at each SNP locus following the above probability distribution using published PAF values, and assuming that the genotype for each locus is independent. The simulation results shows that every single simulated genotype possesses at least 31 homozygous loci out of 80.

As in the analysis of mBDA qPCR assay performance, the main caveats of this analysis are that we assume genotype independence across different SNP loci, and that the intended cell line is diploid. If there is strong genetic linkage between different SNPs on the same chromosome, then it is possible that fewer loci are homozygous than expected (e.g. one copy of the chromosome is genotype A-A-A-A-A, and the other copy is genotype a-a-a-a-a). If the cell line has copy number variations or hyperploidy, then the probability that the SNP loci that are affected is homozygous would be reduced, also resulting in lower than the expected number of homozygous SNPs out of 80.

For a given intended cell line, more than 21 SNP loci of the 80-plex mBDA panel may be homozygous.

Even in these cases, we recommend to down-select the list of SNPs to 21-plex for constructing mBDA qPCR assay for detecting arbitrary contaminants for several reasons. First, at higher multiplex, there is higher chance of primer dimers forming stochastically, which could result in false-positive signals due to the intercalating dye readout. For the qPCR assays we tested here, we observed little primer dimer formation even up to 40-plex. However, given that there are $\binom{2N}{2}$ pairwise combinations of primers that could potentially produce primer dimers, reducing from 40-plex to 21-plex reduces the amount of potential primer dimers by more than 4-fold.

Second, the primer concentration adjustment to achieve relatively uniform Ct values for each single-plex BDA system. In higher-plex mBDA qPCR assays, more work is required for primer concentration adjustment. For these reasons, we do not recommend constructing mBDA qPCR assays with higher plex than needed.

#### Section S9: Detection of Known Contaminants in Intended Cell Lines of Unknown Genotype

For many specialized human cell lines, SNP genotyping by NGS to determine the composition of a qPCR mBDA kit for detecting arbitrary contaminants may require more effort than researchers are willing to spend. For these rarely used cell lines, we show that mBDA qPCR assays can be used to detect common known contaminants. Here, we show proof of concept on detecting HeLa contaminations, because HeLa is the single most frequent source of human cell and DNA contamination.

To detect HeLa contamination in an arbitrary human cell line, we selected 3 HeLa-specific sequence variants from the Catalogue Of Somatic Mutations In Cancer (COSMIC) database (Table S10-1). These variants include 2 SNPs and 1 deletion (15 nt), and were not reported to be present in any other of the other 1014 cell lines in the COSMIC database. Although it is possible that an arbitrary intended cell line may not be extensively genotyped and thus could potentially include 1 of the 3 variants in common with HeLa, probabilistically it is highly unlikely that the intended cell line would possess all 3 variants.

To independently observe the simultaneous presence of all 3 variants in the same sample, we use 3 Taqman probes with spectrally distinct fluorophores (Cy5, TEX612, and HEX). The Taqman probes each bind their respective amplicon at a sequence downstream of the mutation of interest, and thus are not allele specific (Fig. S10-1). Detection of the HeLa-specific mutations is thus based on the suppression of PCR amplification of wildtype alleles at the mutant loci.

Fig. S10-2 and S10-3 show the experimental qPCR results of the 3-plex mBDA assay detection HeLa contaminants in 8 different intended cell lines. For each intended cell line, we tested a pure genomic DNA sample (0% HeLa) as well as samples with 1% and 3% HeLa genomic DNA spike-in. For each sample, qPCR reactions were run in triplicate.

Given the intrinsic variability in Ct values for qPCR reactions, especially for 0% samples, we note that we can mathematically increase the confidence of contamination calls be integrating the Ct values of all 3 HeLa-specific mutants into a single metric, an adjusted median Ct. In the HEX channel, the mean Ct value for 1% HeLa across all 8 intended cell lines was 32.68; for Cy5, 29.84; for TEX615, 30.86. Thus, the HEX channel Ct values are generally higher than TEX615 by 1.83 cycles, and the Cy5 channel Ct values are generally lower than TEX615 by 1.02 cycles. The adjusted median Ct for a sample is calculated as Median([$Ct_{HEX}$-1.83, $Ct_{TEX615}$, $Ct_{Cy5}$ + 1.02]).

From this adjusted median Ct value metric, we see that there is over a 3 cycle Ct difference between 0% HeLa and 1% HeLa for all intended cell lines. Setting an adjusted median Ct cutoff of 33, all uncontaminated intended cell lines have a higher value, and all 1% HeLa contaminated cell lines have a lower value.

| No. | CDS Mutation | Strand | primary Genotype | HeLa Genotype | Chr | Position(GRCh37) | Taqman Probe Dye |
|---|---|---|---|---|---|---|---|
| 1 | INTS3 c.2268_2282del15 | + | CGGAGAGCTGCTGAA/CG...AA | CG...AA/- | 1 | 153741392-153741406 | HEX |
| 2 | APEH c.937C>T | + | C/C | C/T | 3 | 9714395 | Cy5 |
| 3 | ATP2C2 c.898C>G | + | C/C | C/G | 16 | 84456848 | TEX615 |

TABLE S9-1: HeLa-specific mutations targeted for mBDA detection.

| Cell Line | Origin Tissue |
|---|---|
| K562 | Human blood (chronic myelogenous leukemia) |
| T24 | Human bladder carcinoma |
| HL60 | Human leukemia |
| NA18537 | Human blood B-Lymphocyte |
| PC3 | Human prostate cancer |
| NA18562 | Human blood B-Lymphocyte |
| HEK293 | Human embryonic kidney |
| NA18537 | human lung adenocarcinoma |

TABLE S9-2: Intended cell lines tested for HeLa contamination detection.

| Design# | FP | RP | blocker | probe |
|---|---|---|---|---|
| INTS3 | AGTTTCCAGATGAAACCTTGAGGA | GGCCCTATGTCCCTAATGCT | CCTTGAGGAGCGGAGAGCTGCTG/3SpC3/ | /5HEX/TATTGACTC /ZEN/TGCACAGGTGAACATTGAGCTCT/3IABkFQ/ |
| APEH | GCTGAGCCCAGACCAATGT | TCTCAACCTCTAGGCCCTCC | AGACCAATGTCGCATTGTCTACCTGC/3SpC3/ | /5Cy5/CAAGAAGAT /TAO/GGGATGGTGGATGAGAGCTGAAG/3IAbRQSp/ |
| ATP2C2 | GGACAGGCTAGGAAAGCAACT | ATGATGGCTCCCCTCCATCT | GAAAGCAACTGACACTCTTCTCCTTTGG/3SpC3/ | /5TEX615/AATCGGTGAGTGAAGCAG TTTCCATACTGGG/3IAbRQSp/ |

TABLE S9-3: Primer and Blocker oligonucleotide sequences for the 3-plex mBDA qPCR to detect HeLa-specific SNPs. We used Taqman probe readouts for this application; Taqman probe sequences are also listed here.

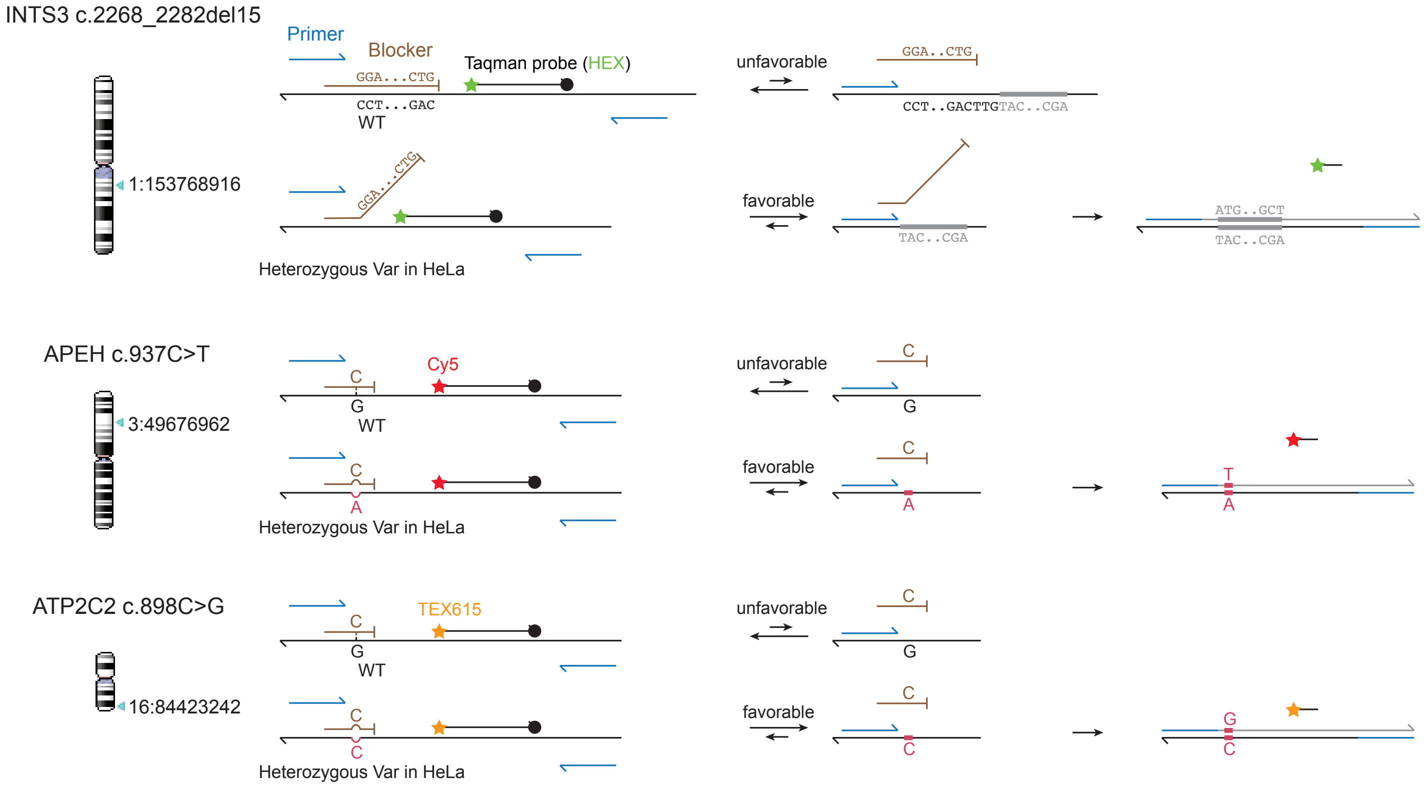


FIG. S9-1 : Illustration of mBDA primers, blockers, and Taqman probes used for detection of 3 HeLa-specific mutations for detection of HeLa contamination in arbitrary intended cell lines. The Taqman probes bind a region of the amplicon downstream of the mutation, and are not allele-specific.

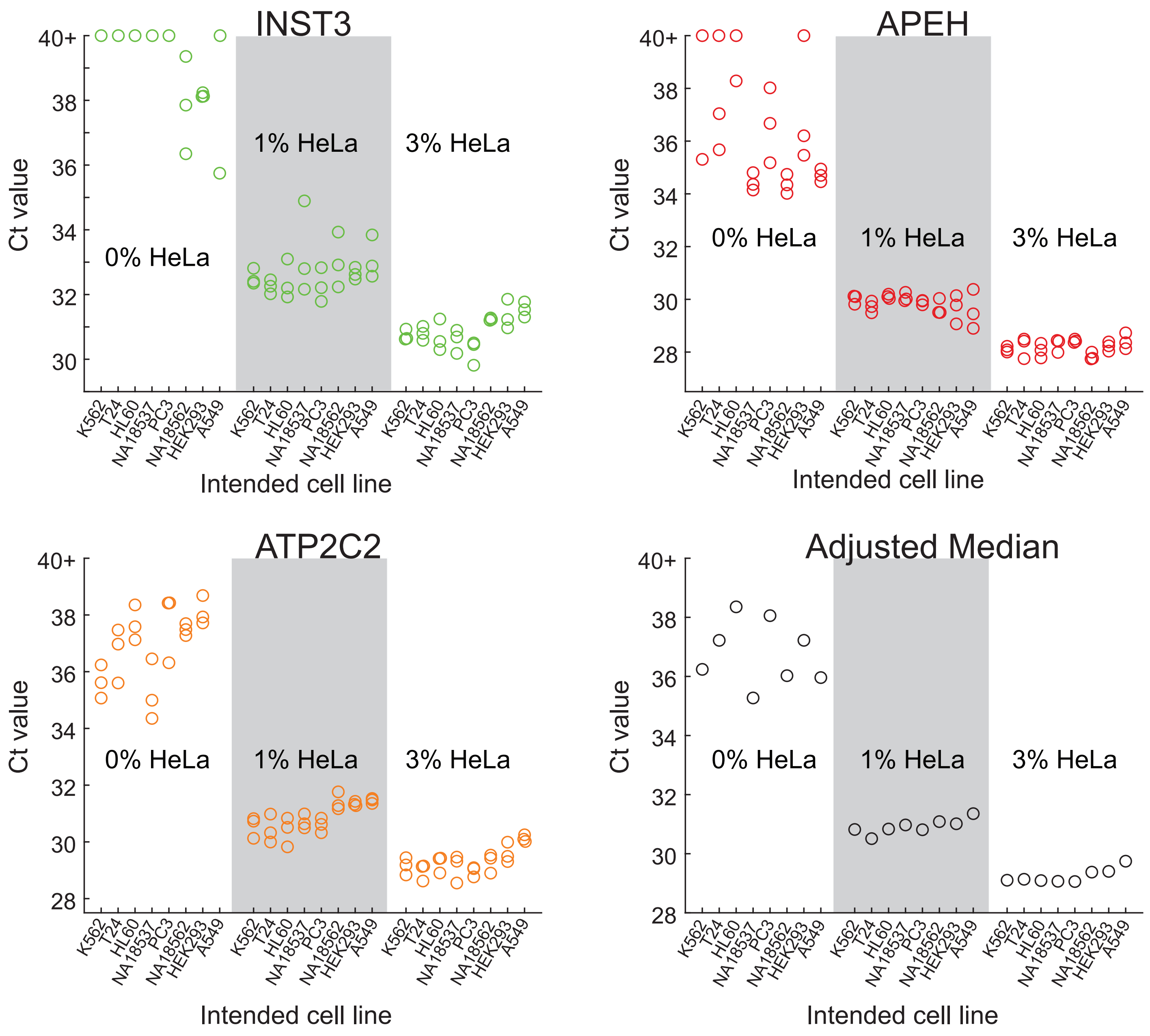


FIG. S9-2 : Summary of qPCR results for mBDA detection of HeLa contaminant in 8 different intended cell lines. Each qPCR reaction was performed in triplicate, using 20 ng of genomic DNA as input (e.g. 1% HeLa indicates 0.2 ng HeLa and 19.8 ng of the intended cell line). The adjusted median Ct value is the median of 9 total Ct values for each sample, with HEX channel Ct values adjusted by -1.83 and Cy5 channel Ct values adjusted by +1.02. The adjusted median Ct value provides the most robust measure of HeLa contamination fraction.

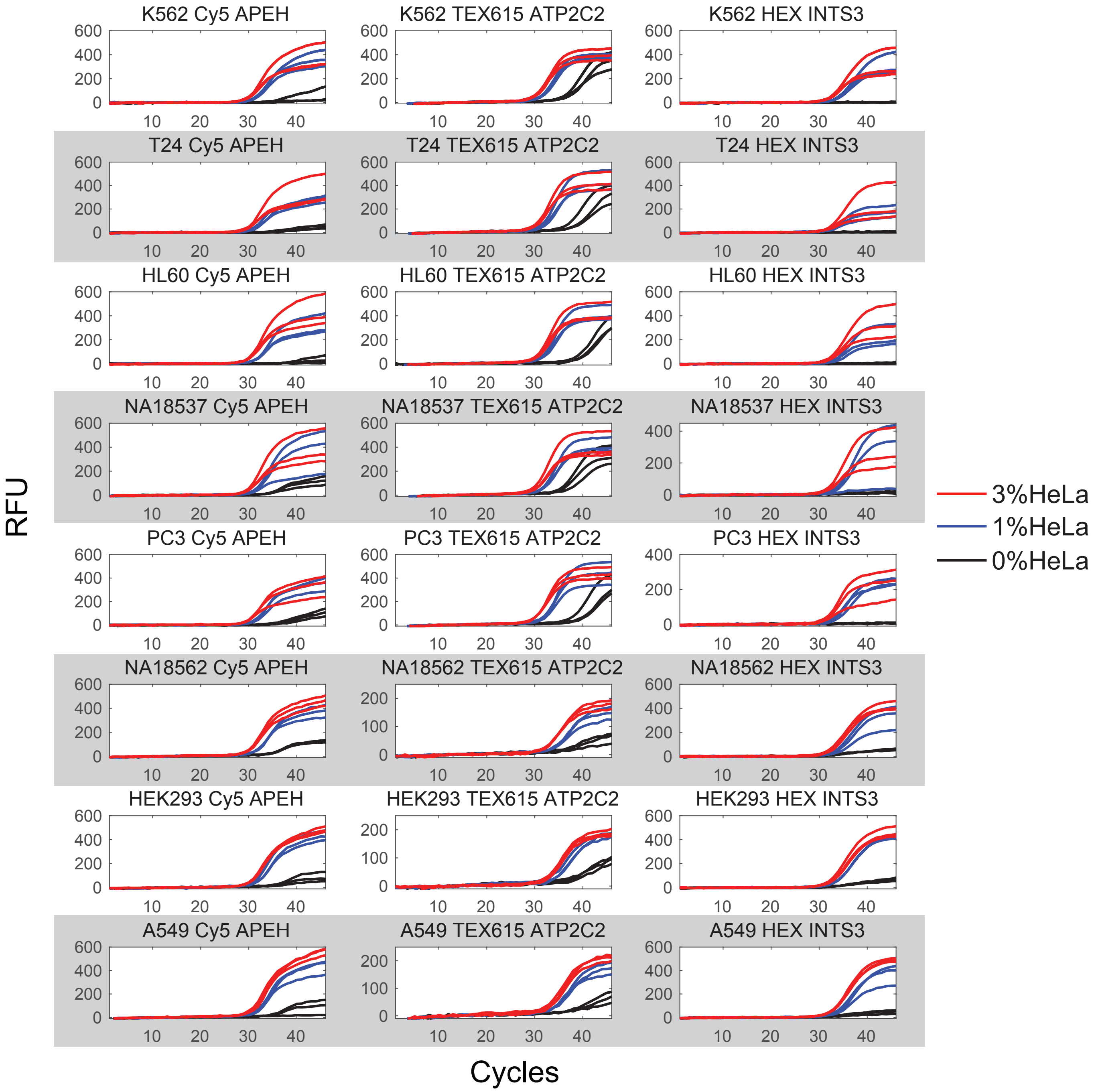


FIG. S9-3 : qPCR traces for the 3-plex mBDA for detecting HeLa contamination.

#### Section S10: Cancer Panel Contents and EF Calibration

We designed the mBDA NGS melanoma panel to include frequently observed mutations and insertions/deletions (indels) in the COSMIC database (Table S10-1). The design of the mBDA primers and blockers follow the principles described in Section S1. The final panel includes 16-plex BDA designs that collectivelyl cover 145 mutations and indels across 9 genes. We calibrated the VAF limit of detection and EF values for 28 of the covered mutations, using reference samples constructed by mixing the NA18562 cell line gDNA with different concentrations of synthetic gBlock DNA bearing the mutations of interest (Integrated DNA Technologies).

In data analysis part, we build 145 mutations/indels variant sequences and each BDA designs also had the WT sequences. The total number of WT and variant sequences are 161. After build the sequences, the perfect matched reads were counted by Matlab codes and saved as WT reads and variants reads in each locus. The VRF, VAF and EF calculations were followed the methods in section S2 and S4.

**Cancer Panel Designs Validation and EF Calculation.** Positive control reference samples were constructed that included a total of 28 of the COSMIC mutations contained with the mBDA panel. The reasons we calibrated these 28 and not others was because these were the mutations that had significant VRF in at least one of the clinical tissue samples. Thus, we actually performed calibration experiments after testing clinical samples, in order to minimize the time and cost of mutation EF calibration.

Reference samples were constructed by mixing synthetic gBlock DNA, each approximately 500 nt with the mutations roughly in the middle of the gBlock with NA18562 human genomic DNA. The gBlocks were diluted to nominally $10^5$ molecules / $\mu$L and then re-quantitated using qPCR vs. the NA18562 gDNA for each locus. Using the corrected concentrations, the gBlocks were then pooled to form reference sample mixtures with mutation VAFs of 10%, 5%, 2%, 1%, 0.5%, and 0.2%. Importantly, the lower VAF references samples were constructed by serial dilution from higher VAF reference samples in NA18562.

Using the observed VRFs from the 5 reference samples with VAFs of 5%, 2%, 1%, 0.5%, and 0.2%, we fitted the EF values as described in Supplementary Section S4. The 0% VAF (pure NA18562) was also run to characterize the VAF limit of detection of the panel. The 10% VAF reference sample was run without blocker to confirm the VAFs of the reference samples. The number of NGS reads used for each library and the on-target rates are shown in Fig. S10-3. The VRF vs. VAF plots are shown in Fig. S10-1, and the relative VAF quantitation accuracy is shown in Fig. S10-2. The EF values for the 28 mutations are summarized in Table S10-4 and plotted in Fig. S10-3.

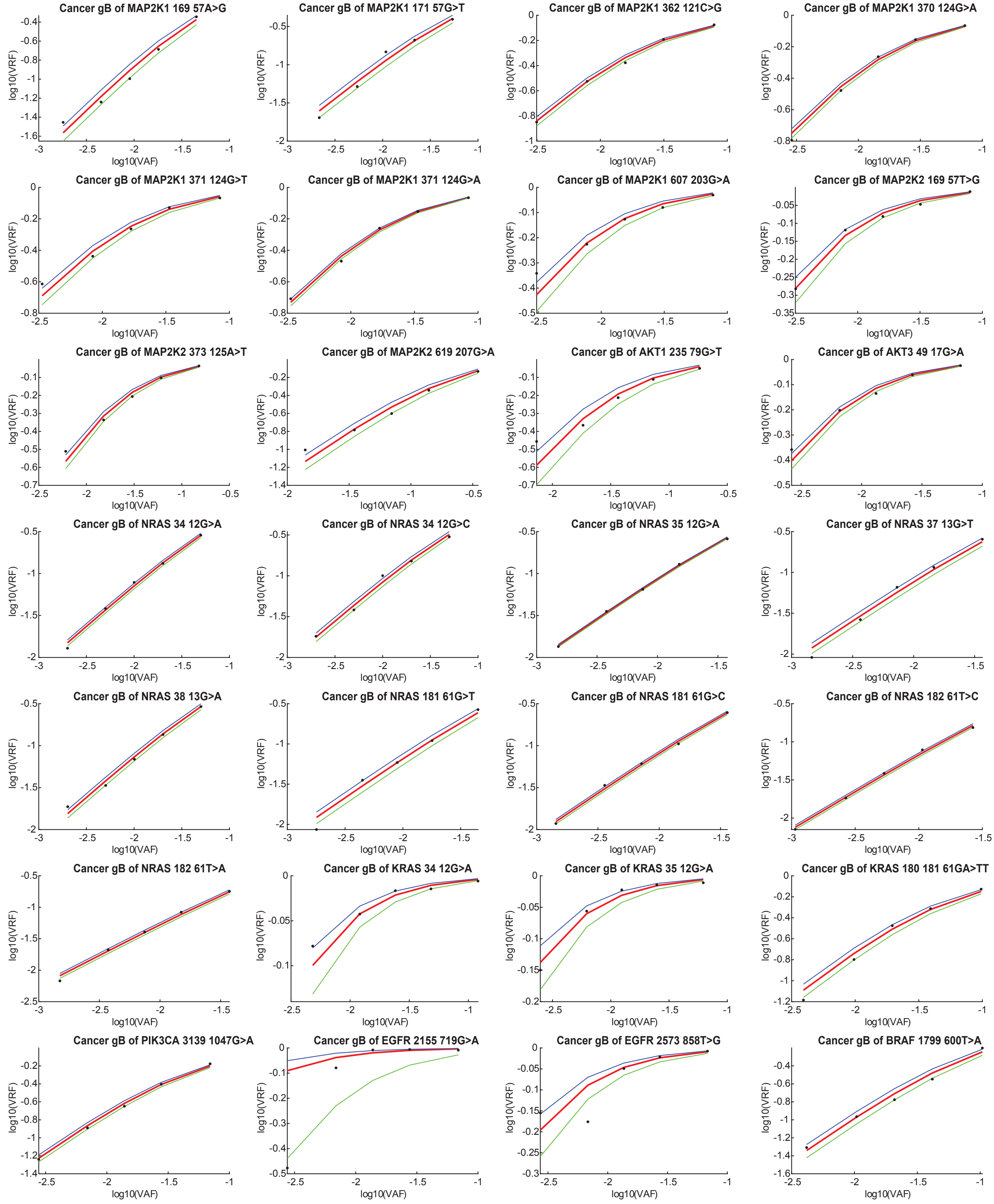


FIG. S10-1. VRF observed for different VAF reference samples for the 28 mutations calibrated. Red lines indicate the mean of five different VAF values, and the blue and green lines show ±1 standard deviation. Black dots indicate observed EF values.

| Number | BDA | Loci Information | COSMIC ID |
|---|---|---|---|
| 1 | MAP2K1_1 | MAP2K1_169_57A>G | COSM5369532 |
| 2 | | MAP2K1_170_57A>C | COSM4756761 |
| 3 | | MAP2K1_171_57G>C | COSM5520914 |
| 4 | | MAP2K1_171_57G>T | COSM1235478 |
| 5 | MAP2K1_2 | MAP2K1_361_121A>T | COSM555601 |
| 6 | | MAP2K1_362_121C>G | COSM1315829 |
| 7 | | MAP2K1_370_124G>A | COSM235614 |
| 8 | | MAP2K1_370 371_124GG>TA | COSM6972727 |
| 9 | | MAP2K1_371_124G>T | COSM1167912 |
| 10 | | MAP2K1_371_124G>C | COSM6986263 |
| 11 | | MAP2K1_371_124G>A | COSM1315861 |
| 12 | MAP2K1_3 | MAP2K1_607_203G>A | COSM232755 |
| 13 | | MAP2K1_608_203A>T | COSM3386991 |
| 14 | MAP2K2_1 | MAP2K2_169_57T>G | COSM3534171 |
| 15 | | MAP2K2_169_57T>C | COSM1235618 |
| 16 | | MAP2K2_170_57T>G | COSM5546577 |
| 17 | | MAP2K2_171_57T>A | COSM3389034 |
| 18 | | MAP2K2_179_60A>C | COSM1236207 |
| 19 | MAP2K2_2 | MAP2K2_373_125A>T | COSM5855815 |
| 20 | | MAP2K2_374_125C>G | COSM5487841 |
| 21 | | MAP2K2_376_126T>C | COSM1731742 |
| 22 | MAP2K2_3 | MAP2K2_619_207G>A | COSM5574290 |
| 23 | AKT1 | AKT1_235_79G>T | COSM159008 |
| 24 | | AKT1_236_79T>C | COSM6598005 |
| 25 | AKT3 | AKT3_49_17G>A | COSM224779 |
| 26 | | AKT3_49_17G>T | COSM6937602 |
| 27 | NRAS_1 | NRAS_34_12G>A | COSM563 |
| 28 | | NRAS_34_12G>C | COSM561 |
| 29 | | NRAS_34_12G>T | COSM562 |
| 30 | | NRAS_35_12G>A | COSM564 |
| 31 | | NRAS_35_12G>C | COSM565 |
| 32 | | NRAS_35_12G>T | COSM566 |
| 33 | | NRAS_34 35_12GG>AA | COSM12723 |
| 34 | | NRAS_34 35_12GG>CC | COSM559 |
| 35 | | NRAS_34 35_12GG>TA | COSM560 |
| 36 | | NRAS_35 36_12GT>AG | COSM144577 |
| 37 | | NRAS_37_13G>A | COSM571 |
| 38 | | NRAS_37_13G>C | COSM569 |
| 39 | | NRAS_37_13G>T | COSM570 |
| 40 | | NRAS_38_13G>A | COSM573 |
| 41 | | NRAS_38_13G>C | COSM575 |
| 42 | | NRAS_38_13G>T | COSM574 |
| 43 | | NRAS_37 38_13GG>AA | COSM24668 |
| 44 | | NRAS_37 38_13GG>TA | COSM568 |
| 45 | | NRAS_37 38_13GG>TT | COSM6970923 |
| 46 | NRAS_2 | NRAS_180 181_61GT>TA | COSM12730 |
| 47 | | NRAS_181_61G>T | COSM580 |
| 48 | | NRAS_181_61G>C | COSM581 |
| 49 | | NRAS_181_61G>A | COSM6005482 |
| 50 | | NRAS_182_61T>G | COSM582 |
| 51 | | NRAS_182_61T>C | COSM584 |
| 52 | | NRAS_182_61T>A | COSM583 |
| 53 | | NRAS_181 182_61TG>AA | COSM12725 |
| 54 | | NRAS_181 182_61TG>GT | COSM5044300 |
| 55 | | NRAS_181 182_61TG>CT | COSM579 |
| 56 | | NRAS_183_61T>G | COSM586 |
| 57 | | NRAS_183_61T>A | COSM585 |
| 58 | | NRAS_182 183_61TT>CC | COSM33693 |
| 59 | | NRAS_182 183_61TT>CA | COSM30646 |
| 60 | | NRAS_181 182 183_61TTG>CTT | COSM53223 |
| 61 | | NRAS_183 184_61CT>TG | COSM26494 |
| 62 | | NRAS_183delA_61delT | COSM5044298 |
| 63 | KRAS_1 | KRAS_33 34_12TG>CT | COSM6950487 |
| 64 | | KRAS_34_12G>A | COSM517 |
| 65 | | KRAS_34_12G>C | COSM518 |
| 66 | | KRAS_34_12G>T | COSM516 |
| 67 | | KRAS_35_12G>A | COSM521 |
| 68 | | KRAS_35_12G>C | COSM522 |
| 69 | | KRAS_35_12G>T | COSM520 |
| 70 | | KRAS_34 35_12GG>AA | COSM13643 |
| 71 | | KRAS_34 35_12GG>AT | COSM34144 |
| 72 | | KRAS_34 35_12GG>CT | COSM514 |
| 73 | | KRAS_34 35_12GG>TA | COSM25081 |

| Number | Design | Loci Information | COSMIC ID |
|---|---|---|---|
| 74 | KRAS_1 | KRAS_34 35_12GG>TT | COSM512 |
| 75 | | KRAS_34 35_12GG>TC | COSM6959653 |
| 76 | | KRAS_35 36_12GT>AA | COSM519 |
| 77 | | KRAS_35 36_12GT>AC | COSM14209 |
| 78 | | KRAS_35 36_12GT>AG | COSM30566 |
| 79 | | KRAS_35 36_12GT>CA | COSM5413585 |
| 80 | | KRAS_35 36_12GT>TC | COSM515 |
| 81 | | KRAS_35 36_12GT>TG | COSM4387522 |
| 82 | | KRAS_34 35 36_12GGT>CTG | COSM1716372 |
| 83 | | KRAS_34 35 36_12GGT>TGC | COSM513 |
| 84 | | KRAS_34 35 36_12GGT>TGG | COSM36281 |
| 85 | | KRAS_36_37insGGT_1236_37insGGT | COSM12655 |
| 86 | | KRAS_34 35 36_12GGT>AGA | COSM249888 |
| 87 | | KRAS_36 37_13TG>AC | COSM4745557 |
| 88 | | KRAS_36 37_13TG>AT | COSM87281 |
| 89 | | KRAS_37_13G>A | COSM528 |
| 90 | | KRAS_37_13G>C | COSM529 |
| 91 | | KRAS_37_13G>T | COSM527 |
| 92 | | KRAS_38_13G>A | COSM532 |
| 93 | | KRAS_38_13G>C | COSM533 |
| 94 | | KRAS_38_13G>T | COSM534 |
| 95 | | KRAS_37 38_13GG>AA | COSM53283 |
| 96 | | KRAS_37 38_13GG>AT | COSM525 |
| 97 | | KRAS_37 38_13GG>CC | COSM5985091 |
| 98 | | KRAS_37 38_13GG>TA | COSM1666974 |
| 99 | | KRAS_37 38_13GG>TT | COSM1685355 |
| 100 | | KRAS_38 39_13GC>AA | COSM87280 |
| 101 | | KRAS_38 39_13GC>AG | COSM30567 |
| 102 | | KRAS_38 39_13GC>AT | COSM531 |
| 103 | | KRAS_38 39_13GC>TG | COSM530 |
| 104 | | KRAS_38 39_13GC>TT | COSM12721 |
| 105 | | KRAS_37 38 39_13GGC>CGT | COSM526 |
| 106 | | KRAS_37 38 39_13GGC>AAG | COSM5415923 |
| 107 | KRAS_2 | KRAS_180 181_61GA>TT | COSM87298 |
| 108 | | KRAS_180 181_61GA>TC | COSM6904743 |
| 109 | | KRAS_181_61G>T | COSM549 |
| 110 | | KRAS_181_61G>C | COSM550 |
| 111 | | KRAS_182_61T>G | COSM551 |
| 112 | | KRAS_182_61T>C | COSM552 |
| 113 | | KRAS_182_61T>A | COSM553 |
| 114 | | KRAS_181 182_61TG>GC | COSM6950503 |
| 115 | | KRAS_182 183_61TT>CG | COSM6932880 |
| 116 | | KRAS_182 183_61TT>GC | COSM6929725 |
| 117 | | KRAS_182 183_61TT>AC | COSM1168052 |
| 118 | | KRAS_182 183_61TT>GA | COSM1166780 |
| 119 | | KRAS_182 183_61TT>CA | COSM6978958 |
| 120 | | KRAS_182 183_61TT>AA | COSM6975270 |
| 121 | | KRAS_183_61T>G | COSM554 |
| 122 | | KRAS_183_61T>A | COSM555 |
| 123 | PIK3CA | PIK3CA_3139_1047G>T | COSM5029129 |
| 124 | | PIK3CA_3139_1047G>A | COSM774 |
| 125 | | PIK3CA_3140_1047T>G | COSM249874 |
| 126 | | PIK3CA_3140_1047T>C | COSM775 |
| 127 | | PIK3CA_3140_1047T>A | COSM776 |
| 128 | | PIK3CA_3141_1047A>T | COSM1041525 |
| 129 | | PIK3CA_3141_1047A>C | COSM24714 |
| 130 | | PIK3CA_3140-3143_1047TGAT>CGAC | COSM6949966 |
| 131 | EGFR_1 | EGFR_2155_719G>A | COSM6252 |
| 132 | | EGFR_2155_719G>T | COSM6253 |
| 133 | | EGFR_2156_719G>C | COSM6239 |
| 134 | EGFR_2 | EGFR_2573_858T>G | COSM6224 |
| 135 | BRAF | BRAF_1789_1790_597delinsTC | COSM1126 |
| 136 | | BRAF_1789_597C>G | COSM470 |
| 137 | | BRAF_1790_597T>A | COSM1125 |
| 138 | | BRAF_1790_597T>G | COSM471 |
| 139 | | BRAF_1790_597T>C | COSM1448593 |
| 140 | | BRAF_1798_1799_600delinsCG | COSM1583011 |
| 141 | | BRAF_1798_1799_600delinsAG | COSM474 |
| 142 | | BRAF_1799_1800_600delinsAA | COSM475 |
| 143 | | BRAF_1799_600T>A | COSM476 |
| 144 | | BRAF_1799_1801_601del | COSM1133 |
| 145 | | BRAF_1801_601A>G | COSM478 |

TABLE S10-1. List of 145 COSMIC mutations included in the melanoma mBDA NGS panel.

| Library | Template | Blocker w/o | Total Reads | On-Target Reads | On-Target Rate |
|---|---|---|---|---|---|
| lib1 | 0.2% | wB | 67,573 | 57,828 | 85.6% |
| lib2 | 0.50% | wB | 91,774 | 78,083 | 85.1% |
| lib3 | 1% | wB | 68,737 | 57,119 | 83.1% |
| lib4 | 2% | wB | 64,953 | 55,079 | 84.8% |
| lib5 | 5% | wB | 70,462 | 60,104 | 85.3% |
| lib6 | 10% | noB | 83,142 | 72,701 | 87.4% |
| lib7 | NA18562 | wB | 57,022 | 48,035 | 84.2% |

TABLE S10-2. Total NGS reads, on-target NGS reads, and on-target rates for the libraries constructed from references samples used for EF calibration. All libraries used 50 ng of input DNA, corresponding to roughly 15,000 haploid NA18562 genomes with a small number of gBlock molecules.

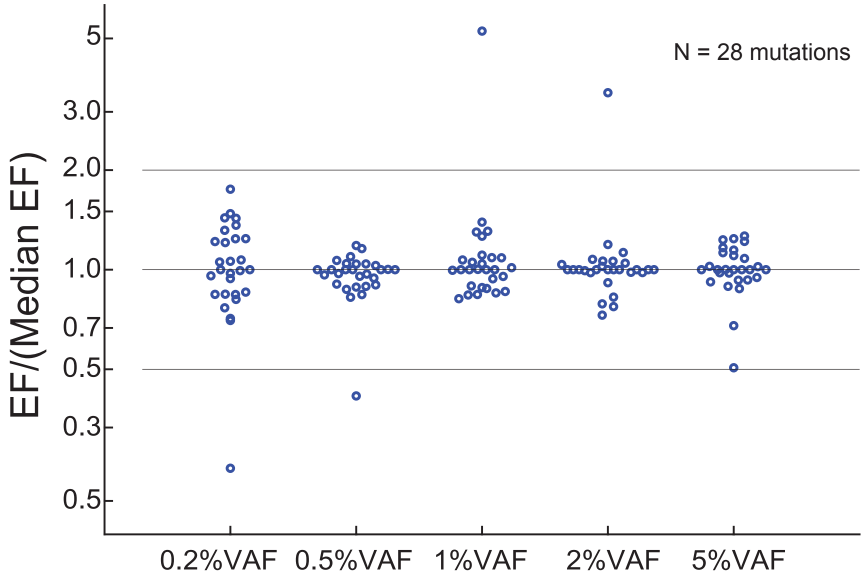


FIG. S10-2. Relative VAF quantitation accuracy, visualized as the observed EF at different VAF values divided by the median EF.

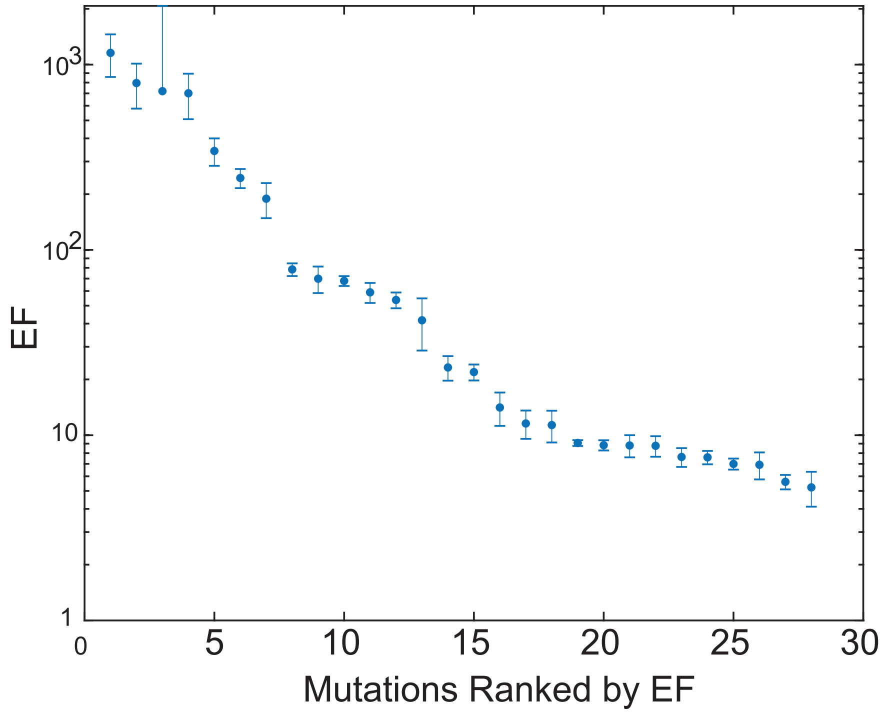


FIG. S10-3. Summary of median EF values for the calibrated 28 mutations. Error bars show minimum and maximum calculated EF values from the 5 different reference samples.

| BDA Name | gBlock No. | gBlock Name | Median EF |
|---|---|---|---|
| MAP2K1_1 | 1 | MAP2K1_169_57A>G | 14 |
| | 2 | MAP2K1_171_57G>T | 11 |
| MAP2K1_2 | 3 | MAP2K1_362_121C>G | 54 |
| | 4 | MAP2K1_370_124G>A | 78 |
| | 5 | MAP2K1_371_124G>T | 70 |
| | 6 | MAP2K1_371_124G>A | 68 |
| MAP2K1_3 | 7 | MAP2K1_607_203G>A | 189 |
| MAP2K2_1 | 8 | MAP2K2_169_57T>G | 342 |
| MAP2K2_2 | 9 | MAP2K2_373_125A>T | 59 |
| MAP2K2_3 | 10 | MAP2K2_619_207G>A | 5 |
| AKT1 | 11 | AKT1_235_79G>T | 42 |
| AKT3 | 12 | AKT3_49_17G>A | 244 |
| NRAS_1 | 13 | NRAS_34_12G>A | 8 |
| | 14 | NRAS_34_12G>C | 9 |
| | 15 | NRAS_35_12G>A | 9 |
| | 16 | NRAS_37_13G>T | 9 |
| | 17 | NRAS_38_13G>A | 8 |
| NRAS_2 | 18 | NRAS_181_61G>T | 7 |
| | 19 | NRAS_181_61G>C | 9 |
| | 20 | NRAS_182_61T>C | 7 |
| | 21 | NRAS_182_61T>A | 6 |
| KRAS_1 | 22 | KRAS_34_12G>A | 796 |
| | 23 | KRAS_35_12G>A | 1157 |
| KRAS_2 | 24 | KRAS_180 181_61GA>TT | 23 |
| PIK3CA | 25 | PIK3CA_3139_1047G>A | 22 |
| EGFR_1 | 26 | EGFR_2155_719G>A | 719 |
| EGFR_2 | 27 | EGFR_2573_858T>G | 701 |
| BRAF | 28 | BRAF_1799_600T>A | 12 |

TABLE S10-3. Median EF values for the 28 mutations calibrated for the melanoma mBDA NGS panel.

#### Section S11: Cancer Panel Analytic Validation vs ddPCR

Digital PCR such as Bio-Rad ddPCR is a commonly used method to validate the low VAF values in NGS results. Here, we report VAF calls using mBDA NGS vs. from ddPCR, on 4 mutations: MAP2K2_373_125_A>T, NRAS_34_12_G>A, and NRAS_181_61_G>T. For the NRAS mutations, Bio-Rad offers pre-validated primers/probes, which we purchased. For the MAP2K2 mutation, we designed primers and probe based on the Bio-Rad user's manual to the best of our abilities. The results are displayed in Fig. S11-1.

| | mBDA NGS panel VAF | | | ddPCR VAF | | |
|---|---|---|---|---|---|---|
| | NA18562 | 0.2% gBlock | 5% gBlock | NA18562 | 0.2% gBlock | 5% gBlock |
| MAP2K2 (p.125 c.373A>T) | 0.06% | 0.60% | 14.60% | 0.00% | 0.39% | 9.50% |
| NRAS-G12S (p.12 c.34G>A) | 0.04% | 0.20% | 5.00% | 0.00% | 0.55% | 11.92% |
| NRAS-Q61K (p.61 c.181C>A) | 0.07% | 0.20% | 4.40% | 0.00% | 0.33% | 10.52% |

TABLE S11-1. Comparison results for mBDA NGS inferred VAF and ddPCR VAF for three different mutations, using internally constructed reference samples.

Compared to the 80-plex mBDA SNP panel described earlier for cell line contamination detection, the melanoma mBDA panel has a higher (worse) inferred VAF for 0% VAF samples, at up to 0.07% VAF compared to 0.02%. Based on the inferred VAFs for nominally 0% VAF samples, we set our VAF limit of detection at roughly 0.2% VAF (3 times our highest observed VAF for 0% samples) for the melanoma mBDA NGS panel. We do not currently have a definitive explanation for why there appears to be a systematic difference in performance between SNP variant alleles vs. cancer mutations. Possibilities include potential epigenetic differences between coding genomics regions (cancer mutations) and noncoding regions (where most SNP loci occur) in the human cell line gDNA.

**Clinical Tumor Tissue Sample Analysis with ddPCR.** Given the single-plex nature of ddPCR and the limited quantity of clinical tissue samples that were available, we performed ddPCR comparison experiments only on a limited number of clinical tumor tissue samples for the NRAS p.G12C and NRAS p.Q61K mutations. The comparison of the inferred VAFs from ddPCR and from mBDA NGS are shown in Table S11-2 and S11-3. We note that the observed discordance between ddPCR and mBDA NGS is in line with expectations given the low number of variant molecules and stochasticity of Poisson distribution.

The notable exception is the NRAS-Q61K c.181C>A mutation VAF for sample FF5, with ddPCR reporting 51% and mBDA NGS reporting 1.21%. Upon careful analysis of the mBDA NGS results, we note that FF5 is reported to have another NRAS mutation, p. Q61K c.180_181delinsTA, at 56% VAF. Thus, the ddPCR results can be thought of either as a false positive if considering only specifically the cDNA mutation, or as a concordance with mBDA NGS if considering all NRAS p.Q61K mutations. The lower FAM fluorescence values of the Variant positive droplets for the FF5 sample indicates however, that this was a "lucky" occurrence for ddPCR, and that other Q61K mutations with different DNA substitutions may not all be detected with equal sensitivity/efficacy.

| Sample | ddPCR VAF | mBDA NGS VAF |
|---|---|---|
| FF-1 | 0.00% | 0.50% |
| FF-4 | 0.04% | 0.00% |
| FF-5 | 0.06% | 0.00% |
| FF-6 | 0.00% | 0.08% |
| FF-10 | 0.05% | 0.26% |
| FF-13 | 0.08% | 0.00% |
| FF-18 | 0.11% | 0.01% |

TABLE S11-2. Comparison the clinical samples' VAF values of ddPCR and mBDA NGS for mutation NRAS-G12S (p.12 c.34G>A).

| Sample | ddPCR VAF | mBDA NGS VAF |
|---|---|---|
| FF-1 | 0.00% | 0.27% |
| FF-4 | 0.00% | 0.05% |
| FF-5 | 51.00% | 1.21% |
| FF-6 | 0.00% | 0.21% |
| FF-10 | 0.00% | 0.22% |
| FF-13 | 0.06% | 0.03% |
| FF-18 | 0.00% | 0.04% |

TABLE S11-3. Comparison the clinical samples' VAF values of ddPCR and mBDA NGS for mutation NRAS-Q61K (p.61 c.181C>A).

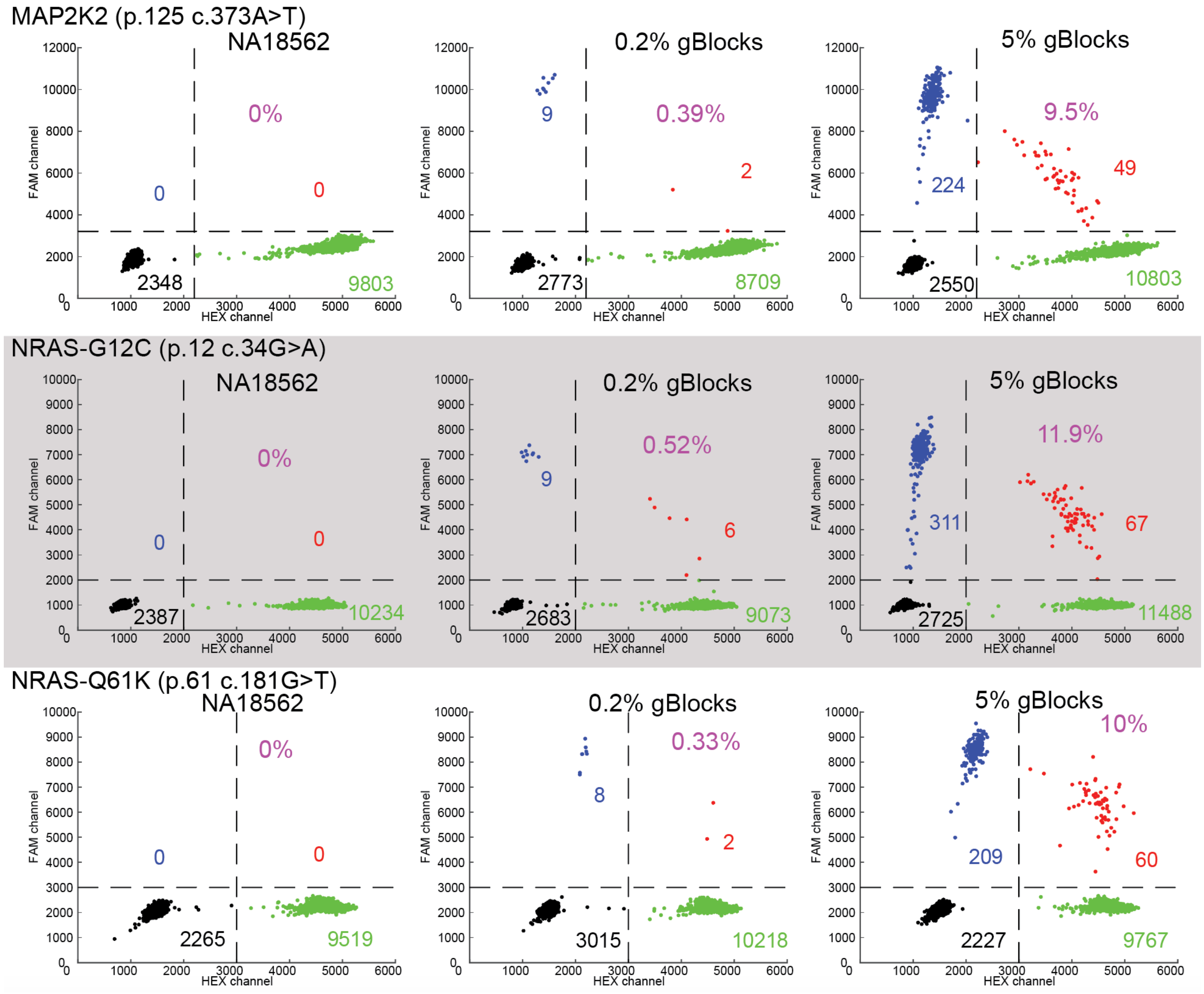


FIG. S11-1. ddPCR results on internal reference samples. Numbers shown are number of droplets in each quadrant. Inferred VAF is calculated via VAF $= \frac{TL+TR}{(TL+TR)+(TR+BR)}$, where $TL$, $TR$, and $BR$ denote the number of top left, top right, and bottom right droplets, respectively.

### NRAS-G12S (p.12 c.34G>A)

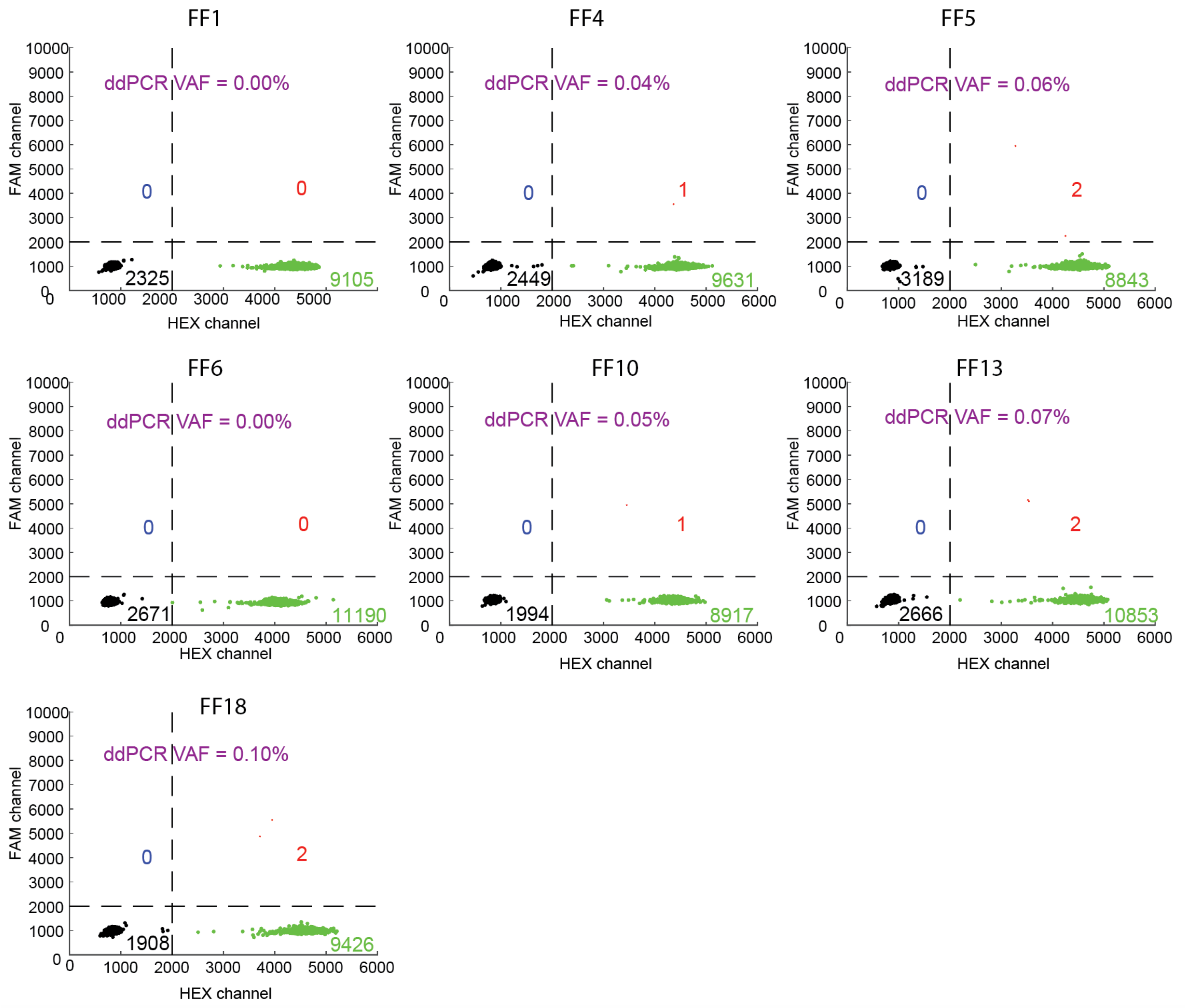


FIG. S11-2. ddPCR results on clinical samples for the NRAS-G12S (p.12 c.34G>A) mutation. Numbers shown are number of droplets in each quadrant. Inferred VAF is calculated via VAF $= \frac{TL+TR}{(TL+TR)+(TR+BR)}$, where $TL$, $TR$, and $BR$ denote the number of top left, top right, and bottom right droplets, respectively.

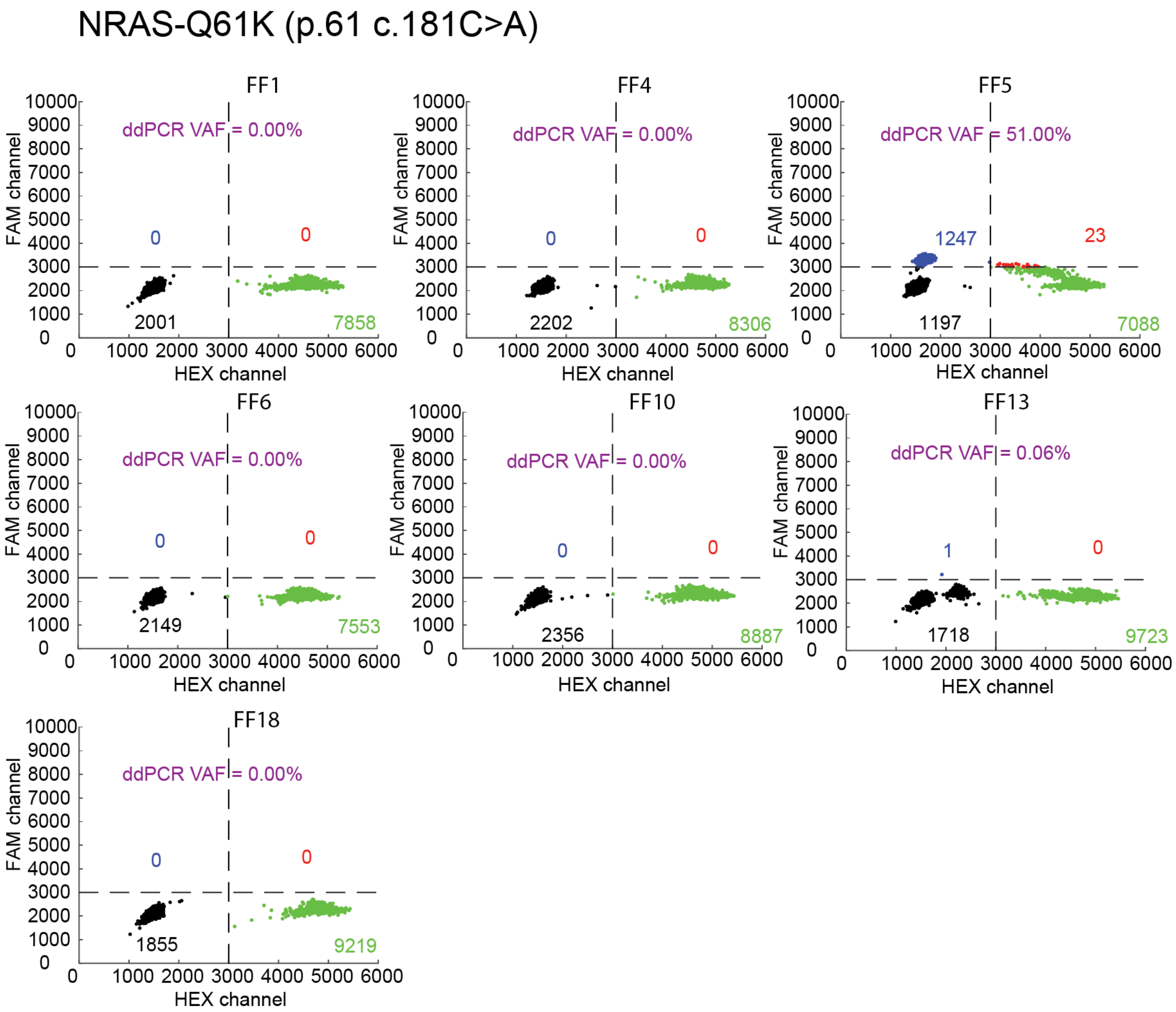


FIG. S11-3. ddPCR results on clinical samples for the NRAS-Q61K (p.61 c.181C>A) mutation. Numbers shown are number of droplets in each quadrant. Inferred VAF is calculated via VAF = $\frac{TL+TR}{(TL+TR)+(TR+BR)}$, where $TL$, $TR$, and $BR$ denote the number of top left, top right, and bottom right droplets, respectively. The sample FF5 was reported by mBDA NGS to have the p.Q61K c.180_181delinsTA mutation at 56% VAF, explaining the lower fluorescence positive droplets in the Top Left quadrant.

**Comparison of Clinical FFPE Tumor Tissue with ddPCR.** To further demonstrate the concordance between mBDA NGS VAFs and ddPCR VAFs, we performed comparison experiments for 4 formalin-fixed, paraffin-embedded (FFPE) tissue samples. Given the limited sample quantity, the fact that ddPCR experiments are single-plex, and the fact that there are limited numbers of validated ddPCR kits, we chose to perform comparison experiments only on the BRAF-V600E and NRAS-Q61K loci (Fig. S11-4 and Fig. S11-5). In all 8 experiments, we achieved qualitative concordance between mBDA and ddPCR. For NRAS-Q61K and Sample 1, the ddPCR VAF was lower than the mBDA VAF (0.03% vs. 0.30%); this could be due to Poisson error of small numbers of molecules.

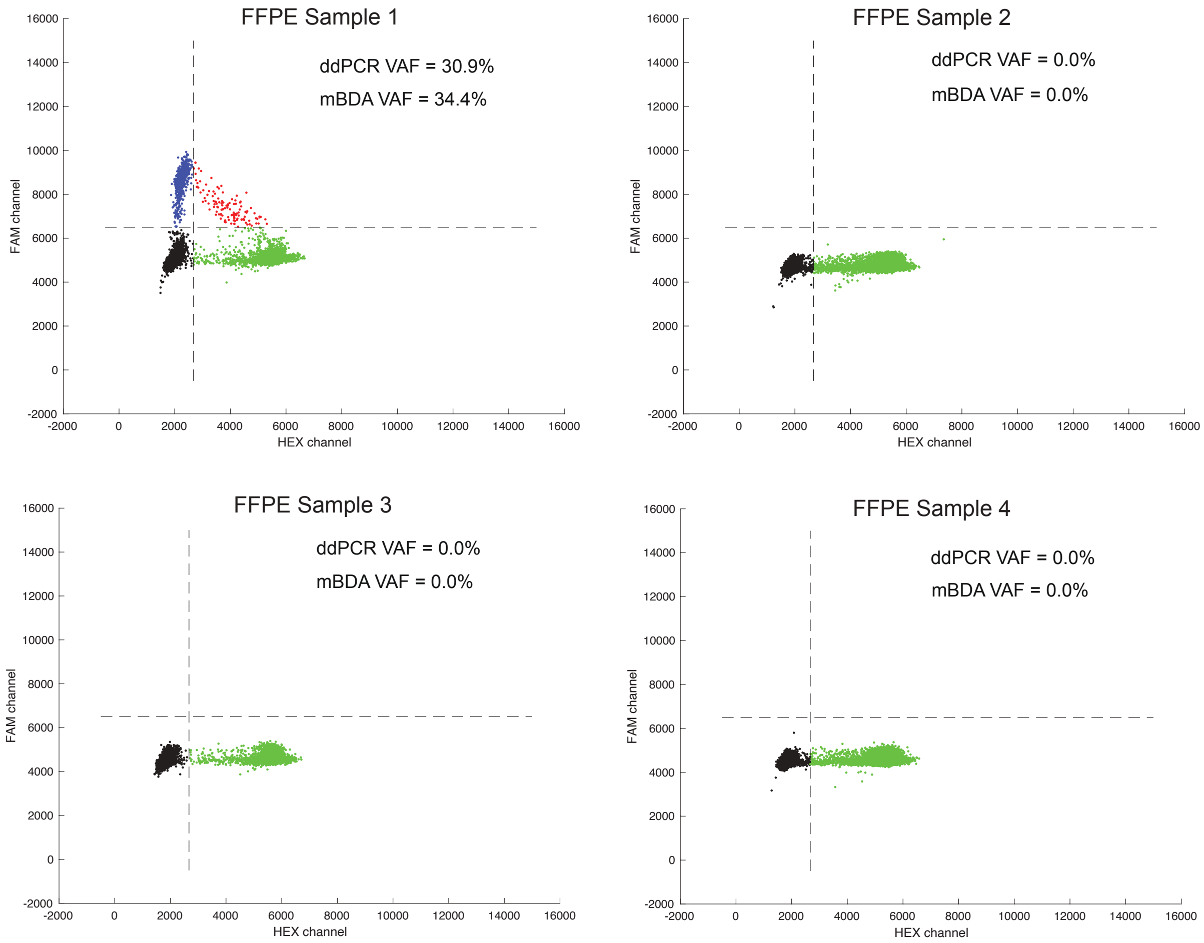


FIG. S11-4. ddPCR results and mBDA results on 4 melanoma FFPE tissue samples on the BRAF-V600E locus. Sample 1 was called positive for BRAF-V600E by both ddPCR and mBDA, and both methods estimated similar VAFs between 30% and 35%.

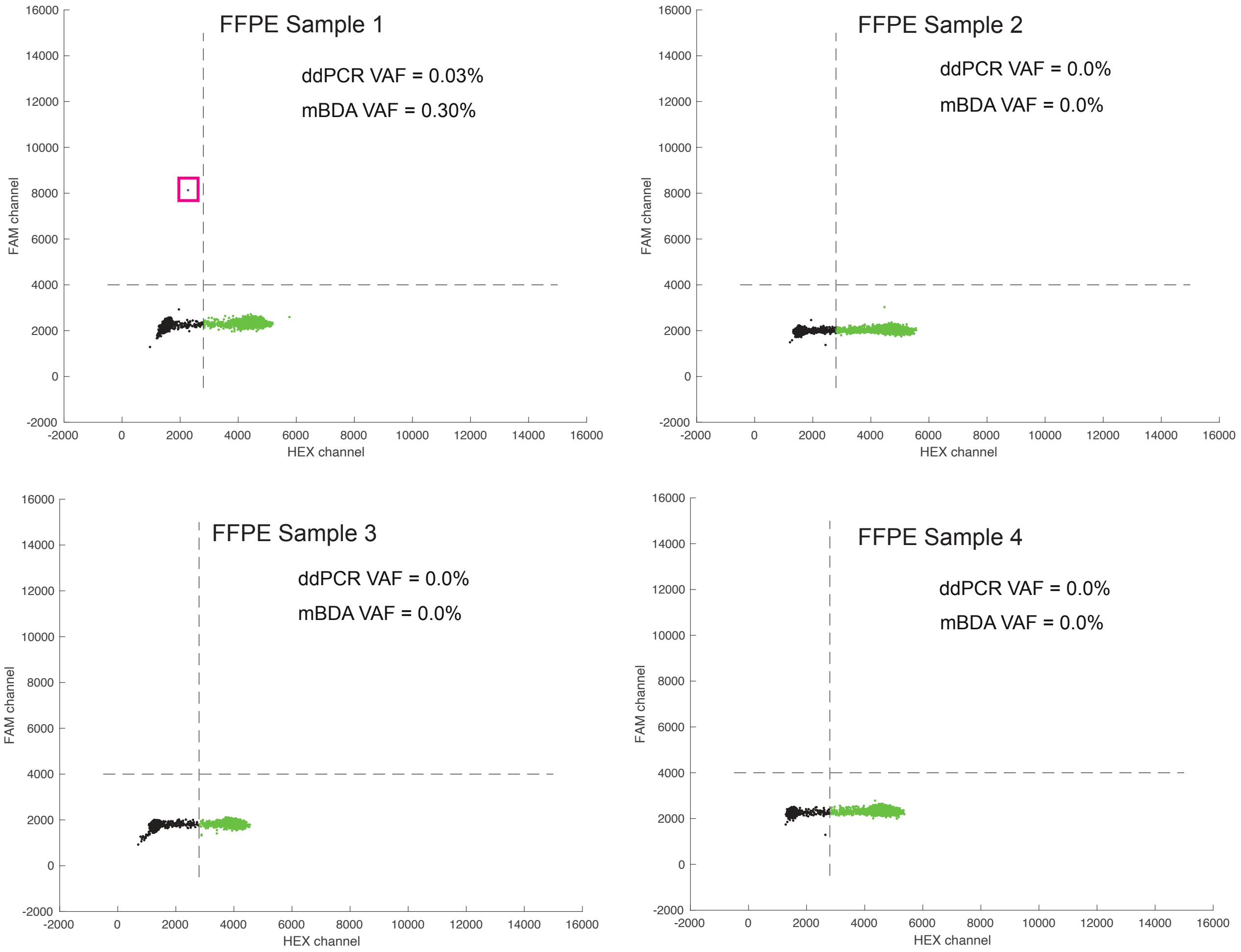


FIG. S11-5. ddPCR results and mBDA results on 4 melanoma FFPE tissue samples on the NRAS-Q61K locus. Sample 1 was called positive for NRAS-Q61K by both ddPCR and mBDA. The ddPCR VAF was lower than the mBDA VAF (0.03% vs. 0.30%); this could be due to Poisson error of small numbers of molecules.
